# How development affects evolution

**DOI:** 10.1101/2021.10.20.464947

**Authors:** Mauricio González-Forero

## Abstract

Natural selection acts on developmentally constructed phenotypes, but how does development affect evolution? This question calls for simultaneous consideration of development and evolution. However, there has been a lack of general mathematical frameworks mechanistically integrating the two, which may have inhibited progress on the question. Here we use a new mathematical framework that mechanistically integrates development into evolution to analyse how development affects evolution. We show that, whilst selection pushes genotypic and phenotypic evolution up the fitness landscape, development determines the admissible evolutionary pathway, such that evolutionary outcomes occur at path peaks rather than landscape peaks. Changes in development can generate path peaks, triggering genotypic or phenotypic diversification, even on constant, single-peak landscapes. Phenotypic plasticity, niche construction, extra-genetic inheritance, and developmental bias alter the evolutionary path and hence the outcome. Thus, extra-genetic inheritance can have permanent evolutionary effects by changing the developmental constraints, even if extra-genetically acquired elements are not transmitted to future generations. Selective development, whereby phenotype construction points in the adaptive direction, may induce adaptive or maladaptive evolution depending on the developmental constraints. Moreover, developmental propagation of phenotypic effects over age enables the evolution of negative senescence. Overall, we find that development plays a major evolutionary role.

## Introduction

Development may be defined as the process that constructs the phenotype across the lifespan. Natural selection must then act upon what development constructs, raising the question of how development affects evolution. Addressing this question might benefit from an integrated consideration of development and evolution, attending to both the mechanisms of evolution and the mechanisms of development. Efforts to address this question have increasingly invoked a need for a synthesis analogous to the modern synthesis between natural selection and genetics of the early twentieth century (Goldschmidt, 1940; Waddington, 1957; Alberch *et al*., 1979; Gould and Lewontin, 1979; Gould, 1980; West-Eberhard, 2003; Pigliucci, 2007; Pigliucci and Müller, 2010; Laland *et al*., 2014, 2015). Key early steps in the modern synthesis involved the formulation of general mathematical frameworks that integrated the mechanisms of natural selection and inheritance, namely, population and quantitative genetics. Similarly, general mathematical frameworks integrating the mechanisms of development into evolution might help advance our understanding of the evolutionary effects of development. Large bodies of empirical and theoretical research, as well as entire scientific journals and scientific societies have been increasingly devoted to establish the effects of development on evolution from different angles (Schlosser and Wagner, 2004; Wagner, 2005; Müller, 2007; Carroll, 2008). However, despite steady growth in research efforts on this topic, there remains a lack of general mathematical frameworks synthesizing the mechanisms of development and evolution.

One of the available frameworks, namely quantitative genetics, provides some of the still fundamental understanding of how development affects evolution. Stemming from Fisher (1918), quantitative genetics describes development by relating genotype to phenotype in terms of regression coefficients (i.e., Fisher’s additive effects of allelic substitution). From this basis, quantitative genetics established a basic principle of adaptation for phenotypic evolution as the climbing of an adaptive topography (Lande, 1979). According to this principle, under certain assumptions, evolution by natural selection can be seen as the climbing of a fitness landscape over phenotype space, where selection pushes evolution in the direction of steepest ascent in the landscape and genetic covariation may divert evolution in a less steep direction. This principle indicates that development affects evolution by shaping the genetic covariation upon which selection acts, where this covariation is quantified by the **G** matrix (Lande, 1979; Charlesworth *et al*., 1982; Cheverud, 1984; Maynard Smith *et al*., 1985; Klingenberg, 2010). This perspective gives development two fundamental possibilities. First, if development yields genetic variation in all directions of phenotype space (such that **G** is non-singular), then development may affect the direction of evolution but evolutionary outcomes still occur at peaks of the fitness landscape. Second, if development yields no genetic variation in some directions of phenotype space (such that **G** is singular), then evolution can stop away from fitness landscape peaks in which case development not only affects the direction of evolution but may also affect where the evolutionary outcomes are in the fitness landscape (Houle, 1991; Kirkpatrick and Lofsvold, 1992; Altenberg, 1995). The first possibility gives development a relatively minor role in evolution, as in such case selection alone pre-defines the evolutionary outcomes (i.e., landscape peaks) and development can at most only influence which outcome is achieved. Instead, the second possibility might give development a relatively major evolutionary role, as selection and development would jointly define the evolutionary outcomes (i.e., away from landscape peaks).

Progress on establishing which of the two possibilities is the case has been difficult. Despite substantial research effort, the answer has remained uncertain, in part due to the difficulties of empirically establishing whether **G** is singular (Kirkpatrick, 2009). More fundamentally, one reason for this uncertainty may have been that equations describing phenotypic evolutionary change have been derived from a regression description of development, where trait values are regressed on gene content, yielding a regression-based understanding of genetic covariation (Fisher, 1918; Lande, 1979). Since regression describes relationships regardless of their underlying mechanisms, this approach to genetic covariation may have limited a mechanistic understanding of the nature of genetic covariation arising from development. In principle, a mechanistic integration of development into evolution could yield a mechanistic understanding of genetic covariation and of the **G**-matrix.

The task of devising a general mathematical framework that mechanistically integrates development into evolution would involve many complexities, which has likely contributed to the persistent lack of such a framework. Calls for synthesis between developmental and evolutionary biology have asked for consideration of the mechanistic basis of phenotype construction, non-linear genotype-phenotype maps, gene-gene and gene-environment interactions, non-normal distributions, far-from-equilibrium evolutionary dynamics, dynamic fitness landscapes, evolution and the nature of the **G**-matrix, evolvability and epigenetics, and a variety of other complexities (Pigliucci and Schlichting, 1997; Pigliucci, 2007). Calls for integration of development into evolution have also highlighted several developmental factors — namely phenotypic plasticity (West-Eberhard, 2003), niche construction (Odling-Smee *et al*., 1996), extra-genetic inheritance (Jablonka and Lamb, 2014), and developmental bias (Arthur, 2004) — as possibly having important, yet unrecognised, evolutionary consequences (Laland *et al*., 2014, 2015). Mathematical frameworks and specific mathematical models integrating some of these complexities have become available in recent decades (e.g., Dieckmann and Law, 1996; Caswell, 2001; Hansen and Wagner, 2001; Day and Bonduriansky, 2011; Mullon and Lehmann, 2018; Lande, 2019; Chantepie and Chevin, 2020; Engen and Sæther, 2021). Additionally, theoretical research has often used individual-based simulations integrating some these complexities (e.g., Salazar-Ciudad and Marín-Riera, 2013; Watson *et al*., 2013; Jones *et al*., 2014a; Miloco and Salazar-Ciudad, 2022). However, it is of particular interest to obtain general mathematical frameworks mechanistically integrating development into evolution to seek deeper insight — general in the sense of encompassing a broad class of models. Despite the progress made, a synthetic mathematical framework unifying these complexities in a general and tractable manner had remained unavailable until recently.

A new mathematical framework (González-Forero, 2021) integrates mechanistic development into evolution while incorporating the elements listed in the previous paragraph by building upon many advances in evolutionary modelling over the last decades (Dieckmann and Law, 1996; Caswell, 2001) (and references in González-Forero 2021). This framework integrates conceptual and mathematical advances from adaptive dynamics (Dieckmann and Law, 1996), matrix population models (Caswell, 2001), and optimal control theory as used in life-history theory (e.g., Schaffer, 1983; Sydsæter *et al*., 2008). The framework yields formulas that relate mechanistic descriptions of development to genetic covariation and to plastic change separate from selection. Here we use this framework to analyse how development affects evolution. This analysis sharpens the principle of adaptation as the climbing of a fitness landscape and provides insights into a wide array of long-standing questions. We focus on developing conceptual understanding and refer the reader to González-Forero (2021) for technical details.

## Framework overview

The framework is based on standard assumptions of adaptive dynamics (Dieckmann and Law, 1996; Metz *et al*., 1996). It considers a resident (i.e., wild-type) population where individuals can be of different ages (i.e., it is age-structured), reproduction is clonal for simplicity, whereby offspring receive the same genotype of their parent, and individuals can interact socially but only with non-relatives for simplicity. The genetic architecture (e.g., number of loci, ploidy, or linkage) need not be specified given our assumption of clonal reproduction, but it may be specified in particular models. Each individual has three types of age-specific traits that we let take continuous values to take derivatives. First, individuals have genotypic traits, which we refer to as the genotype and which are directly specified by genes. For instance, genotypic traits may be a continuous representation of nucleotide sequence or may be life-history traits assumed to be under direct genetic control. Genotypic traits correspond to control variables in the terminology of optimal control theory. Second, individuals have phenotypic traits, which we refer to as the phenotype and which are constructed over life depending on the genotype, developmental history, environment, social interactions, and their interaction, and where such phenotype construction is subject to developmental constraints. For instance, phenotypic traits include morphology and behavior. Phenotypic traits correspond to state variables in the terminology of optimal control theory. Third, individuals have environmental traits, which we refer to as the environment and which describe the individual’s local environment, possibly modified by the individual, social partners, sources exogenous to the population, and their interaction, and where such environmental alteration is subject to environmental constraints. For instance, environmental traits include ambient temperature or humidity, which the individual may adjust, such as by roosting in the shade. This terminology contrasts with standard adaptive dynamics terminology, which would call genotypic traits phenotypes while here the phenotype is only traits that are developed; also standard adaptive dynamics terminology would define the environment as global, including everything outside the individual while here the environment is only local to the individual to model niche construction. Once the resident population achieves carrying capacity, rare mutant individuals arise who have a marginally different genotype from the resident genotype, drawn from an unbiased distribution of the deviation of mutant genotypes from the resident. Thus, we assume mutation is rare, weak, and unbiased. Population dynamics is deterministic so the only source of stochasticity is mutation. We assume that if rare mutants increase in frequency, they achieve fixation, which establishes a new resident, thus yielding evolutionary change.

The framework uses the following notation. The genotype across all genotypic traits for a mutant individual of age *a* ∈ {1,…, *N*_a_} is described by the vector 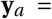 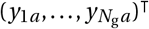 of *N*_g_ genotypic traits (control variables; e.g., *y_ia_* may be a continuous variable representing — via a differentiable approximation of the Heaviside step function — whether nucleotide *I* = ⌈*i* /*n*⌉ is present at locus *J* = *i* − ⌊*i* /*n*⌋*n* at age *a* for *n* loci, in which case *N*_g_ = 4*n* with four nucleotides). While the genotype of an individual may often be constant with age, we let it depend on age as genotypic traits vary with age in life-history models. The phenotype of a mutant individual of age *a* is described by the vector 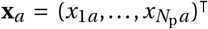 of *N*_p_ phenotypic traits (state variables; e.g., *x_ia_* may be the size of trait *i* at age *a*). The environment of a mutant individual of age *a* is described by the vector 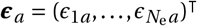 of *N*_e_ mutually independent environmental traits (e.g., *ϵ*_*ia*_ may be the temperature experienced at age *a*). A mutant individual’s phenotype, genotype, and environment across all ages are described respectively by the block-column vectors 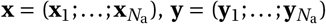, and 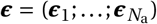 (the semicolon indicates a linebreak). The framework finds that two aggregates of traits have special importance. First, we aggregate genotype and phenotype into a single vector called the “geno-phenotype” (Feldman and Zhivotovsky 1992 use the term phenogenotype). Thus, the geno-phenotype of a mutant individual of age *a* is described by **z**_*a*_ = (**x**_*a*_; **y**_*a*_). The mutant geno-phenotype across all ages is **z** = (**x**; **y**). Second, we similarly aggregate genotype, phenotype, and environment into a single vector called the “geno-envo-phenotype”. The geno-envo-phenotype of a mutant individual of age *a* is described by the block-column vector **m**_*a*_ = (**z**_*a*_; ***ϵ**_a_*). The mutant geno-envo-phenotype across all ages is **m** = (**z**; ***ϵ***). Resident variables are denoted analogously with an over-bar (e.g., 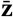). We assume that the mutant genotype **y** is a random variable, whose deviation from the resident is given by the mutational distribution *M* (**y**− **ȳ**) with mean zero and vanishingly small and unbiased variance (i.e., 0 < tr(cov[**y**, **y**]) ≪ 1 and *M* is even).

To mechanistically incorporate development, we describe an individual’s phenotype at a given age as a function of her genotype, phenotype, and environment at the immediately preceding age and of the social interactions experienced at that age. Thus, the phenotype of a mutant individual at age *a* + 1 satisfies the developmental constraint

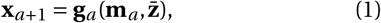

where the developmental map 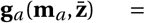 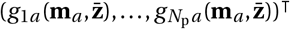 depends on the mutant’s genotype, phenotype, and environment at age *a* and on the genotype and phenotype of social partners of any age. For simplicity, we assume that the developmentally initial phenotype **x**_1_ is constant. We also assume that the genotype **y** is developmentally independent, that is, each *y_ia_* is entirely determined by the relevant genotypic trait (e.g., nucleotide presence at the relevant locus) but is not influenced by other genotypic traits (e.g., nucleotide presence at other loci), the phenotype, or environment (i.e., in the terminology of optimal control theory, we assume open-loop control; Sydsæter *et al*. 2008). Eq. (1) is a constraint in that the phenotype **x** cannot take arbitrary values but only those satisfying that equation. The developmental map can evolve and take any (differentiable) form as the genotype, phenotype, and environment evolve. The developmental map can also change over development (i.e., the functions **g**_*a*_ may be different to **g**_*j*_ for *j* ≠ *a*), for example, with metamorphosis.

To mechanistically incorporate niche construction and plasticity, we describe an individual’s environment as a function of the genotype and phenotype of the individual or social partners, and of processes exogenous to the population. The dependence of the environment on the genotype and phenotype of herself and social partners can describe niche construction by her or her social partners. The dependence of the environment on exogenous processes can describe, for instance, eutrophication or climate change caused by members of other species. For simplicity, we assume that exogenous environmental change is slow to allow the population to achieve the carrying capacity. Thus, a mutant’s environment at age *a* satisfies the environmental constraint

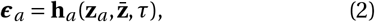

where the environmental map 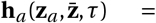 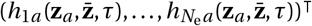 depends on the mutant’s genotype and phenotype at that age (e.g., from behavioural choice or environmental modification), on her social partners’ genotypes and phenotypes at any age (e.g., from social niche construction), and on the evolutionary time *τ* due to exogenous environmental change. The environment can then change over development (i.e., ***ϵ***_*a*_ may be different from ***ϵ***_*j*_ for *a* ≠ *j*) and evolution (either as the population evolves or due to exogenous causes). Eq. (2) is also a constraint in that the environment ***ϵ*** can only take values allowed by that equation. Although social interactions are part of the environment (***ϵ***_a_ depends on 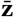), the developmental map (1) may also directly depend on social interactions to allow for modelling social interactions without having to also consider environmental traits. A mutant’s fertility 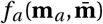 and survival probability 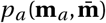 at age *a* depend on her genotype, phenotype, and environment at that age, and on the genotype, phenotype, and environment of her social partners of any age.

The developmental constraint (1) incorporates: developmental bias, in that the phenotype **x** may be predisposed to develop in certain ways; phenotypic plasticity, in that the same genotype **y** can generate a different phenotype **x** under a different environment ***ϵ***; adaptive phenotypic plasticity, for example, via somatic selection or reinforcement learning (e.g., if *g_ia_* is proportional to the gradient 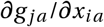 of a material payoff *g_ja_* earned at age *a* that increases fitness); niche construction, in that environmental traits depend on developmentally constructed traits of the individual and social partners; and extra-genetic inheritance, in that an individual’s phenotype at age *a* + 1 can be an identical or modified copy of non-relatives’ phenotypes (this includes Jablonka and Lamb’s (2014) notion that extra-genetic inheritance involves the copying or reconstruction of others’ phenotype rather than their genotype). Thus, the developmental constraint (1) formalizes the verbal definition of developmental constraints “as biases on the production of variant phenotypes or limitations on phenotypic variability caused by the structure, character, composition, or dynamics of the developmental system” (Maynard Smith *et al*., 1985). Moreover, the developmental constraint (1) can evolve as explained above, be non-linear, and mechanistically describe development, gene-gene interaction, and gene-environment interaction. To explore suggestions of adaptation without selection (e.g., Laland *et al*., 2015), we allow for the possibility that the developmental map **g**_*a*_ depends on the selection gradient 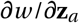 (defined below), in which case we say that development is selective whereby development has information about the adaptive direction. Allowing for selective development does not violate the assumptions of the developmental constraint (1), even though such perfectly idealised selective development may be biologically infeasible. Mechanistically, selective development might occur to some extent with individual or social learning, or with somatic selection (West-Eberhard, 2003).

The framework describes the evolutionary developmental (evo-devo) dynamics as follows. At each evolutionary time *τ*, the developmental dynamics of the resident phenotype are given by the developmental constraint (1) evaluated at the resident genotype. In turn, the evolutionary dynamics of the resident genotype are given by the canonical equation of adaptive dynamics (Dieckmann and Law, 1996; Dieckmann *et al*., 2006; Durinx *et al*., 2008)

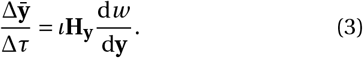

Throughout, derivatives are evaluated at **y** = **ȳ** and are in matrix calculus notation (Supplementary Information section S1; we use an equal sign in Eq. (3) and evolutionarily dynamic equations below but they are strictly first-order approximations). The evo-devo dynamics are thus given by the evolutionary dynamics of the genotype (Eq. 3) and the concomitant developmental dynamics of the phenotype (Eq. 1) (a juxtaposition already used by- Parvinen *et al*. 2013 and Metz *et al*. 2016). The right-hand side of Eq. (3) has three components. First, *ι* is a non-negative scalar measuring mutational input, which is proportional to the mutation rate and the carrying capacity. Second, **H_y_** = cov[**y**, **y**] is the mutational covariance matrix (H for heredity). Third, d*w* /d**y** is the total selection gradient of the genotype, which measures total genotypic selection, that is, total directional selection on genes considering the ability of genes to affect the phenotype (from the chain rule, the total derivative of a function *f* (*x*, *y*) with respect to *y*, where *x* = *g* (*y*), is d *f* /d*y* = (*∂f* /*∂x*)(d*x*/d*y*) + *∂f* /*∂y*). In contrast, *∂w* /*∂***y** measures direct directional selection on genes without considering such ability (*∂w* /*∂***y** is traditionally assumed zero, but is not zero in standard life-history models where the genotype modulates resource allocation to fertility and is thus under direct selection; González-Forero, 2021). Thus, *∂w* /*∂***y** is a vector that points in the direction of steepest increase in fitness in genotype space without constraints, whereas d*w* /d**y** is a vector that points in the direction of steepest increase in fitness in genotype space subject to the developmental (1) and environmental (2) constraints. González-Forero (2021) finds closed-form formulas for the total selection gradient of the genotype. Because of age-structure, a mutant’s relative fitness is

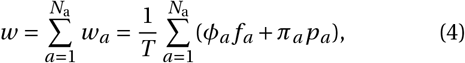

where *w_a_* is the mutant’s relative fitness at age *a*, *T* is generation time, and *ϕ_a_* and *π_a_* are the forces (Hamilton, 1966) of selection on fertility and survival at age *a* (given in Eqs. S5 in the Supplementary Information). The quantities *T*, *ϕ_a_*, and *π_a_* depend on resident but not mutant values. The total selection gradient of the genotype is a form of Caswell’s (1982, 2001) total derivative of fitness, Charlesworth’s (1994) total differential of the population’s growth rate, van Tienderen’s (1995) integrated sensitivity of the population’s growth rate, and Morrissey’s (2014, 2015) extended selection gradient.

## Results

### Recovery of classic but insufficient results

We investigate how development as described by Eq. (1) affects evolution by analysing dynamically sufficient equations describing the long-term evolution of a developed phenotype 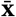 as the climbing of a fitness landscape. Before arriving at such equations, we first describe a mechanistic version of the Lande (1979) equation and explain why it is generally insufficient to describe long-term evolution of developed phenotypes.

Assume for now that there is (i) no niche construction (*∂**ϵ***^⊤^/*∂***z** = **0**), (ii) no social development 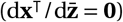 (iii) no exogenous environmental change (*∂**ϵ***/*∂τ* = **0**), and (iv) no direct genotypic selection (*∂w* /*∂***y** = **0**). Then, González-Forero (2021) shows that the (expected) evolutionary dynamics of the resident phenotype (as Δ*τ* → 0) satisfy

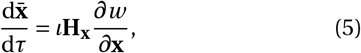

where **H_x_** guarantees that the developmental (1) and environmental (2) constraints are satisfied at all times. The right-hand side of Eq. (5) has two different components relative to those of Eq. (3). First, *∂w* /*∂***x** corresponds to Lande’s selection gradient, which measures direct directional selection on the phenotype: it is a vector that points in the direction of steepest increase in fitness in phenotype space without constraints. Second, Eq. (5) depends on the *mechanistic* additive genetic covariance matrix of the phenotype

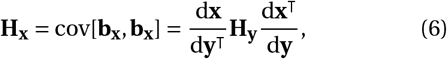

where the *mechanistic* breeding value of **x** is defined as

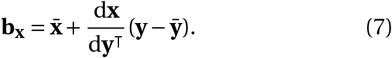

The key difference of Eq. (7) with breeding value is that the latter is defined using a matrix of regression coefficients in place of d**x**/d**y**^⊤^ (Lynch and Walsh, 1998; Walsh and Lynch, 2018). Such regression coefficients are Fisher’s (1918) additive effects of allelic substitution and the matrix formed by them is Wagner’s (1984) developmental matrix. Whereas such regression coefficients are obtained via least squares, the matrix d**x**^⊤^/d**y** is obtained from the mechanistic description of development in Eqs. (1) and (2). Thus, the matrix d**x**^⊤^/d**y** is a mechanistic counterpart of both Fisher’s (1918) additive effects of allelic substitution and Wagner’s (1984) developmental matrix. Consequently, **H_x_** is a mechanistic counterpart of Lande’s (1979) **G**-matrix. González-Forero (2021) obtained closed-form formulas for d**x**^⊤^/d**y** from the developmental (1) and environmental (2) constraints, which allow for a mechanistic understanding of genetic covariation. The fact that mechanistic breeding value is obtained from d**x**^⊤^/d**y** rather than regression coefficients means that breeding value and mechanistic breeding value have different properties. In particular, while breeding value is uncorrelated with the residual of predicting the phenotype because of least squares, mechanistic breeding value can be correlated with the residual (i.e., for a single developed trait *x* = *b_x_* + *e_x_* with residual *e_x_*, but cov[*x*, *e_x_*] ≠ 0). Consequently, the classic partition of phenotypic variance into additive genetic and “environmental” variances does not hold with mechanistic breeding value. Hence, narrow-sense mechanistic heritability of a single developed trait *x* defined with the traditional formula var[*b_x_*]/var[*x*] can be greater than one.

Eq. (5) may then be understood as a mechanistic Lande (1979) equation, apart from some differences due to our adaptive dynamics assumptions but these differences do not affect the points we make. Indeed, the expression for relative fitness (4) has the form of that in the Lande (1982) equation for quantitative genetics under age structure. The expression for the mechanistic additive genetic covariance matrix of the phenotype (6) has the same form of previous expressions obtained under quantitative genetics assumptions, which in place of our closed-form derivatives had regression coefficients or derivatives of unknown form (see Eq. II of Fisher 1918, Eq. + of Wagner 1984, Eq. 3.5b of Barton and Turelli 1987, and Eq. 4.23b of Lynch and Walsh 1998; see also Eq. 22a of Lande 1980, Eq. 3 of Wagner 1989, and Eq. 9 of Charlesworth 1990). While in quantitative genetics the matrix corresponding to **H_y_** would describe the realized standing variation in gene content, here **H_y_** describes the expected variation in genotype due only to mutation in the current evolutionary time and so we call it the mutational covariance matrix. This different meaning of **H_y_** in the two frameworks does not mean that **H_y_** in our framework induces absolute constraints that are absent under quantitative genetics assumptions: our definition of **H_y_** allows for mutational variation in all directions of genotype space, or only in some directions (i.e., **H_y_** may be full rank, or less than full rank), as in quantitative genetics. While Lande’s (1979) **G** matrix is the covariance matrix of breeding values of the phenotypes, **H_x_** is the covariance matrix of the mechanistic breeding values of the phenotypes. Mechanistic breeding value (7) is the first-order estimate of the phenotype with respect to genotype as predictor, which corresponds to the quantitative genetics notion of breeding value as the best linear estimate of the phenotype with respect to gene content as predictor (Falconer and Mackay, 1996; Lynch and Walsh, 1998). Hence, genotypic traits play here an analogous role to that of gene content in quantitative genetics in that they emerge as the relevant first-order predictor of the phenotype to describe inheritance.

In contrast to the Lande (1979) equation, the mechanistic Lande Eq. (5) has been derived from a mechanistic account of development so we have formulas to relate development (1) to mechanistic breeding value and so to mechanistic genetic covariation **H_x_**. These formulas guarantee that the developmental (1) and environmental (2) constraints are satisfied at all times. The formulas show that 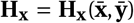 is generally a function of the resident genotype **ȳ**, in particular, if development is non-linear, in which case the developmental matrix d**x**^⊤^/d**y** depends on **ȳ**. Specifically, gene-gene interaction (products between *y*’s in the developmental map), gene-phenotype interaction (products between *y*’s and *x*’s), and gene-environment interaction (products between *y*’s and *ϵ*’s) cause the resident genotype **ȳ** to remain in d**x**^⊤^/d**y**. Thus, in general, the right-hand side of the mechanistic Lande Eq. (5) depends on the resident genotype **ȳ** if the mutation rate, carrying capacity, mutational covariation **H_y_**, the developmental matrix d**x**^⊤^/d**y**, or direct phenotypic selection *∂w* /*∂***x** depend on the resident genotype **ȳ**. Consequently, Eq. (5) is under-determined as it generally depends on the resident genotype but does not describe the evolutionary dynamics of the resident genotype. Hence, Eq. (5) cannot be used alone to describe the evolutionary dynamics of the resident phenotype in general.

This problem equally applies to the Lande equation. Since breeding value depends on the regression coefficients of phenotype on gene content at the current time, such regression coefficients depend on the current state of the population, including allele frequency. Thus, the Lande equation depends on allele frequency but it does not describe allele frequency change. The standard approach to address this problem is to assume Fisher’s infinitesimal model, whereby each phenotype is controlled by an arbitrarily large number of loci so in the short term allele frequency change per locus is negligible (Barton *et al*., 2017; Hill, 2017; Walsh and Lynch, 2018). Then, **G** is assumed constant or its evolution is described by the Bulmer equation, which considers change in **G** due to change in linkage disequilibrium while still assuming negligible allele frequency change (Lande and Arnold, 1983; Turelli, 1988; Barton *et al*., 2017). Such an approach allows for the Lande equation to describe evolution only in the short term, where allele frequency change per locus remains negligible (Barton *et al*., 2017). However, longterm evolution involves non-negligible allele frequency change, thus limiting the ability of the Lande equation to describe longterm phenotypic evolution.

In turn, the canonical equation of adaptive dynamics (Dieckmann and Law, 1996) describes longterm phenotype evolution, but where the phenotype does not have developmental constraints. In our terminology, the canonical equation of adaptive dynamics describes the long-term evolution of genotypic traits, which do not have developmental constraints (Eq. 3; Dieckmann and Law, 1996). Thus, the canonical equation does not describe the long-term evolution of developed traits as an adaptive topography. The canonical equation for function-valued traits (Dieckmann *et al*., 2006) describes the evolution of genotypic traits that can affect the construction of phenotypic traits (Parvinen *et al*., 2013), but this equation still describes evolution in gradient form only for genotypic traits, not phenotypic traits (i.e., not for state variables; indeed, given the age-block-structure of **y**, Eq. (3) may be understood as the canonical equation for function-valued traits in discrete age). Equations describing the long-term evolution in gradient form of phenotypic traits (i.e., with explicit developmental constraints, that is, of state variables) such as Eq. (5), where formulas for **H_x_** guarantee that the developmental constraints are met, do not seem to have been available until recently (González-Forero, 2021). Yet, the dynamic insufficiency of Eq. (5) limits its use to analyse how the evolution of developed traits proceeds in the fitness landscape. (For a different view of how the canonical and Lande equations are related see pp. 1084-1086 of Geritz *et al*. 2016).

### Development blocks evolutionary change

Even though the mechanistic Lande Eq. (5) is insufficient to describe the long-term evolution of developed phenotypes particularly if development is non-linear, such equation can be extended to allow for this.

To see this, remove the assumption (iv) above, and still assume that there is (i) no niche construction (*∂**ϵ***^⊤^/*∂***z** = **0**), (ii) no social development 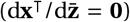, and (iii) no exogenous environmental change (*∂**ϵ***/*∂τ* = **0**). Then, González-Forero (2021) shows that the evolutionary dynamics of the resident phenotype and genotype are given by

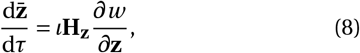

where **H_z_** guarantees that the developmental (1) and environmental (2) constraints are satisfied at all times. Eq. (8) now depends on the mechanistic additive genetic covariance matrix of the geno-phenotype

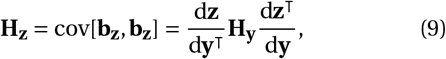

where the mechanistic breeding value **b_z_** of the geno-phenotype is given by Eq. (7) replacing **x** with **z**.

Eq. (8) is an extended mechanistic Lande equation that, in contrast to the Lande equation or its mechanistic counterpart (5), is dynamically sufficient and so describes the long-term evolution of developed traits as an adaptive topography. Eq. (8) is dynamically sufficient because it describes the dynamics of all the variables involved by simultaneously describing genotypic and phenotypic evolution since **z** = (**x**; **y**) includes both genotype and phenotype. In particular, the matrix 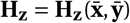, is still a function of the resident phenotype and genotype, but now the evolutionary dynamics of both are described by Eq. (8). The right-hand side of Eq. (8) may also be a function of the resident environmental traits, but because of assumptions (i) and (iii), niche construction and exogenous environmental change are absent so the environment remains constant.

As Eq. (8) describes the long-term evolution of developed traits as an adaptive topography, we can now use it to analyse the effect of development on genetic covariation and evolution. A biologically crucial property of Eq. (8) is that it implies that there are always absolute genetic constraints to adaptation within the assumptions made (absolute constraints mean that the constraining matrix is singular; Houle, 2001; Klingenberg, 2005, 2010). Indeed, Eq. (8) simultaneously describes genotypic and phenotypic evolution, so the climbing of the fitness landscape is in geno-phenotype space rather than only phenotype space as in the Lande equation. Yet, because the geno-phenotype **z** contains the genotype **y**, genetic covariation in geno-phenotype is absolutely constrained by the genotypic space (i.e., **H_z_** is always singular because d**z**^⊤^/d**y** has fewer rows than columns). That is, to achieve dynamic sufficiency, one cannot generally consider any traits for evolutionary analysis, but must consider both the phenotype and genotype, and since the phenotype and genotype are related by development, genetic covariation in geno-phenotype space is absolutely constrained. Thus, there cannot be genetic variation in as many directions in geno-phenotype space as there are phenotypes across life (i.e., **H_z_** has at least *N*_a_ *N*_p_ eigenvalues that are exactly zero). Along such directions evolutionary change is blocked (i.e., along the directions of the eigenvectors corresponding to the zero eigenvalues of **H_z_**). In this sense, development can be seen as blocking evolutionary change in some directions. Therefore, evolutionary stasis 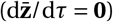 can occur away from landscape peaks, where direct directional selection persists (*∂w* /*∂***z** ≠ **0**; see also Houle 1991, Kirkpatrick and Lofsvold 1992, and Altenberg 1995). Thus, consideration of direct directional selection alone (*∂w* /*∂***z**) is not sufficient for predicting possible evolutionary outcomes, which depend also on **H_z_**. Predicting evolutionary outcomes still depends on **H_z_** even if there is no genotypic selection (i.e., if *∂w* /*∂***y** = **0**).

The singularity of **H_z_** and generalizations thereof has important implications, which we detail below. One immediate implication is that, without absolute mutational constraints (i.e., if **H_y_** is non-singular), evolutionary outcomes are jointly defined by direct selection and development as described by d**x**^⊤^/d**y**, which is influenced by gene-gene, gene-phenotype, and gene-environment interactions. Another immediate implication is that this singularity may help explain common empirical observations of evolutionary stasis in wild populations despite directional selection and genetic variation, termed the “paradox of stasis” (Merilä *et al*., 2001; Kirkpatrick, 2009; Kingsolver and Diamond, 2011). Because **H_z_** is singular, evolutionary stasis is expected to generally occur with persistent directional selection and genetic variation, although there are other explanations for the paradox of stasis, including measurement error (Merilä *et al*., 2001; Estes and Arnold, 2007; Haller and Hendry, 2013; Morrissey, 2015).

Even though plasticity may be present in the extended mechanistic Lande Eq. (8), plasticity has no evolutionary effect under the assumptions made in that equation. Indeed, the formulas for **H_z_** show that, under the framework’s assumptions, plasticity (i.e., *∂***z**^⊤^/*∂**ϵ***) only affects the evolutionary dynamics by interacting with niche construction or exogenous environmental change (see below). Thus, although in Eq. (8) there may be plasticity, it has no effect since by assumptions (i) and (iii) niche construction and exogenous environmental change are absent so the environment remains constant, but plasticity requires environmental change.

### Development determines the path

While development can be seen as blocking evolutionary change, development can be more specifically seen as determining the admissible evolutionary path. This is because one must follow the evolution of the genotype and phenotype for a dynamically sufficient phenotypic adaptive topography, and the phenotype is related to the genotype by the developmental constraint (1) along which evolution is allowed. Consequently, if there are no absolute mutational constraints and no exogenous environmental change, development and selection jointly define the evolutionary equilibria and development determines which of these equilibria are admissible.

To see this, continue to assume that there is (i) no niche construction (*∂**ϵ***^⊤^/*∂***z** = **0**), (ii) no social development 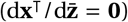, and (iii) no exogenous environmental change (*∂**ϵ***/*∂τ* = **0**). Then, the evolutionary dynamics of the resident phenotype and genotype are equivalently given by

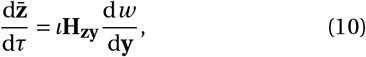

where **H_zy_** guarantees that the developmental (1) and environmental (2) constraints are satisfied at all times. This equation is closely related to Morrissey’s (2014) Eq. 4 but they differ as Morrissey’s equation is for the evolutionary change of the phenotype with linear development rather than geno-phenotype with possibly non-linear development, considers the total selection gradient of the phenotype rather than the genotype, considers regression-based rather than mechanistic genetic covariation (if the classic partition of phenotypic variance is to hold), and is dynamically insufficient as it is a rearrangement of the Lande (1979) equation. Eq. (10) depends on the total selection gradient of the genotype, which under assumptions (i-iii), is

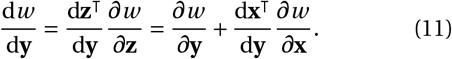

Thus, total genotypic selection depends on the developmental matrix d**x**^⊤^/d**y** (see Fig. S1 for a description of all the factors affecting total genotypic selection removing assumptions i-iii). Eq. (10) also depends on the mechanistic additive genetic cross-covariance matrix between the geno-phenotype and genotype

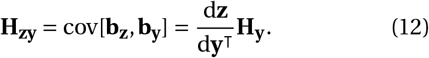

Crucially, the transpose matrix of the total effects of genotype on geno-phenotype, d**z**/d**y**^⊤^, is non-singular because the genotype is developmentally independent by assumption. Thus, Eq. (10) implies that evolutionary stasis occurs when total genotypic selection vanishes (d*w* /d**y** = **0**) if there are no absolute mutational constraints. That is, total genotypic selection can identify evolutionary equilibria, even though direct directional selection typically cannot. This is a different way of showing that evolutionary equilibria are generally defined jointly by direct selection and development if there are no absolute mutational constraints. Moreover, there are always an infinite number of evolutionary equilibria, because d*w* /d**y** = **0** provides fewer equations than those in 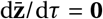. Since the developmental constraint (1) must be satisfied at all times, this provides the remaining equations and the admissible evolutionary trajectory. That is, the developmental constraints not only influence the evolutionary equilibria, but also provide the admissible evolutionary trajectory and so the admissible equilibria.

To gain intuition, consider the following simple example. Let there be one phenotype *x* and one genotypic trait *y*, and let age structure be collapsed such that development occurs instantaneously. Collapsing age structure is a heuristic simplification, but doing so makes **x** and **y** scalars, which can then be easily visualised. Also, let there be no social interactions, no niche construction, no exogenous environmental change, and no density dependence. By removing social interactions and density dependence, the latter of which is also a heuristic simplification, evolutionary change can be not only abstractly but also visually described as the climbing of a fitness landscape. The extended mechanistic Lande Eq. (8) states that selection can be seen as “pushing” the geno-phenotype uphill on the fitness landscape in geno-phenotype space in the direction of steepest ascent, whereas genetic covariation diverts evolutionary change in a possibly less steep direction. If genetic constraints on the traits considered were not absolute (here, if **H_z_** were non-singular), the population would eventually reach a landscape peak, which is commonly implicitly or explicitly assumed. Yet, as the traits considered must be both the genotype and the phenotype arising from the developmental constraint (1), genetic constraints are necessarily absolute (i.e., **H_z_** is singular). Hence, evolution is restricted to an admissible path on the landscape where the developmental constraint is met (Fig. 1a-c; the computer code used to generate all the figures is in the Supplementary Information). Adaptive evolution may thus be understood as the climbing of the fitness landscape along an admissible path determined by development. The evolutionary process eventually reaches a path peak if there are no absolute mutational constraints (Fig. 1a-c). Formally, a path peak is a point in geno-phenotype space where the developmental constraints are met and fitness is locally and totally maximized with respect to genotype: here “totally” means after substituting in the fitness function for both the developmental and environmental constraints. Selection response vanishes at path peaks which are not necessarily landscape peaks (Fig. 1a-c). The admissible path yields an elevation profile of fitness, namely, the total fitness landscape of the genotype, on which adaptation is constrained to occur, and can have peaks and valleys created by the developmental map (Fig. 1d-f). Hence, selection pushes the genotype uphill on the total fitness landscape of the genotype, but in this profiled landscape evolutionary change is not blocked provided that there are no absolute mutational constraints. Consequently, without absolute mutational constraints, evolutionary stasis generally occurs at a peak on the total fitness landscape of the genotype, even though this does not generally correspond to a peak on the fitness landscape of the phenotype and genotype. The **H_z_**-matrix evolves as an emergent property as the resident phenotype and genotype evolve, and genetic variances and covariances may substantially increase or decrease if development is non-linear (consistent with previous individual-based simulations; Miloco and Salazar-Ciudad, 2022), rather than being approximately constant as is often considered under short-term evolution with an infinite number of loci (Fisher, 1918; Barton *et al*., 2017) which we do not assume (Fig. 1g-i). Ultimately, development constrains the path of adaptation and thus defines its outcome jointly with selection (Fig. 1j-l). Overall, development has a permanent dual role in adaptation by influencing evolutionary equilibria (blue lines in Fig. 1j-l) and determining the admissible evolutionary path (red line in Fig. 1j-l).

**Figure 1:**
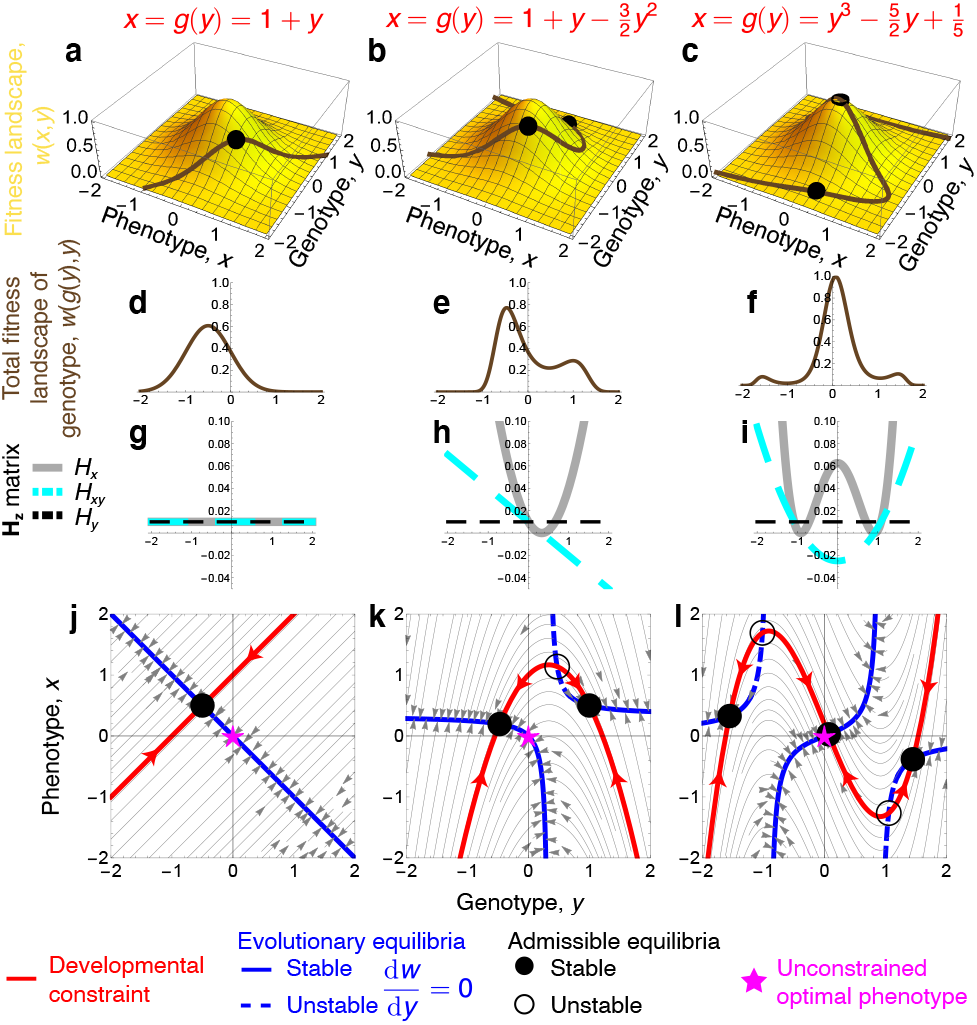
Development determines the evolutionary path. **a-c**, Fitness landscape for a phenotype *x* and a genotype *y*, with three developmental maps. The developmental map determines the path on the landscape as *x* can only take values along the path. **d-f**, The total fitness landscape of the genotype gives the elevation profile considering developmental and environmental constraints. **g-i**, Mechanistic additive genetic covariances depend on the developmental map and evolve as the genotype evolves. **j-l**, The evolutionary dynamics occur along the developmental constraint (red; gray arrows are parallel or antiparallel to leading eigenvectors of **H_z_**, called “genetic lines of least resistance”; Schluter, 1996). Evolutionary outcomes (black dots) occur at path peaks and thus depend on development. Evolutionary equilibria (blue) are infinite in number because **H_z_** is singular and occur when total genotypic selection vanishes despite persistent direct selection. The intersection of the developmental map and evolutionary equilibria yields the admissible evolutionary equilibria. Different developmental maps yield different evolutionary trajectories and outcomes with the same single-peak landscape. Developmental bias is quantified by the slope of the red line (*∂x*/*∂y*). Evolutionary change satisfies 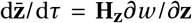, where **H_z_** = (d**z**/d*y*)*H_y_* (d**z**^⊤^/d*y*) = (d*x*/d*y*, 1)^⊤^ *H_y_* (d*x*/d*y*, 1) = *H_y_* ((d*x*/d*y*)^2^, d*x*/d*y*; d*x*/d*y*, 1) = (*H_x_*, *H_xy_*; *H_xy_*, *H_y_*), which is singular. Hence, the mechanistic additive genetic variance of *x* is *H_y_* (d*x*/d*y*)^2^. The total effects of a mutant’s genotype on her geno-phenotype are d**z**^⊤^/d*y* = (d*x*/d*y*, d*y* /d*y*) = (d*x*/d*y*, 1). Fitness is *w* (*x*, *y*) = exp(−*x*^2^ − *y*^2^) and mutational variance is *H_y_* = 0.01 (so that 0 ≤ tr(**H_y_**) ≪ 1). As we use our heuristic assumptions of collapsed age structure and no density dependence, we use here a simpler fitness function than that in Eq. (4). Functional forms are chosen for illustration.

That evolution stops at path peaks rather than landscape peaks may help explain abundant empirical data. Empirical estimation of selection has found that direct stabilizing selection is rare relative to direct directional selection across thousands of estimates in more than 50 species (Kingsolver *et al*., 2001; Kingsolver and Diamond, 2011). The rarity of stabilizing selection has been puzzling under the common assumption that evolutionary outcomes occur at landscape peaks, where stabilizing selection is prevalent (Charlesworth *et al*., 1982; Walsh and Lynch, 2018). In contrast, the rarity of stabilizing selection is consistent with evolutionary outcomes necessarily occurring at path peaks. Indeed, if path peaks occur outside landscape peaks, evolutionary outcomes occur with persistent direct directional selection (Houle, 1991; Kirkpatrick and Lofsvold, 1992; Altenberg, 1995) and relatively weak direct stabilizing selection (because direct stabilizing selection is relatively strong only near landscape peaks). Thus, evolution generally stopping at path peaks outside landscape peaks may help explain the otherwise puzzling observation of rare stabilizing selection. There are other explanations for the rarity of stabilizing selection (Haller and Hendry, 2013; Morrissey, 2015), and Morrissey’s is closely related to ours.

### Development can drive diversification

The evolution of phenotypic and genetic diversity is typically explained in terms of selection or drift (Gavrilets and Losos, 2009; Doebeli, 2011), but the relevance of development has been less clear. We find that development can drive the evolution of phenotypic and genotypic diversity, even in a constant, single-peak fitness landscape, where selection provides an evolutionary force while development translates it into diversification. This can happen in two ways. First, if the developmental map changes inducing a shifted path peak, the population can evolve different phenotypes and genotypes (Fig. 2a-f). Second, the developmental map may change, generating new path peaks, if it is or becomes non-linear, such that the population may evolve different phenotypes and genotypes depending on the initial conditions (Fig. 2g-l). Diversification may then occur if a population subdivides and one of the descendant populations undergoes changes in their developmental map due to changes in its genotype, environment, or how the phenotype is constructed. Mathematically, multiple path peaks can arise from non-linear development because it can generate bifurcations altering the number of admissible evolutionary equilibria even though the number of evolutionary equilibria remains the same (i.e., infinite). This is a form of evolutionary branching (Geritz *et al*., 1998), which may be called evo-devo branching, where the admissible path creates a valley in the total fitness landscape of the genotype so there is disruptive total genotypic selection (Fig. 1e; see also Morrissey 2015 where stabilizing total selection is analysed). There may not be a valley in the fitness landscape so no disruptive direct selection on the phenotype or genotype (Fig. 1b) as evo-devo branching only requires disruptive total genotypic selection possibly created by development rather than direct disruptive selection. Thus, evo-devo branching may occur in situations where evolutionary branching driven by direct disruptive selection would not occur. Hence, development can lead to phenotypic and genotypic diversification, even with a constant single-peak fitness landscape in geno-phenotype or phenotype space. Consequently, phenotypic and genotypic diversification might arise from the evolution of development.

**Figure 2:**
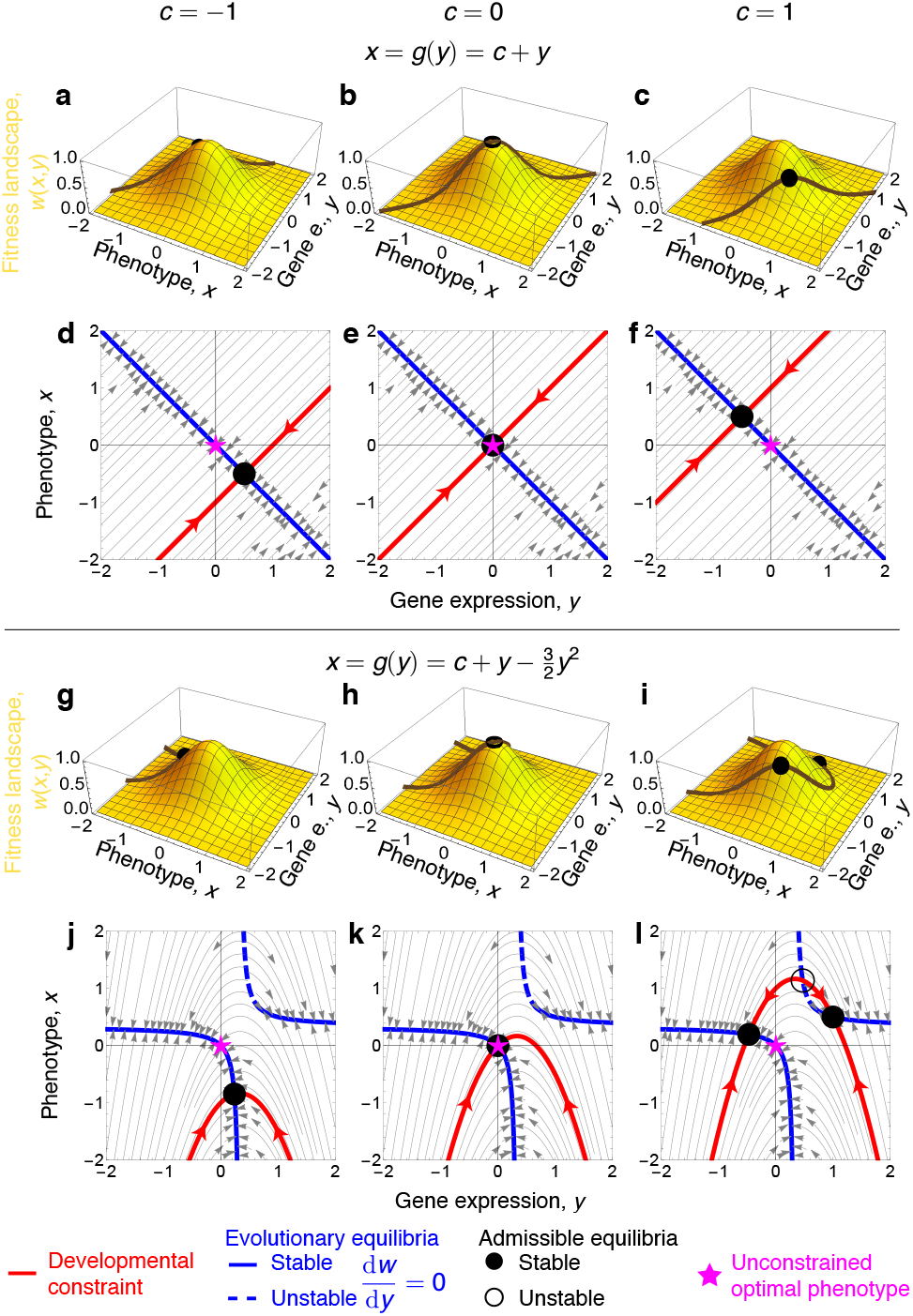
Development-driven diversification. Development may drive phenotypic and genotypic diversification in two ways. First, by shifting path peaks (**a**-**f**). Second, by creating path peaks (**g**-**l**). The evolutionary dynamics are given as in the legend of Fig. 1. As *c* changes, development *g* changes, but total developmental bias d*x*/d*y* does not change, so the evolutionary equilibria remain the same (blue); however, the admissible evolutionary equilibria do change (open and solid circles) and so does the outcome even though the fitness landscape is constant and single peaked.

Our analysis partly substantiates a classic explanation of punctuated equilibria in terms of developmental constraints, an explanation that has been dismissed in the light of previous theory. Punctuated equilibria refer to long periods of morphological stasis alternated by periods of rapid change as observed in the fossil record (Eldredge and Gould, 1972; Hunt *et al*., 2015). The dominant explanation for punctuated equilibria is that fitness landscapes remain relatively invariant for long periods so there is stabilizing selection for long periods at an unconstrained optimum, and fitness landscapes subsequently undergo substantial change particularly during adaptive radiations triggering directional selection (Charlesworth *et al*., 1982; Walsh and Lynch, 2018). A classic alternative explanation is that (a) developmental constraints prevent change for long periods and (b) revolutions in the developmental program bring sudden change (Gould, 1980). The ability of constraints to prevent change has been thought to be refuted by the common observation of selection response, sometimes under seeming developmental constraints (e.g., Beldade *et al*., 2002), while revolutions in the developmental program have been refuted on other grounds (Charlesworth *et al*., 1982; Eldredge *et al*., 2005; Klingenberg, 2010). Our analysis substantiates the view that developmental constraints may prevent change for long periods, despite possible selection response: indeed, even if developmental constraints halt a population at a path peak, selection response is possible since path peaks may change, either by change in the path (e.g., by change in the genotype, environment, or how the phenotype is constructed) or the landscape (e.g., by artificial selection as in Beldade *et al*. 2002). Thus, available evidence does not necessarily rule out that developmental constraints can prevent change for long periods in some directions. Moreover, punctuated equilibria could arise by change in the developmental map, rather than only by change of the fitness landscape. In particular, gradual change in the developmental map as the genotype, phenotype, or environment gradually evolve may yield sudden path-peak creation events (sudden since such events are bifurcations, which are sudden by definition; Fig. 2g-l; see also Fig. 3.4 of Metz 2011). Path-peak creation events yielding sudden phenotypic diversification might then generate punctuated equilibria, arising by gradual evolution of the developmental map, without revolutions in the developmental program. Furthermore, by shifting the admissible path, the evolution of development may similarly lead to the crossing of valleys in a multi-peak fitness landscape if a path peak moves from a landscape peak to another. However, a factor limiting adaptation is valleys in the total fitness landscape of the genotype.

### Niche construction reshapes the path

Niche construction — whereby organisms affect their environment — has been suggested to be a developmental factor having major evolutionary effects (Odling-Smee *et al*., 1996; Laland *et al*., 2014, 2015). How does niche construction affect evolution? We find that niche construction has a dual effect on adaptation. First, niche construction (*∂**ϵ***^⊤^/*∂***z**) reshapes the fitness landscape as previously known (Odling-Smee *et al*., 1996; Laland *et al*., 2014, 2015). Second, niche construction reshapes the path on the landscape by altering absolute genetic constraints.

To see this, we relax the assumption that niche construction is absent (*∂**ϵ***^⊤^/*∂***z** ≠ **0**), whilst maintaining our assumptions of (ii) no social development 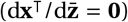, and (iii) no exogenous environmental change (*∂**ϵ***/*∂τ* = **0**). The evolutionary dynamics of the resident phenotype and genotype are then given by

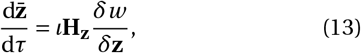

where **H_z_** guarantees that the developmental (1) and environmental (2) constraints are met at all times. Eq. (13) depends on the total immediate selection gradient of the phenotype and genotype

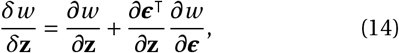

which measures total immediate selection considering environmental constraints but not developmental constraints. So, selection pushes uphill, no longer on the fitness landscape that ignores environmental constraints, but on the reshaped total immediate fitness landscape that considers such constraints (Fig. 3a-e). Eq. (14) shows that such reshaping of the fitness landscape is done by the interaction of niche construction and environmental sensitivity of selection (*∂w* /*∂**ϵ***) (a term coined by Chevin *et al*. 2010 for a more specific notion). The total immediate fitness landscape can have new peaks and valleys relative to the fitness landscape. Additionally, Eq. (13) depends on **H_z_**, which still has the form in Eq. (9) so it is still singular but now depends on the interaction of niche construction and plasticity (*∂***x**^⊤^/*∂**ϵ***). Thus, development determines the path on the total immediate fitness landscape, but now this path is reshaped by the interaction of niche construction and plasticity (Fig. 3f-j). Consequently, niche construction reshapes the fitness landscape by interacting with environmental sensitivity of selection, and reshapes the admissible path on the landscape by interacting with phenotypic plasticity. Thus, niche construction affects both selection and development, where selection and development jointly determine the evolutionary outcomes if there are no absolute mutational constraints.

**Figure 3:**
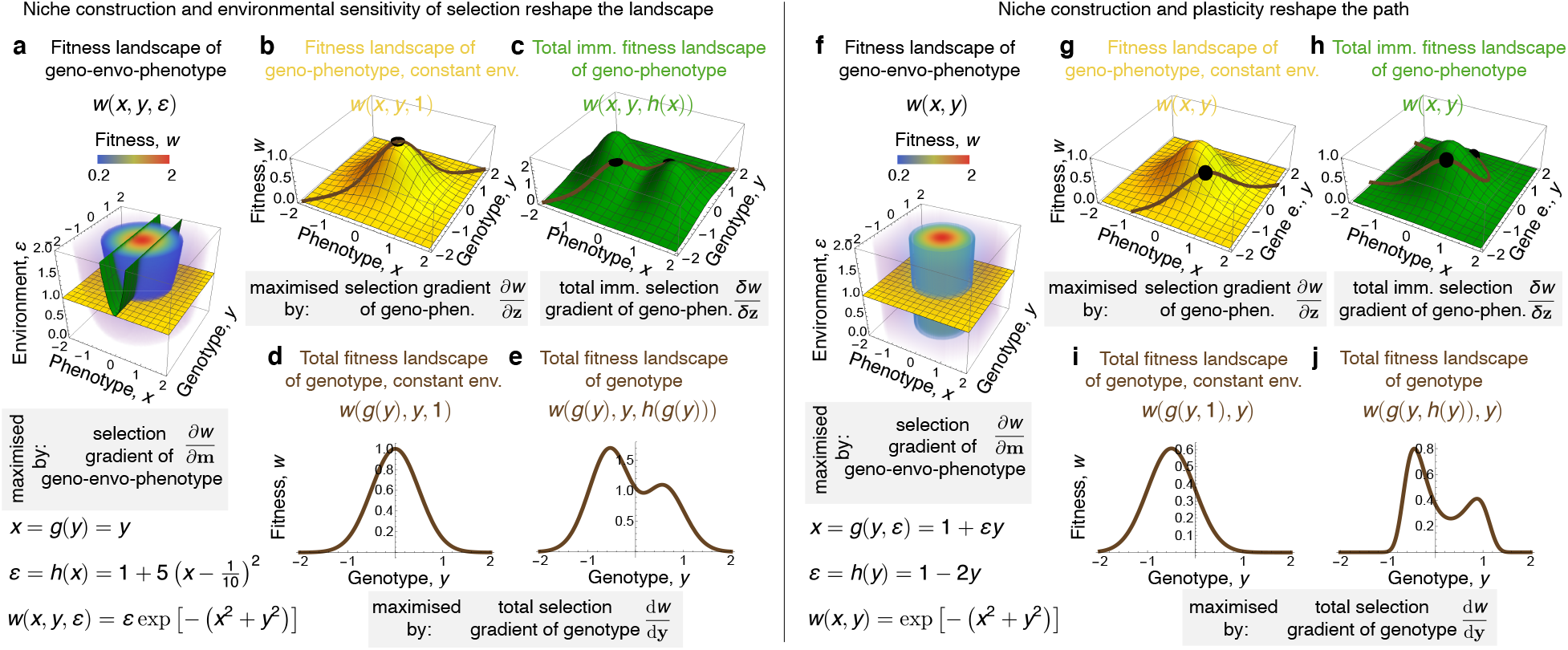
Niche construction reshapes the fitness landscape and the evolutionary path. **a-e**, There is niche construction by the phenotype *x* which non-linearly affects the environmental trait, which in turn increases fitness. **a**, Fitness landscape vs. phenotype, genotype, and environment. Slices for constant environment (yellow) and for the environmental constraint (green) are respectively shown in **b** and **c**. The interaction of niche construction (*∂ϵ* /*∂x*) and environmental sensitivity of selection (*∂w* /*∂ϵ*) reshapes the landscape (from **b**to **c**) by affecting total immediate selection *δw* /*δ***z** but it does not affect genetic covariation **H_z_**. **f-j**, There is niche construction by the genotype *y* which linearly affects the environmental trait, which in turn increases the phenotype. The interaction of niche construction (*∂ϵ* /*∂y*) and plasticity (*∂x*/*∂ϵ*) reshapes the path (from **g** to **h**). This interaction affects genetic covariation **H_z_** but does not affect total immediate selection *δw* /*δ***z**. In all panels, evolutionary change satisfies 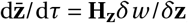, where *δw* /*δ***z** = *∂w* /*∂***z** + (*∂ϵ* /*∂***z**) (*∂w* /*∂ϵ*) and **H_z_** is given in Fig. 1. In **a-e**, d*x*/d*y* = *∂x*/*∂y*, whereas in **f-j**, d*x*/d*y* = *∂x*/*∂y* + (*∂ϵ* /*∂y*)(*∂x*/*∂ϵ*).

Eq. (13) could also be seen as corresponding to an extended Lande equation describing genotypic and phenotypic evolution where environmental traits are not explicitly considered in a quantitative genetics analysis. Now, Eq. (13) generally depends on the resident environment 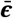, which evolves due to niche construction. Provided that the environmental map is known or assumed, the environment is then known so Eq. (13) still yields a dynamically sufficient description of longterm phenotypic evolution despite environmental evolution. Yet, if one is interested in understanding how the evolution of the environment proceeds on the fitness landscape, an equation describing the evolution of the environment in gradient form would be needed. González-Forero (2021) derives such an equation, showing it has the form of a further extended mechanistic Lande equation describing the evolutionary dynamics of the geno-envo-phenotype with an associated mechanistic additive genetic covariance matrix that is also necessarily singular for the same reasons given above.

### Social development alters the outcome

Extra-genetic inheritance, including social learning and epigenetic inheritance, has also been suggested to be a developmental factor having major evolutionary effects (Baldwin, 1896; Laland *et al*., 2014, 2015). Current understanding is that extra-genetic inheritance has only transient evolutionary effects if elements so inherited are transmitted over only a few generations (e.g., Jablonka and Lamb, 2014; Charlesworth *et al*., 2017; Walsh and Lynch, 2018; Beltran *et al*., 2020). For instance, the permanent evolutionary effects discussed by Day and Bon-duriansky (2011) and Bonduriansky and Day (2018) involve the coevolution of genetic and extra-genetic elements, which requires transmission of extra-genetic elements over multiple generations. In contrast, we find that extra-genetic inheritance can have permanent evolutionary effects even without transmission of extra-genetic elements over multiple generations, because it changes development and so the absolute genetic constraints and the admissible evolutionary path.

Extra-genetic inheritance can be seen as a particular form of social development 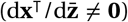, where the developed phenotype depends on the phenotype and genotype of social partners. For instance, acquiring a skill or epimark from a role model or a parent, respectively, involves an interaction between the recipient and donor, and these extra-genetic elements affect the subsequent phenotype of the individual (although recall we assume that social interactions are only with non-relatives for simplicity). Social development can also mechanistically describe both indirect genetic effects, where the genes of social partners affect an individual’s phenotype (Moore *et al*., 1997), and social phenotypic effects that need not stem from genes (e.g., if the individual’s developed traits depend on social partners’ environmentally induced developed traits). How does social development, including extra-genetic inheritance, affect evolution? We find that, since it alters the developmental map, social development alters the evolutionary outcome (i.e., the admissible evolutionary equilibria) by altering the admissible evolutionary path without necessarily altering the evolutionary equilibria themselves. Thus, extra-genetically inherited elements can have permanent evolutionary effects without being transmitted over many generations. What is needed is that the ability to acquire the (possibly evolving) extra-genetic element persists over many generations so that individuals have a consistently socially altered development over many generations. To see this, consider the following.

Social development introduces a complication to evoutionary analysis as follows. With social development, the developed phenotype depends on the frequency of other phenotypes and genotypes and even if a phenotype is fixed in the population, offspring may develop a different phenotype if the fixed parental phenotype developed in a different social context (i.e., the phenotype may not breed true despite clonal reproduction). We handle this complication with the following notions. We say a genotype, phenotype, and environment **m**\*\* (**x**\*\*; **y**\*\*; ***ϵ***\*\*) is a socio-devo equilibrium if and only if an individual with such genotype and environment developing in the context of individuals having such genotype, phenotype, and environment develops their same phenotype. A socio-devo equilibrium is socio-devo stable if and only if socio-devo dynamics are locally stable to small perturbations in the phenotype (González-Forero, 2021). A phenotype in socio-devo stable equilibrium breeds true when fixed. Although social development can yield socio-devo unstable phenotypes, our analyses apply only to socio-devo stable phenotypes (which are guaranteed to be so if all the eigenvalues of 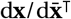 have absolute value strictly less than one, which we assume). Thus, for socio-devo stable phenotypes, their evolutionary dynamics are described by the following equations. Allowing for niche construction (*∂**ϵ***^⊤^/*∂***z** ≠ 0) social development 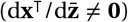, but assuming (iii) that there is no exogenous environmental change (*∂**ϵ***/*∂τ* = **0**), then the evolutionary dynamics of the resident phenotype and genotype are given by

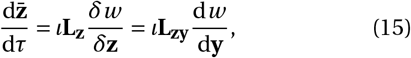

where both **L_z_** and **L_zy_** guarantee that the developmental (1) and environmental (2) constraints are met at all times. This equation now depends on the mechanistic additive socio-genetic cross-covariance matrix of the geno-phenotype (L for legacy)

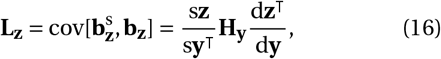

where the stabilized mechanistic breeding value of **z** is defined as

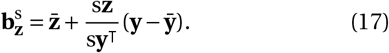

The matrix s**z**/s**y**^⊤^ is the stabilized total derivative of **z** with respect to **y**^⊤^, which measures the total effects of **y**^⊤^ on **z** after the effects of social development have stabilized in the population; s**z**/s**y**^⊤^ reduces to d**z**/d**y**^⊤^ if development is not social. The **L_z_**-matrix generalizes **H_z_** by including the effects of social development, so if development is non-social **L_z_** reduces to **H_z_** (**L_z_** is analogous to another generalization of the **G**-matrix used in the indirect genetic effects literature; Moore *et al*., 1997). As for **H_z_**, the matrix **L_z_** is always singular because the geno-phenotype contains the genotype (i.e., because d**z**^⊤^/d**y** has fewer rows than columns). Hence, also under social development, evolutionary stasis can generally occur with persistent total immediate selection on the phenotype and genotype. In turn, the second equality in Eq. (15) depends on the additive socio-genetic cross-covariance matrix between the geno-phenotype and genotype

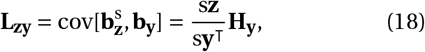

which generalizes the corresponding matrix **H_zy_** we had in Eq. (12) to consider social development. Similarly to **H_zy_**, this matrix **L_zy_** is non-singular if **H_y_** is non-singular because the genotype is developmentally independent by assumption. Thus, evolutionary equilibria still occur when total genotypic selection vanishes (d*w* /d**y** = **0**) if there are no absolute mutational constraints.

The singularity of **L_z_** entails that social development, including extra-genetic inheritance, may alter both the evolutionary equilibria (by altering d*w* /d**y**) and the admissible path (by altering **L_zy_**; Fig. 4). Note that *ȳ* in Fig. 4**a**,**b** can be interpreted as being an extra-genetically acquired element that modifies development, but such acquired effect is not transmitted to future generations as the individual transmits the *y* it has. Yet, such extra-genetically acquired element alters the developmental constraint and evolutionary equilibria, thus altering the evolutionary outcome relative to that without extra-genetic inheritance (Fig. 1**a**,**d**,**g**,**j**). Alternatively, social development may not affect the evolutionary equilibria if total genotypic selection d*w* /d**y** is independent of social partners’ geno-phenotype 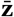, but social development may still affect the admissible evolutionary path given by the developmental constraint in Eq. (1) (by altering only **L_zy_**; Fig. 5). Similarly, *ȳ* in Fig. 5**a**,**b** can also be interpreted as being an extra-genetically acquired element that modifies development, but it is not transmitted to future generations and still alters the evolutionary outcome relative to that without extra-genetic inheritance (Fig. 1**b**,**e**,**h**,**k**). Indeed, in both cases, social development (e.g., extra-genetic inheritance) affects the evolutionary path and so the admissible evolutionary equilibria and hence the evolutionary outcome. Mathematically, this results because **L_z_** is singular, so changing the path generally changes the outcome. Thus, because **L_z_** is singular, extra-genetic inheritance can have permanent evolutionary effects even if extra-genetically acquired elements are not transmitted to other individuals.

**Figure 4:**
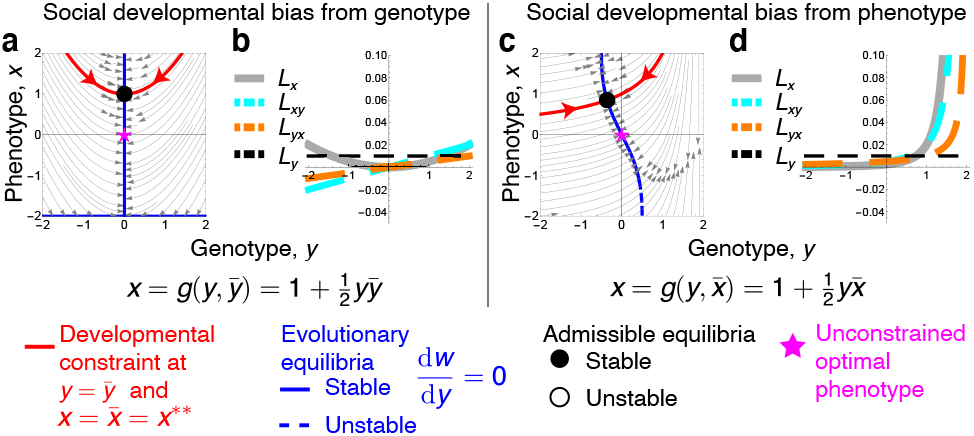
Social development alters the evolutionary outcome. **a-b**, There is social development, particularly social developmental bias from the genotype (*∂x*/*∂ȳ* ≠ 0). Here social development introduces a non-linearity relative to the developmental map of Fig. 1**a**,**d**,**g**,**j**, which changes the evolutionary equilibria (blue), admissible path (red), and outcome (solid dot). **c-d**, As in **a-b**, but there is social developmental bias from the phenotype 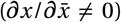. The developmental map is that of **a-b** replacing *ȳ* with 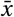. This also changes the evolutionary equilibria, admissible path, and outcome. In all panels, evolutionary change satisfies 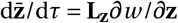, where *w* (*x*, *y*) = exp(−*x*^2^ − *y*^2^) and **L_z_** = (s**z**/s*y*)*H_y_* (d**z**^⊤^/d*y*) = (*L_x_*, *L_xy_*; *L_yx_*, *L_y_*), with s**z**/s*y* = (s*x*/s*y*; 1). For **a-b**, s*x*/s*y* = d*x*/d*y* + d*x*/d*ȳ* (from Layer 5, Eq. 2a in González-Forero 2021). Similarly, for **c-d**, 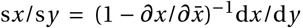, with d**z**^⊤^/d*y* and s*x*/s*y* evaluated at the socio-devo equilibrium *x*\*\* that solves *x*\*\* = *g* (*y*, *x*\*\*). Since development is social, a fitness landscape visualization as in Fig. 1a is not possible. In **c**, the stream plot (gray arrows) is only locally accurate around the developmental constraint where the socio-devo equilibrium holds. The 1/2 in the developmental constraint is used to meet the assumption 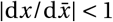 but does not affect the points made.

**Figure 5:**
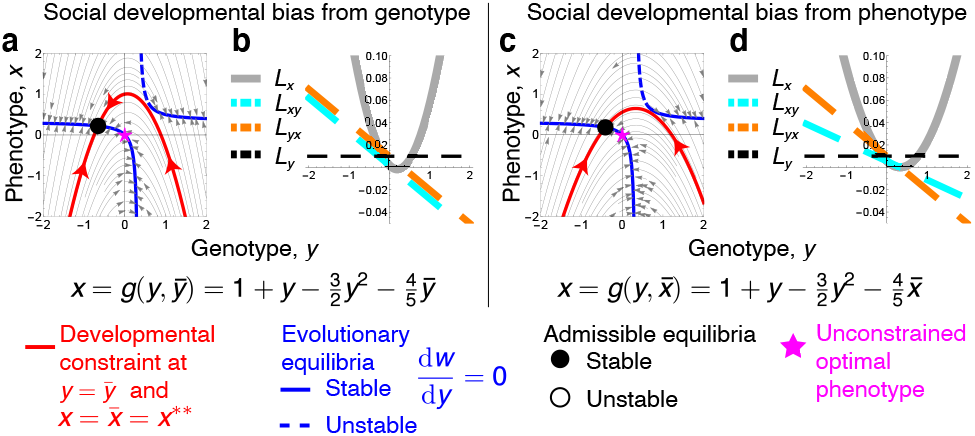
Social development can alter the evolutionary outcome without altering evolutionary equilibria. The plots are as in Fig. 4, except that now the developmental maps are a linear modification of those in Fig. 1**b**,**e**,**h**,**k**. Relative to that figure, social development does not alter total developmental bias d*x*/d*y* so it does not affect evolutionary equilibria (blue). However, social development affects the admissible path (red) and so the evolutionary outcome (solid dot). Note *L_x_* can be negative in **b** as it gives the covariance between the stabilized mechanistic breeding value of *x* and the mechanistic breeding value of *x* (in contrast, *H_x_* gives the variance of the mechanistic breeding value of *x*, which is non-negative). The 4/5 in the developmental constraint is used to meet the assumption 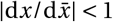 but does not affect the points made.

When social development alters only the admissible path, it does so because of social developmental bias 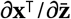; e.g., extra-genetic inheritance and indirect-genetic effects), and/or the interaction of social niche construction 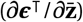 and plasticity (*∂***x**^⊤^/*∂**ϵ***) (from Layer 3, Eq. 3 of González-Forero 2021).

### Plastic response alters the path

Research on the evolutionary effects of plasticity has intensified, with a particular focus on whether adaptive plastic change precedes genetic evolution (Waddington, 1961; West-Eberhard, 2003). How does plasticity affect evolution? We find that plasticity also has a dual evolutionary effect. First, we saw that plasticity (*∂***x**^⊤^/*∂**ϵ***) alters the evolutionary path by interacting with niche construction (i.e., endogenous environmental change; *∂**ϵ***^⊤^/*∂***z**). This is because niche construction alters the environment, which through plasticity alters the developed phenotype and so the evolutionary path (by altering **L_z_**). Second, plasticity alters the evolutionary path by interacting with exogenous environmental change (e.g., eutrophication or climate change caused by other species; *∂**ϵ***/*∂τ*). This is similarly because exogenous environmental change alters the environment, which through plasticity alters the developed phenotype and so the evolutionary path (but not by directly altering **L_z_**). With exogenous environmental change, the evolutionary dynamics comprise selection response and exogenous plastic response (see also Chevin *et al*., 2010), the latter of which modifies the developmental map and thus the evolutionary path as evolutionary time advances. Through exogenous plastic response, the evolutionary effects of development go beyond modulating genetic covariation. Exogenous plastic response has some ability to increase adaptation separate from selection response via selective development, although this ability is limited in various ways.

To see this, assume now that there is niche construction (*∂**ϵ***^⊤^/*∂***z** ≠ **0**), social development 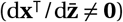, and exogenous environmental change (*∂**ϵ***/*∂τ* ≠ **0**). Then, the evolutionary dynamics of the resident phenotype and genotype are given by

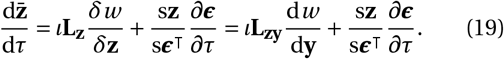

The matrix s**z**/s***ϵ***^⊤^ describes what we call stabilized plasticity of the phenotype and genotype. Stabilized plasticity includes the effects of socio-devo stabilization and reduces to total plasticity d**z**/d***ϵ***^⊤^ if development is non-social. The first term in Eq. (19) is selection response (*ι***L_z_***δw* /*δ***z**) whereas the second term is exogenous plastic response ((s**z**/s***ϵ***^⊤^)(*∂**ϵ***/*∂τ*)), which makes the developmental map evolve as the environment exogenously changes (Fig. 6). Eq. (19) shows that exogenous plastic response can induce selection response, which allows for plastic change to preceed genetic change (Waddington, 1961; West-Eberhard, 2003). For example, if the population is at a path peak so there is no selection response (**L_z_***δw* /*δ***z** = **0**) and there is exogenous plastic response, this generally changes the path thus inducing future selection response (e.g., in Fig. 6**a**, selection response is absent at the two path peaks at *x* = 1/2 at *τ* = 0, but environmental change and plasticity change the path creating selection response at these points and the path peaks move approximately to *x* = 0 and *x* = 0.18 at *τ* = 1000 in Fig. 6**c**). In the idealized case where development is selective in the sense that the developmental map is a function of the selection gradient, stabilized plasticity can depend on the selection gradient. With selective development, exogenous plastic response can induce evolution either towards or away landscape peaks depending on the developmental constraints and initial conditions, as the population is trapped in local peaks of the total fitness landscape of the genotype (Fig. 6). Exogenous plastic response with selective development has a limited ability to induce evolution towards a landscape peak since plasticity must depend on developmental-past environments, which induces a developmental lag in the plastic response (Supplementary Information section S2). Additionally, exogenous plastic response with selective development may induce evolution away a landscape peak when the rate of environmental change switches sign (Supplementary Information section S2).

**Figure 6:**
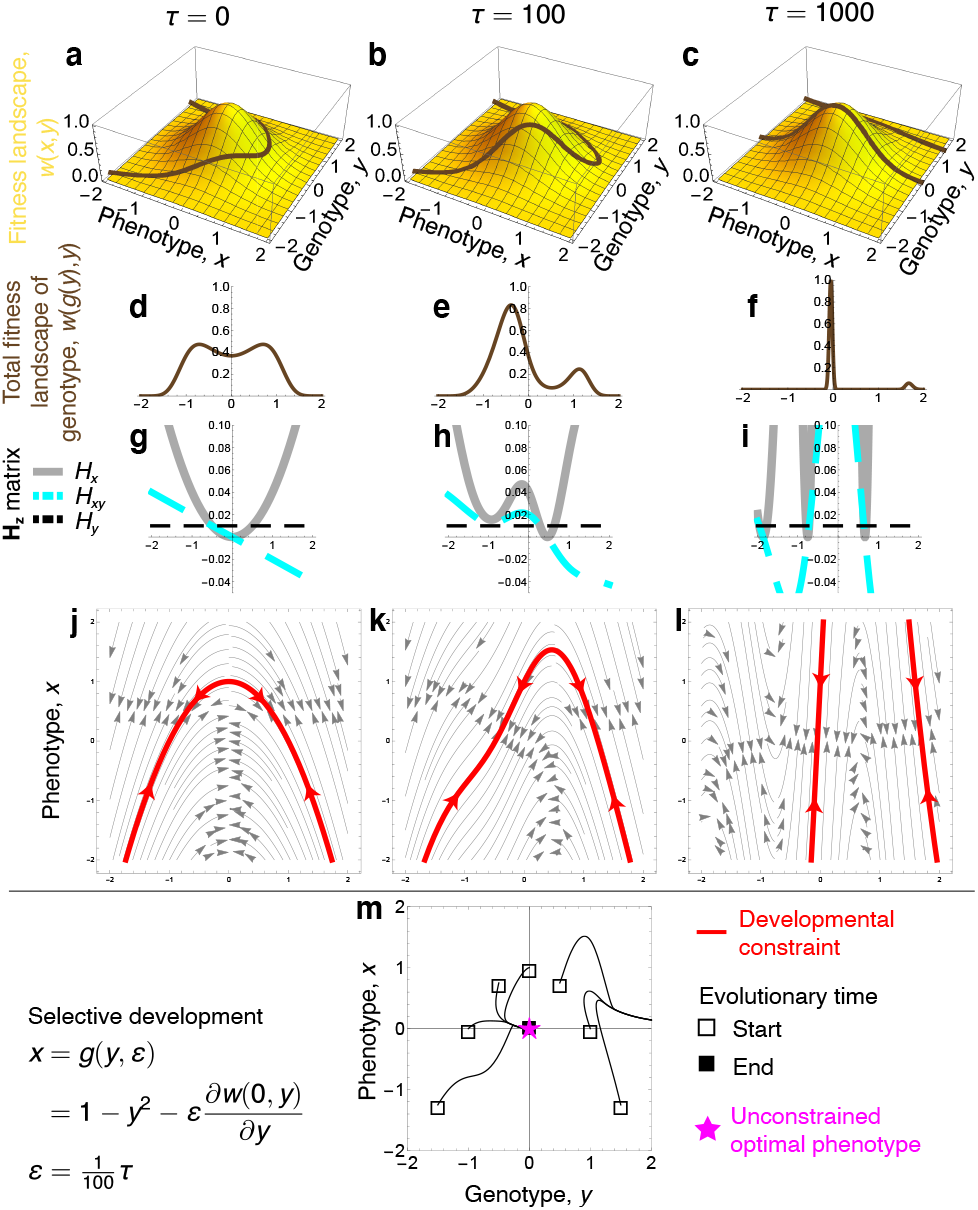
Exogenous plastic response changes the evolutionary path. We let there be exogenous environmental change and plasticity, but no social development, so **L_z_** reduces to **H_z_** and no niche construction, so plasticity does not affect **H_z_** but only the exogenous plastic response. For illustration, development is selective by letting plasticity equal the selection gradient at the optimal phenotype, *∂x*/*∂ϵ* = *∂w* (0, *y*)/*∂y*. **a-l**, Exogenous environmental change induces exogenous plastic response which raises part of the path on the landscape. **m**, Resulting evolutionary dynamics showing that selective development induces some evolutionary trajectories to converge to the landscape peak, whereas others do not by getting trapped in another path peak due to their initial condition. Evolutionary change satisfies 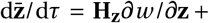 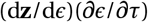, where d**z**/d*ϵ* = (d*x*/d*ϵ*; d*y* /d*ϵ*) = (*∂x*/*∂ϵ*; 0) and **H_z_** is given in Fig. 1. Fitness is *w* (*x*, *y*) = exp(−*x*^2^ − *y*^2^).

### Development enables negative senescence

Thus far, we have considered evolutionary effects of development that occur even without considering that development takes time. Indeed, in our illustrative examples we have let age structure be collapsed so development happens instantaneously. We now show that explicit consideration of age progression in development may help explain an otherwise puzzling ageing pattern.

Individuals may show senescence as they age, that is, decreasing survival or fertility after the onset of reproduction. Leading hypotheses for the evolution of senescence are based on the fact that the forces of selection decline with age (*ϕ_a_* and *π_a_* in Eq. (4), and Eqs. S8 in the Supplementary Information; Hamilton, 1966). For example, if a mutation has a beneficial effect on survival or fertility early in life and a pleiotropic, deleterious effect of similar magnitude late in life, the mutation would be favoured (Medawar, 1952; Williams, 1957). The universality of declining forces of selection has suggested that senescence should be universal (Hamilton, 1966), at least for organisms with a bodily reset at conception (Lehtonen, 2020). However, empirical research has increasingly reported organisms with negative senescence (Jones *et al*., 2014b), that is, where survival or fertility increase with age after the onset of reproduction, and models have suggested that this may stem from indeterminate growth (Vaupel *et al*., 2004; Lehtonen, 2020).

Consistently with the latter, we find that developmental propagation — whereby an early-acting mutation has a large phenotypic effect late in life — can drive the evolution of negative senescence. For example, if a mutation in genotypic trait *i* has a deleterious effect on survival or fertility at an early age *a*, so that 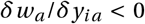, and a pleiotropic, beneficial effect on survival or fertility of similar magnitude at a later age *j* > *a*, so that 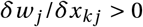 for phenotype *k*, then total immediate selection is weaker at the later age because of declining selection forces (i.e., 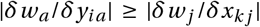). Yet, crucially, such late-beneficial mutation is still favoured if total genotypic selection on it is positive, that is, if

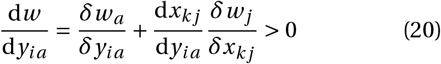

(Supplementary Information section S3), where the effect of the early mutation on the late phenotype is

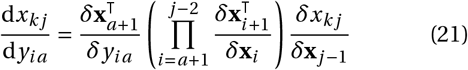

from Eq. C17 of González-Forero 2021). This effect of the early mutation on the late phenotype may increase (or decrease) with the phenotype age *j* because of developmental propagation of phenotypic effects as age advances, a propagation that is described by the term in parentheses in Eq. (21): in particular, indeterminate growth allows for indefinite developmental propagation. So total genotypic selection may increase with the age *j* at which the pleiotropic effect is expressed. Consequently, a mutation with late beneficial effects may be favoured despite early deleterious effects. Thus, developmental propagation may enable selection to shape traits at advanced age, allowing for adaptive evolution of increasing survival or fertility as ageing proceeds despite decreasing selection forces after the onset of reproduction. This suggests that allowing for developmental propagation in models with explicit development in which negative senescence fails to evolve (Nelson and Masel, 2017) might still enable negative senescence (perhaps via cell maintenance).

## Discussion

Evolution by natural selection can be understood as the climbing of a fitness landscape under some assumptions. This was first established for genetic evolution (Wright, 1937) and was later established for phenotypic evolution under the tacit assumption that there is negligible genetic evolution (Lande, 1976, 1979; Barton *et al*., 2017; Hill, 2017; Walsh and Lynch, 2018). However, long-term phenotypic evolution involves non-negligible genetic evolution. We have shown that understanding long-term phenotypic evolution as the climbing of a fitness landscape requires that genotypic and phenotypic evolution are considered simultaneously. Doing so necessarily makes the associated constraining matrix (**H**, **L**, or **G**) singular because the phenotype is related to the genotype via development. Thus, long-term phenotypic evolution necessarily involves absolute genetic constraints, where the evolutionary path is given by the developmental process that associates genes to phenotype.

Consequently, we have shown that development affects evolution by determining the evolutionary path. This sharpens the basic principle of adaptation as the climbing of the fitness landscape (Lande, 1979) (sensu Dieckmann and Law 1996), such that evolutionary outcomes occur at best at peaks on the path rather than on the landscape. Developmental constraints thus stop evolution at path peaks that can happen mid-way on a fitness slope. Thus, selection and development jointly define the evolutionary outcomes if there are no absolute mutational constraints. Selective development, if feasible, can bring some of the path closer to the landscape peak by means of exogenous plastic response. Our findings substantiate several although not all intuitions previously given in relation to the role of development in evolution (Waddington, 1959; Gould and Lewontin, 1979; West-Eberhard, 2003; Pigliucci, 2007; Jablonka and Lamb, 2014; Laland *et al*., 2014, 2015). In particular, seemingly disparate development-related factors previously suggested to be evolutionarily important (Laland *et al*., 2014, 2015) — namely phenotypic plasticity (West-Eberhard, 2003), niche construction (Odling-Smee *et al*., 1996), extra-genetic inheritance (Jablonka and Lamb, 2014), and developmental bias (Arthur, 2004) — variously alter development, and so the absolute genetic constraints and consequently the evolutionary outcome. Our analysis offers answers to various major questions, namely the origin of diversity, the punctuated fossil record, the paradox of stasis, the rarity of stabilizing selection, and the origin of negative senescence. Our observations entail that an understanding of development is instrumental for evolutionary understanding: change in development alone changes evolutionary outcomes by changing absolute socio-genetic constraints (**L_z_**), even if direct (*∂w* /*∂***z**) or total (d*w* /d**z** or d*w* /d**y**) selection remain constant (Fig. 5). These observations call for empirical estimation of developmental maps, for instance, via the rapidly developing methods to estimate dynamic equations from data (Schmidt and Lipson, 2009; Brunton *et al*., 2016; Ghadami and Epureanu, 2022, and papers in the special issue). Overall, our analysis finds that development has major evolutionary consequences.

## Supporting information

Computer Code

## Acknowledgements

I thank Andy Gardner for extensive support throughout this project, by discussing, reading, and commenting on the many drafts, and offering interpretation, advice, and funding. I thank N.W. Bailey, M.W. Feldman, K.N. Laland, R. Lande, L.C. Mikula, A.J. Moore, and M.B. Morrissey for comments on previous versions of the manuscript, and J.A.J. Metz, I. Salazar-Ciudad, and D.M. Shuker for discussion. I thank J. Masel, M. Pavlicev, and two anonymous reviewers for detailed criticism. This work was funded by an ERC Consolidator Grant to A. Gardner (grant no. 771387), by the School of Biology of the University of St Andrews, and by a John Templeton Foundation grant to K.N. Laland and T. Uller (grant ID 60501). The author has no conflict of interest to declare.

## References

Alberch, P., Gould, S.J., Oster, G.F. and Wake, D.B. (1979). Size and shape in ontogeny and phylogeny. Paleobiology, 5, 296–317.

Altenberg, L. (1995). Genome growth and the evolution of the genotype-phenotype map. In W. Banzhaf and F. H. Eeckman, editors, Evolution and biocomputation, volume 899 of Lecture Notes in Computer Science, pages 205–259. Springer-Verlag.

Arthur, W. (2004). Biased Embryos and Evolution. Cambridge Univ. Press, Cambridge, UK.

Baldwin, J.M. (1896). A new factor in evolution. Am. Nat., 30, 441–451.

Barton, N.H. and Turelli, M. (1987). Adaptive landscapes, genetic distance and the evolution of quantitative characters. Genet. Res., 49, 157–173.

Barton, N.H., Etheridge, A.M. and Véber, A. (2017). The infinitesimal model: definition, derivation, and implications. Theor. Popul. Biol., 118, 50–73.

Beldade, P., Koops, K. and Brakefield, P.M. (2002). Developmental constraints versus flexibility in morphological evolution. Nature, 416, 844–847.

Beltran, T., Shahrezaei, V., Katju, V. and Sarkies, P. (2020). Epimutations driven by small RNAs arise frequently but most have limited duration in Caenorhabditis elegans. Nat. Ecol. Evol., 4, 1539–1548.

Bonduriansky, R. and Day, T. (2018). Extended Heredity. Princeton Univ. Press, Princeton, NJ, USA.

Brunton, S.L., Proctor, J.L. and Kutz, J.N. (2016). Discovering governing equations from data by sparse identification of non-linear dynamical systems. Proc. Natl. Acad. Sci. USA, 113, 3932–3937.

Carroll, S.B. (2008). Evo-devo and an expanding evolutionary synthesis: a genetic theory of morphological evolution. Cell, 134, 25–36.

Caswell, H. (1982). Optimal life histories and the agespecific costs of reproduction. J. Theor. Biol., 98, 519–529.

Caswell, H. (2001). Matrix Population Models. Sinauer, Sunderland, MA, USA, 2nd edition.

Chantepie, S. and Chevin, L.M. (2020). How does the strength of selection influence genetic correlations? Evol. Lett., 4–6, 468–478.

Charlesworth, B. (1990). Optimization models, quantitative genetics, and mutation. Evolution, 44, 520–538.

Charlesworth, B. (1994). Evolution in age-structured populations. Cambridge Univ. Press, 2nd edition.

Charlesworth, B., Lande, R. and Slatkin, M. (1982). A neo-Darwinian commentary on macroevolution. Evolution, 36, 474–498.

Charlesworth, D., Barton, N.H. and Charlesworth, B. (2017). The sources of adaptive variation. Proc. R. Soc. B, 284, 20162864.

Cheverud, J.M. (1984). Quantitative genetics and developmental constraints on evolution by selection. J. Theor. Biol., 110, 155–171.

Chevin, L.M., Lande, R. and Mace, G.M. (2010). Adaptation, plasticity, and extinction in a changing environment: Towards a predictive theory. PLOS Biology, 8(4), 1–8.

Day, T. and Bonduriansky, R. (2011). A unified approach to the evolutionary consequences of genetic and non-genetic inheritance. Am. Nat., 178, E18–E36.

Dieckmann, U. and Law, R. (1996). The dynamical theory of coevolution: a derivation from stochastic ecological processes. J. Math. Biol., 34(5), 579–612.

Dieckmann, U., Heino, M. and Parvinen, K. (2006). The adaptive dynamics of function-valued traits. J. Theor. Biol., 241, 370–389.

Doebeli, M. (2011). Adaptive Diversification. Princeton Univ. Press, Princeton, NJ, USA.

Durinx, M., (Hans) Metz, J.A.J. and Meszéna, G. (2008). Adaptive dynamics for physiologically structured population models. J. Math. Biol., 56, 673–742.

El-dredge, N. and Gould, S.J. (1972). Punctuated equilibria: an alternative to phyletic gradualism. In J. M. Thomas, editor, Models in Paleobiology, pages 82–115. Free-man, Cooper and Company.

Eldredge, N., Thompson, J.N., Brakefield, P.M., Gavrilets, S., Jablonski, D., Jeremy et al (2005). The dynamics of evolutionary stasis. Paleobiology, 31, 133–145.

Engen, S. and Sæther, B.E. (2021). Structure of the G-matrix in relation to phenotypic contributions to fitness. Theor. Popul. Biol., 138, 43–56.

Estes, S. and Arnold, S.J. (2007). Resolving the paradox of stasis: models with stabilizing selection explain evolutionary divergence on all timescales. Am. Nat., 169, 227–244.

Falconer, D.S. and Mackay, T.F.C. (1996). Introduction to Quantitative Genetics. Pearson Prentice Hall, Harlow, England, 4th edition.

Feldman, M.W. and Zhivotovsky, L.A. (1992). Gene-culture coevolution: toward a general theory of vertical transmission. Proc. Natl. Acad. Sci. USA, 89, 11935–11938.

Fisher, R. (1918). XV.—The correlation between relatives on the supposition of Mendelian inheritance. Trans. Roy. Soc. Edinb., 52, 399–433.

Gavrilets, S. and Losos, J.B. (2009). Adaptive radiation: Contrasting theory with data. Science, 323, 732–737.

Geritz, S.A.H., Kisdi, E., Meszéna, G. and Metz, J.A.J. (1998). Evolutionarily singular strategies and the adaptive growth and branching of the evolutionary tree. Evol. Ecol., 12, 35–57.

Geritz, S.A.H., Metz, J.A.J. and Rueffler, C. (2016). Mutual invadability near evolutionarily singular strategies for multivariate traits, with special reference to the strongly convergence stable case. J. Math. Biol., 72, 1081–1099.

Ghadami, A. and Epureanu, B.I. (2022). Data-driven prediction in dynamical systems: recent developments. Phil. Trans. R. Soc. A, 380, 20210213.

Gold-schmidt, R.B. (1940). The Material Basis of Evolution. Yale Univ. Press, New Haven, CT, USA.

González-Forero, M. (2021). A mathematical framework for evo-devo dynamics. In review at Theor. Popul. Biol. Preprint: https://www.biorxiv.org/content/10.1101/2021.05.17.4

Gould, S.J. (1980). Is a new and general theory of evolution emerging? Paleobiology, 6, 119–130.

Gould, S.J. and Lewontin, R.C. (1979). The spandrels of San Marco and the Panglossian paradigm: a critique of the adaptationist programme. Proc. R. Soc. Lond. B, 205, 581–598.

Haller, B.C. and Hendry, A.P. (2013). Solving the paradox of stasis: squashed stabilizing selection and the limits of detection. Evolution, 68, 483–500.

Hamilton, W.D. (1966). The moulding of senescence by natural selection. J. Theor. Biol., 12, 12–45.

Hansen, T.F. and Wagner, G.P. (2001). Modeling genetic architecture: a multilinear theory of gene interaction. Theor. Popul. Biol., 59, 61–86.

Hill, W.G. (2017). “Conversion” of epistatic into additive genetic variance in finite populations and possible impact on long-term selection response. J. Anim. Breed. Genet., 134, 196–201.

Houle, D. (1991). Genetic covariance of fitness correlates: what genetic correlations are made of and why it matters. Evolution, 45, 630–648.

Houle, D. (2001). Characters as the units of evolutionary change. In G. P. Wagner, editor, The Character Concept in Evolutionary Biology, pages 109–140. Academic Press, San Diego, CA, USA.

Hunt, G., Hopkins, M.J. and Lidgard, S. (2015). Simple versus complex models of trait evolution and stasis as a response to environmental change. Proc. Natl. Acad. Sci. USA, 112, 4885–4890.

Jablonka, E. and Lamb, M.J. (2014). Evolution in Four Dimensions. The MIT Press, London, England, revised edition.

Jones, A.G., Bürger, R. and Arnold, S.J. (2014a). Epistasis and natural selection shape the mutational architecture of complex traits. Nat. Comm., 5, 3709.

Jones, O.R., Scheuerlein, A., Salguero-Gómez, R., Camarda, C.G., Schaible, R., Casper, B.B. et al (2014b). Diversity of ageing across the tree of life. Nature, 505, 169–173.

Kingsolver, J.G. and Diamond, S.E. (2011). Phenotypic selection in natural populations: What limits directional selection? Am. Nat., 177, 346–357.

Kingsolver, J.G., Hoekstra, H.E., Hoekstra, J.M., Berrigan, D., Vignieri, S.N., Hill, C.E. et al (2001). The strength of phenotypic selection in natural populations. Am. Nat., 157, 245–261.

Kirkpatrick, M. (2009). Patterns of quantitative genetic variation in multiple dimensions. Genetica, 136, 271–284.

Kirkpatrick, M. and Lofsvold, D. (1992). Measuring selection and constraint in the evolution of growth. Evolution, 46, 954–971.

Klingenberg, C.P. (2005). Variation, chapter 11. Developmental constraints, modules, and evolvability, pages 219–247. Academic Press, Burlington, MA, USA.

Klingenberg, C.P. (2010). Evolution and development of shape: integrating quantitative approaches. Nat. Rev. Genet., 11, 623–635.

Laland, K., Uller, T., Feldman, M., Sterelny, K., Müller, G.B., Moczek, A. et al (2014). Does evolutionary theory need a rethink? Yes, urgently. Nature, 514, 161–164.

Laland, K.N., Uller, T., Feldman, M.W., Sterelny, K., Müller, G.B., Moczek, A. et al (2015). The extended evolutionary synthesis: its structure, assumptions and predictions. Proc. R. Soc. B, 282, 20151019.

Lande, R. (1976). Natural selection and random genetic drift in phenotypic evolution. Evolution, 30, 314–334.

Lande, R. (1979). Quantitative genetic analysis of multivariate evolution applied to brain: body size allometry. Evolution, 34, 402–416.

Lande, R. (1980). The genetic covariance between characters maintained by pleiotropic mutations. Genetics, 94, 203–215.

Lande, R. (1982). A quantitative genetic theory of life history evolution. Ecology, 63, 607–615.

Lande, R. (2019). Developmental integration and evolution of labile plasticity in a complex quantitative character in a multiperiodic environment. Proc. Natl. Acad. Sci. USA, 116, 11361–11369.

Lande, R. and Arnold, S.J. (1983). The measurement of selection on correlated characters. Evolution, 37, 1210–1226.

Lehtonen, J. (2020). Longevity and the drift barrier: bridging the gap between Medawar and Hamilton. Evol. Lett., 4, 382–393.

Lynch, M. and Walsh, B. (1998). Genetics and Analysis of Quantitative Traits. Sinauer, Sunderland, MA, USA.

Maynard Smith, J., Burian, R., Kauffman, S., Alberch, P., Campbell, J., Goodwin, B. et al (1985). Developmental constraints and evolution. Q. Rev. Biol.

Medawar, P.B. (1952). An unsolved problem of biology. H. K. Lewis, London, UK.

Merilä, J., Sheldon, B. and Kruuk, L. (2001). Explaining stasis: microevolutionary studies in natural populations. Genetica, 112(1), 199–222.

Metz, J.A.J. (2011). Thoughts on the geometry of meso-evolution: collecting mathematical elements for a postmodern synthesis. In F. A. C. C. Chalub and J. F. Rodrigues, editors, The Mathematics of Darwin’s Legacy, pages 193–231. Springer.

Metz, J.A.J., Geritz, S.A.H., Meszéna, G., Jacobs, F.J.A. and van Heerwaarden, J.S. (1996). Adaptive dynamics, a geometrical study of the consequences of nearly faithful reproduction. In S. J. v. . S. M. Verduyn Lunel, editor, Stochastic and spatial structures of dynamical systems, pages 183–231. Amsterdam, North Holland.

Metz, J.A.J., Staňková, K. and Johansson, J. (2016). The canonical equation of adaptive dynamics for life histories: from fitnessreturns to selection gradients and Pontryagin’s maximum principle. J. Math. Biol., 72(4), 1125–1152.

Miloco, L. and Salazar-Ciudad, I. (2022). Evolution of the G matrix under non-linear genotypephenotype maps. Am. Nat., 199, 420–435.

Moore, A.J., Brodie III, E.D. and Wolf, J.B. (1997). Interacting phenotypes and the evolutionary process: I. direct and indirect genetic effects of social interactions. Evolution, 51(5), 1352–1362.

Morrissey, M.B. (2014). Selection and evolution of causally covarying traits. Evolution, 68, 1748–1761.

Morrissey, M.B. (2015). Evolutionary quantitative genetics of non-linear developmental systems. Evolution, 69, 2050–2066.

Müller, G.B. (2007). Evo-Devo: extending the evolutionary synthesis. Nat. Rev. Genet., 8, 943–949.

Mullon, C. and Lehmann, L. (2018). Eco-evolutionary dynamics in metacommunities: Ecological inheritance, helping within species, and harming between species. Am. Nat., 192, 664–686.

Nelson, P. and Masel, J. (2017). Intercellular competition and the inevitability of multicellular aging. Proc. Natl. Acad. Sci. USA, 114, 12982–12987.

Odling-Smee, F.J., Laland, K.N. and Feldman, M.W. (1996). Niche construction. Am. Nat., 147, 641–648.

Parvinen, K., Heino, M. and Dieckmann, U. (2013). Function-valued adaptive dynamics and optimal control theory. J. Math. Biol., 67, 509–533.

Pigliucci, M. (2007). Do we need an extended evolutionary synthesis? Evolution, 61, 2743–2749.

Pigliucci, M. and Müller, G.B., editors (2010). Evolution: The Extended Synthesis. The MIT Press, Cambridge, MA, USA.

Pigliucci, M. and Schlichting, C.D. (1997). On the limits of quantitative genetics for the study of phenotypic evolution. Acta Biotheor., 45, 143–160.

Salazar-Ciudad, I. and Marín-Riera, M. (2013). Adaptive dynamics under development-based genotype-phenotype maps. Nature, 497, 361–364.

Schaffer, W.M. (1983). The application of optimal control theory to the general life history problem. Am. Nat., 121, 418–431.

Schlosser, G. and Wagner, G.P., editors (2004). Modularity in Development and Evolution. The Univ. of Chicago Press, Chicago, IL, USA.

Schluter, D. (1996). Adaptive radiation along genetic lines of least resistance. Evolution, 50, 1766–1774.

Schmidt, M. and Lipson, H. (2009). Distilling free-form natural laws from experimental data. Science, 324, 81–85.

Sydsæter, K., Hammond, P., Seierstad, A. and Strom, A. (2008). Further Mathematics for Economic Analysis. Prentice Hall, 2nd edition.

Turelli, M. (1988). Phenotypic evolution, constant covariances, and the maintenance of additive variance. Evolution, 42, 1342–1347.

van Tienderen, P.H. (1995). Life cycle trade-offs in matrix population models. Ecology, 76, 2482–2489.

Vaupel, J.W., Baudisch, A., Dölling, M., Roach, D.A. and Gampe, J. (2004). The case for negative senescence. Theor. Pop. Biol., 65, 339–351.

Waddington, C.H. (1957). The Strategy of the Genes. George Allen & Unwin, London, UK.

Waddington, C.H. (1959). Evolutionary adaptation. Perspect. Biol. Med., 2, 379–401.

Waddington, C.H. (1961). Genetic assimilation. Adv. Genet., 10, 257–293.

Wagner, A. (2005). Robustness and Evolvability in Living Systems. Princeton Univ. Press, Princeton, NJ, USA.

Wagner, G.P. (1984). On the eigenvalue distribution of genetic and phenotypic dispersion matrices: Evidence for a nonrandom organization of quantitative character variation. J. Math. Biol., 21, 77–95.

Wagner, G.P. (1989). Multivariate mutation-selection balance with constrained pleiotropic effects. Genetics, 122, 223–234.

Walsh, B. and Lynch, M. (2018). Evolution and Selection of Quantitative Traits. Oxford Univ. Press, Oxford, UK.

Watson, R.A., Wagner, G.P., Pavlicev, M., Weinreich, D.M. and Mills, R. (2013). The evolution of phenotypic correlations and “developmental memory”. Evolution, 68, 1124–1138.

West-Eberhard, M.J. (2003). Developmental Plasticity and Evolution. Oxford Univ. Press, Oxford, UK.

Williams, G.C. (1957). Natural selection, and the evolution of senescence. Evolution, 11, 398–411.

Wright, S. (1937). The distribution of gene frequencies in populations. Proc. Natl. Acad. Sci. USA, 23, 307–320.

