## Supplementary Information for "How development affects evolution"

#### Contents

**S1 Matrix calculus notation** **1**

**S2 Exploring adaptation without selection** **1**

**S3 Development enables negative senescence** **3**

#### S1 Matrix calculus notation

Following Caswell (2019), we use the following notation from matrix calculus. The Jacobian matrix of a vector $\mathbf{a} \in \mathbb{R}^{n\times 1}$ with respect to a vector $\mathbf{b} \in \mathbb{R}^{m\times 1}$ in its standard or transposed form is, respectively,

$$\frac{\partial \mathbf{a}}{\partial \mathbf{b}^\intercal} = \begin{pmatrix} \frac{\partial a_1}{\partial b_1} & \cdots & \frac{\partial a_1}{\partial b_m} \\ \vdots & \ddots & \vdots \\ \frac{\partial a_n}{\partial b_1} & \cdots & \frac{\partial a_n}{\partial b_m} \end{pmatrix} \in \mathbb{R}^{n\times m} \quad \text{or} \quad \frac{\partial \mathbf{a}^\intercal}{\partial \mathbf{b}} = \begin{pmatrix} \frac{\partial a_1}{\partial b_1} & \cdots & \frac{\partial a_n}{\partial b_1} \\ \vdots & \ddots & \vdots \\ \frac{\partial a_1}{\partial b_m} & \cdots & \frac{\partial a_n}{\partial b_m} \end{pmatrix} \in \mathbb{R}^{m\times n}. \tag{S1}$$

The transpose of $\partial\mathbf{a}/\partial\mathbf{b}^\intercal$ is $(\partial\mathbf{a}/\partial\mathbf{b}^\intercal)^\intercal = \partial\mathbf{a}^\intercal/\partial\mathbf{b}$. The analogous notation applies for total derivatives.

#### S2 Exploring adaptation without selection

A debated question is whether adaptation may arise from plasticity without selection (West-Eberhard, 2003; Laland *et al.*, 2015). This question may be addressed using Eq. (19), whereby the evolutionary dynamics of developed phenotypes consists of selection response and exogenous plastic response. Here we ask if exogenous plastic response alone can lead to adaptation in a best-case scenario for adaptive plasticity, where development is selective.

From Eq. (19) in the main text, we have that exogenous plastic response of the geno-phenotype is

$$\left(\frac{\mathrm{s}\mathbf{z}}{\mathrm{s}\boldsymbol{\epsilon}^\intercal}\frac{\partial\boldsymbol{\epsilon}}{\partial\tau}\right)\bigg|_{\mathbf{y}=\bar{\mathbf{y}}}.$$

More specifically, from Layer 7, Eq. 1a of González-Forero (2021), we have that the exogenous plastic response of the phenotype is

$$\left(\frac{\mathrm{s}\mathbf{x}}{\mathrm{s}\boldsymbol{\epsilon}^\intercal}\frac{\partial\boldsymbol{\epsilon}}{\partial\tau}\right)\bigg|_{\mathbf{y}=\bar{\mathbf{y}}}.$$

To make the exogenous plastic response as adaptive as possible, we seek to write the stabilized plasticity of the phenotype $\mathrm{s}\mathbf{x}/\mathrm{s}\boldsymbol{\epsilon}^\intercal|_{\mathbf{y}=\bar{\mathbf{y}}}$ in terms of selection gradients, by letting development be selective. Using Layer 5, Eqs. 2b and 1; Layer 4, Eqs. 5 and 4; and Layer 3, Eq. 4 of González-Forero (2021), we have that the exogenous plastic response of the phenotype is

$$\frac{\mathrm{s}\mathbf{x}}{\mathrm{s}\boldsymbol{\epsilon}^\intercal}\frac{\partial\boldsymbol{\epsilon}}{\partial\tau} = \frac{\mathrm{s}\mathbf{x}}{\mathrm{s}\bar{\mathbf{x}}^\intercal}\frac{\mathrm{d}\mathbf{x}}{\mathrm{d}\boldsymbol{\epsilon}^\intercal}\frac{\partial\boldsymbol{\epsilon}}{\partial\tau} = \left(\mathbf{I}-\frac{\mathrm{d}\mathbf{x}}{\mathrm{d}\bar{\mathbf{x}}^\intercal}\right)^{-1}\frac{\mathrm{d}\mathbf{x}}{\mathrm{d}\boldsymbol{\epsilon}^\intercal}\frac{\partial\boldsymbol{\epsilon}}{\partial\tau} = \left(\mathbf{I}-\frac{\mathrm{d}\mathbf{x}}{\mathrm{d}\mathbf{x}^\intercal}\frac{\delta\mathbf{x}}{\delta\bar{\mathbf{x}}^\intercal}\right)^{-1}\frac{\mathrm{d}\mathbf{x}}{\mathrm{d}\mathbf{x}^\intercal}\frac{\partial\mathbf{x}}{\partial\boldsymbol{\epsilon}^\intercal}\frac{\partial\boldsymbol{\epsilon}}{\partial\tau}.$$

The two rightmost matrices are direct plasticity and exogenous environmental change, which give the vector of direct exogenous plastic response:

$$\frac{\partial \mathbf{x}}{\partial \boldsymbol{\epsilon}^\top}\frac{\partial \boldsymbol{\epsilon}}{\partial \tau} = \left(\sum_{j=1}^{N_\mathrm{a}} \frac{\partial \mathbf{x}_a}{\partial \boldsymbol{\epsilon}_j^\top}\frac{\partial \boldsymbol{\epsilon}_j}{\partial \tau}\right) = \left(\frac{\partial \mathbf{x}_a}{\partial \boldsymbol{\epsilon}_{a-1}^\top}\frac{\partial \boldsymbol{\epsilon}_{a-1}}{\partial \tau}\right) = \left(\frac{\partial \mathbf{g}_{a-1}}{\partial \boldsymbol{\epsilon}_{a-1}^\top}\frac{\partial \boldsymbol{\epsilon}_{a-1}}{\partial \tau}\right) = \left(\sum_{k=1}^{N_\mathrm{e}} \frac{\partial g_{i,a-1}}{\partial \epsilon_{k,a-1}}\frac{\partial \epsilon_{k,a-1}}{\partial \tau}\right),$$

where the second equality follows from Layer 2, Eq. 2c of González-Forero (2021). The last expression is in terms of age-specific direct plasticity $\partial g_{i,a-1}/\partial \epsilon_{k,a-1}$, which we now make a function of the selection gradient of the corresponding phenotype. Thus, we let development be selective. Specifically, let

$$\frac{\partial g_{i,a-1}}{\partial \epsilon_{k,a-1}} = V_{ik,a,a-1}\frac{\partial w}{\partial x_{i,a-1}}, \tag{S2}$$

for $a \in \{2,\ldots,N_\mathrm{a}\}$ and some differentiable function $V_{ik,a,a-1}(\mathbf{m}_{a-1},\bar{\mathbf{z}})$, where the selection gradient $\partial w/\partial x_{i,a-1}$ is a function of $\mathbf{m}_{a-1}$ and $\bar{\mathbf{z}}$ from Eq. (4) of the main text. Consequently, our assumptions for the developmental constraint (1) of the main text are met where the developmental map $\mathbf{g}_a$ is a function of mutant traits at the same age. It would be convenient that $\partial g_{i,a-1}/\partial \epsilon_{k,a-1}$ were also a function of $\partial \epsilon_{k,a-1}/\partial \tau$ to cancel exogenous environmental change and keep only selection gradients, but this is not allowed because this violates our assumption in the developmental constraint (1) of the main text that $\mathbf{g}_a$ depends on the environment at that age but not on the rate of exogenous environmental change: if we were to let the developmental map depend on the rate of exogenous environmental change, there would be a third term in the equation of the evolutionary dynamics of the phenotype, a term that would depend on the exogenous environmental acceleration over evolutionary time.

Now, let us define the matrix

$$\mathbf{V}_{a+1,a} = \begin{pmatrix} V_{11,a+1,a} & \cdots & V_{1N_\mathrm{e},a+1,a} \\ \vdots & \ddots & \vdots \\ V_{N_\mathrm{p}1,a+1,a} & \cdots & V_{N_\mathrm{p}N_\mathrm{e},a+1,a} \end{pmatrix} \in \mathbb{R}^{N_\mathrm{p}\times N_\mathrm{e}},$$

for $a \in \{1,\ldots,N_\mathrm{a}-1\}$. Then, using Eq. (S2), we can write the matrix of age-specific direct plasticity for age $a \in \{2,\ldots,N_\mathrm{a}\}$ as

$$\begin{aligned}\frac{\partial \mathbf{x}_a}{\partial \boldsymbol{\epsilon}_{a-1}^\top} &= \begin{pmatrix} \frac{\partial w}{\partial x_{1,a-1}}V_{11,a+1,a} & \cdots & \frac{\partial w}{\partial x_{1,a-1}}V_{1N_\mathrm{e},a+1,a} \\ \vdots & \ddots & \vdots \\ \frac{\partial w}{\partial x_{N_\mathrm{p},a-1}}V_{N_\mathrm{p}1,a+1,a} & \cdots & \frac{\partial w}{\partial x_{N_\mathrm{p},a-1}}V_{N_\mathrm{p}N_\mathrm{e},a+1,a} \end{pmatrix} \\ &= \begin{pmatrix} \frac{\partial w}{\partial x_{1,a-1}} & 0 & \cdots & 0 \\ 0 & \frac{\partial w}{\partial x_{2,a-1}} & \cdots & 0 \\ \vdots & \vdots & \ddots & \vdots \\ 0 & 0 & \cdots & \frac{\partial w}{\partial x_{N_\mathrm{p},a-1}} \end{pmatrix} \begin{pmatrix} V_{11,a+1,a} & \cdots & V_{1N_\mathrm{e},a+1,a} \\ \vdots & \ddots & \vdots \\ V_{N_\mathrm{p}1,a+1,a} & \cdots & V_{N_\mathrm{p}N_\mathrm{e},a+1,a} \end{pmatrix} \\ &= \mathrm{diag}\left(\frac{\partial w}{\partial \mathbf{x}_{a-1}}\right)\mathbf{V}_{a,a-1},\end{aligned}$$

where $\mathrm{diag}(\mathbf{x})$ is the diagonal matrix with vector $\mathbf{x}$ in its main diagonal. Hence, we can write the matrix of direct plasticity as

$$\frac{\partial \mathbf{x}}{\partial \boldsymbol{\epsilon}^\top} = \mathbf{UV},$$

where we define a matrix involving selection gradients as

$$\mathbf{U} = \begin{pmatrix} \mathbf{0} & \mathbf{0} & \cdots & \mathbf{0} \\ \mathbf{0} & \mathrm{diag}\left(\frac{\partial w}{\partial \mathbf{x}_1}\right) & \cdots & \mathbf{0} \\ \vdots & \vdots & \ddots & \vdots \\ \mathbf{0} & \mathbf{0} & \cdots & \mathrm{diag}\left(\frac{\partial w}{\partial \mathbf{x}_{N_\mathrm{a}-1}}\right) \end{pmatrix} = \mathrm{diag}\left(\left(\mathbf{0}; \mathrm{diag}\left(\frac{\partial w}{\partial \mathbf{x}_1}\right); \cdots; \mathrm{diag}\left(\frac{\partial w}{\partial \mathbf{x}_{N_\mathrm{a}-1}}\right)\right)\right)$$

and another matrix with the unspecified functions as

$$\mathbf{V} = \begin{pmatrix} \mathbf{0} & \mathbf{0} & \cdots & \mathbf{0} & \mathbf{0} \\ \mathbf{V}_{21} & \mathbf{0} & \cdots & \mathbf{0} & \mathbf{0} \\ \mathbf{0} & \mathbf{V}_{32} & \cdots & \mathbf{0} & \mathbf{0} \\ \vdots & \vdots & \ddots & \vdots & \vdots \\ \mathbf{0} & \mathbf{0} & \cdots & \mathbf{V}_{N_\mathrm{a},N_\mathrm{a}-1} & \mathbf{0} \end{pmatrix}.$$

Then, with selective development of the form (S2), the vector of direct exogenous plastic response is of the form

$$\frac{\partial \mathbf{x}}{\partial \boldsymbol{\epsilon}^\intercal}\frac{\partial \boldsymbol{\epsilon}}{\partial \tau} = \mathbf{U}\mathbf{V}\frac{\partial \boldsymbol{\epsilon}}{\partial \tau} = \left(\sum_{j=1}^{N_\mathrm{a}} \frac{\partial \mathbf{x}_a}{\partial \boldsymbol{\epsilon}_j^\intercal}\frac{\partial \boldsymbol{\epsilon}_j}{\partial \tau}\right) = \left(\frac{\partial \mathbf{x}_a}{\partial \boldsymbol{\epsilon}_{a-1}^\intercal}\frac{\partial \boldsymbol{\epsilon}_{a-1}}{\partial \tau}\right) = \left(\mathrm{diag}\left(\frac{\partial w}{\partial \mathbf{x}_{a-1}}\right)\mathbf{V}_{a,a-1}\frac{\partial \boldsymbol{\epsilon}_{a-1}}{\partial \tau}\right).$$

Thus, the best-case scenario for adaptation via exogenous plastic response given by Eq. (S2) yields a matrix $\mathbf{U}$ that involves the selection gradient with a lag in developmental time (the $a$-th block entry of the vector of direct exogenous plastic response depends on a diagonal matrix of the direct selection gradients of the phenotype at age $a-1$). Consequently, if there is an optimum phenotypic value at each age and it changes substantially with age, selective development of the form (S2) may yield maladaptive exogenous plastic response. Moreover, adaptation via exogenous plastic response with selective development of the form (S2) is prevented if the rate of change in exogenous environmental change changes in sign over evolutionary time (i.e., if $\partial \epsilon_{ka}/\partial \tau$ changes sign).

### S3 Development enables negative senescence

Here we derive Eq. (20) of the main text. From Eq. (19) in the main text, we have that evolutionary change in the geno-phenotype due to natural selection is given by

$$\iota \mathbf{L}_{\mathbf{zy}} \left.\frac{\mathrm{d}w}{\mathrm{d}\mathbf{y}}\right|_{\mathbf{y}=\bar{\mathbf{y}}}, \tag{S3a}$$

where the mechanistic socio-genetic cross-covariance matrix between the geno-phenotype and genotype is

$$\mathbf{L}_{\mathbf{zy}} = \frac{\mathrm{s}\mathbf{z}}{\mathrm{s}\mathbf{y}^\intercal}\mathbf{H}_\mathbf{y} = \begin{pmatrix}\mathbf{L}_{\mathbf{xy}} \\ \mathbf{H}_\mathbf{y}\end{pmatrix} \tag{S3b}$$

and

$$\mathbf{H}_\mathbf{y} = \mathrm{cov}[\mathbf{y},\mathbf{y}] \tag{S3c}$$

is equivalently the mutational covariance matrix (of the genotype) and the mechanistic additive genetic covariance matrix of the genotype. The matrix $\mathrm{s}\mathbf{z}/\mathrm{s}\mathbf{y}^\intercal$ is a "stabilized" total derivative and it is non-singular because of our assumption that the genotype is developmentally independent (Appendix H of González-Forero, 2021). Hence, $\mathbf{L}_{\mathbf{zy}}$ is non-singular if $\mathbf{H}_\mathbf{y}$ is non-singular. Therefore, if $\iota \neq 0$ and $\mathbf{H}_\mathbf{y}$ is non-singular, then selection on the geno-phenotype vanishes if and only if $\mathrm{d}w/\mathrm{d}\mathbf{y} = \mathbf{0}|_{\mathbf{y}=\bar{\mathbf{y}}}$.

The total selection gradient of the genotype is

$$\left.\frac{\mathrm{d}w}{\mathrm{d}\mathbf{y}}\right|_{\mathbf{y}=\bar{\mathbf{y}}} = \left.\left(\frac{\delta w}{\delta \mathbf{y}} + \frac{\mathrm{d}\mathbf{x}^\intercal}{\mathrm{d}\mathbf{y}}\frac{\delta w}{\delta \mathbf{x}}\right)\right|_{\mathbf{y}=\bar{\mathbf{y}}} \tag{S4}$$

(Layer 4 Eq. 22 of González-Forero 2021).

Eq. (4) in the main text gives a mutant's relative fitness in terms of generation time, which is

$$T = \sum_{j=1}^{N_\mathrm{a}} j \ell_j^\circ f_j^\circ, \tag{S5a}$$

(Eq. 6 of González-Forero 2021; Charlesworth 1994, Eq. 1.47c; Bulmer 1994, Eq. 25, Ch. 25; Bienvenu and Legendre 2015, Eqs. 5 and 12). The superscript $\circ$ denotes evaluation at $\mathbf{y} = \bar{\mathbf{y}}$, so $f_j^\circ$ and $p_j^\circ$ are, respectively, the fertility and survival probability of a neutral mutant at age $j$. A mutant's relative fitness (Eq. 4) also depends on the forces of selection on fertility and survival, which are respectively

$$\phi_j = \ell_j^\circ \tag{S5b}$$

$$\pi_j = \frac{1}{p_j^\circ}\sum_{k=j+1}^{N_\mathrm{a}} \ell_k^\circ f_k^\circ, \tag{S5c}$$

(Eqs. 7 of González-Forero 2021; Hamilton 1966 and Caswell 1978, his Eqs. 11 and 12), where the survivorship of neutral mutants is $\ell_j^\circ = \prod_{i=1}^{j-1} p_i^\circ$.

From Eqs. (S3), if the genotypic traits are mutationally uncorrelated (i.e., $\mathbf{H}_\mathbf{y}$ is diagonal), the evolutionary change of the $i$-th genotypic trait at age $a$, $\bar{y}_{ia}$, due to selection is given by

$$\iota H_{y_{ia}} \left.\frac{\mathrm{d}w}{\mathrm{d}y_{ia}}\right|_{\mathbf{y}=\bar{\mathbf{y}}}. \tag{S6}$$

Since $H_{y_{ia}}$ is a variance, it is non-negative, so the $i$-th resident genotypic trait at age $a$ increases over evolutionary time if and only if the total selection gradient of this genotypic trait is positive, provided that there is mutational variation for this genotypic trait at that age (i.e., $\iota H_{y_{ia}} > 0$). From (S4), the total selection gradient of the genotypic trait $y_{ia}$ is

$$\left.\frac{\mathrm{d}w}{\mathrm{d}y_{ia}}\right|_{\mathbf{y}=\bar{\mathbf{y}}} = \left.\left(\frac{\delta w}{\delta y_{ia}} + \sum_{k=1}^{N_\mathrm{p}} \sum_{j=1}^{N_\mathrm{a}} \frac{\mathrm{d}x_{kj}}{\mathrm{d}y_{ia}} \frac{\delta w}{\delta x_{kj}}\right)\right|_{\mathbf{y}=\bar{\mathbf{y}}} = \left.\left(\frac{\delta w_a}{\delta y_{ia}} + \sum_{k=1}^{N_\mathrm{p}} \sum_{j=1}^{N_\mathrm{a}} \frac{\mathrm{d}x_{kj}}{\mathrm{d}y_{ia}} \frac{\delta w_j}{\delta x_{kj}}\right)\right|_{\mathbf{y}=\bar{\mathbf{y}}}, \tag{S7}$$

where the second equality follows because total immediate derivatives do not consider developmental constraints.

Let us now see that the forces of selection decrease, or remain constant, with age as has been long established (Hamilton, 1966; Wensink *et al.*, 2017; Caswell and Shyu, 2017). The change in the force on fertility with age is

$$\phi_{j+1} - \phi_j = \ell_{j+1}^\circ - \ell_j^\circ = \ell_j^\circ p_j^\circ - \ell_j^\circ = \ell_j^\circ (p_j^\circ - 1) \leq 0, \tag{S8a}$$

where the rightmost inequality follows because $p_j^\circ$ is a probability. Hence, the force on fertility is non-increasing with age. In turn, the change in the force on survival with age is

$$\begin{aligned} \pi_{j+1} - \pi_j &= \frac{1}{p_{j+1}^\circ} \sum_{k=j+2}^{N_\mathrm{a}} \ell_k^\circ f_k^\circ - \frac{1}{p_j^\circ} \sum_{k=j+1}^{N_\mathrm{a}} \ell_k^\circ f_k^\circ \\ &= \left(\frac{1}{p_{j+1}^\circ} - \frac{1}{p_j^\circ}\right) \sum_{k=j+2}^{N_\mathrm{a}} \ell_k^\circ f_k^\circ - \frac{1}{p_j^\circ} \ell_{j+1}^\circ f_{j+1}^\circ, \end{aligned} \tag{S8b}$$

which is non-positive if $p_j^\circ \to p_{j+1}^\circ$ as is the case if the survival probability changes smoothly with age.

Now, suppose that the genotypic trait $y_{ia}$ has a deleterious effect on survival or fertility at an early age $a$ such that $\delta w_a / \delta y_{ia} < 0$, and a pleiotropic, beneficial effect on survival or fertility of similar magnitude at a later age $j > a$ such that $\delta w_j / \delta x_{kj} > 0$ for some phenotype $x_{kj}$, but no other fitness effects. Then, total immediate selection is weaker at the later age because of declining selection forces (i.e., $|\delta w_a / \delta y_{ia}| \geq |\delta w_j / \delta x_{kj}|$). Yet, from (S7) we obtain that such genotypic trait is favoured if its total effect on the phenotype is sufficiently large, that is, if

$$\left.\left(\frac{\delta w_a}{\delta y_{ia}} + \frac{\mathrm{d}x_{kj}}{\mathrm{d}y_{ia}} \frac{\delta w_j}{\delta x_{kj}}\right)\right|_{\mathbf{y}=\bar{\mathbf{y}}} > 0.$$

In such case, from (S6), the resident genotype $\bar{y}_{ia}$ increases if its mutation rate and mutational variance are non-zero.

### References


Bienvenu, F. and Legendre, S. (2015). A new approach to the generation time in matrix population models. *Am. Nat.*, **185**, 834–843.

Bulmer, M. (1994). *Theoretical Evolutionary Ecology*. Sinauer, Sunderland, MA, USA.

Caswell, H. (1978). A general formula for the sensitivity of population growth rate to changes in life history parameters. *Theor. Popul. Biol.*, **14**, 215–230.

Caswell, H. (2019). *Sensitivity Analysis: Matrix Methods in Demography and Ecology*. Springer Open, Cham, Switzerland.

Caswell, H. and Shyu, E. (2017). *Senescence, selection gradients and mortality*, chapter 4, pages 56–82. Cambridge Univ. Press, Cambridge, UK.

Charlesworth, B. (1994). *Evolution in age-structured populations*. Cambridge Univ. Press, 2nd edition.

González-Forero, M. (2021). A mathematical framework for evo-devo dynamics. In review at *Theor. Popul. Biol.* Preprint: https://www.biorxiv.org/content/10.1101/2021.05.17.444499v3.

Hamilton, W.D. (1966). The moulding of senescence by natural selection. *J. Theor. Biol.*, **12**, 12–45.

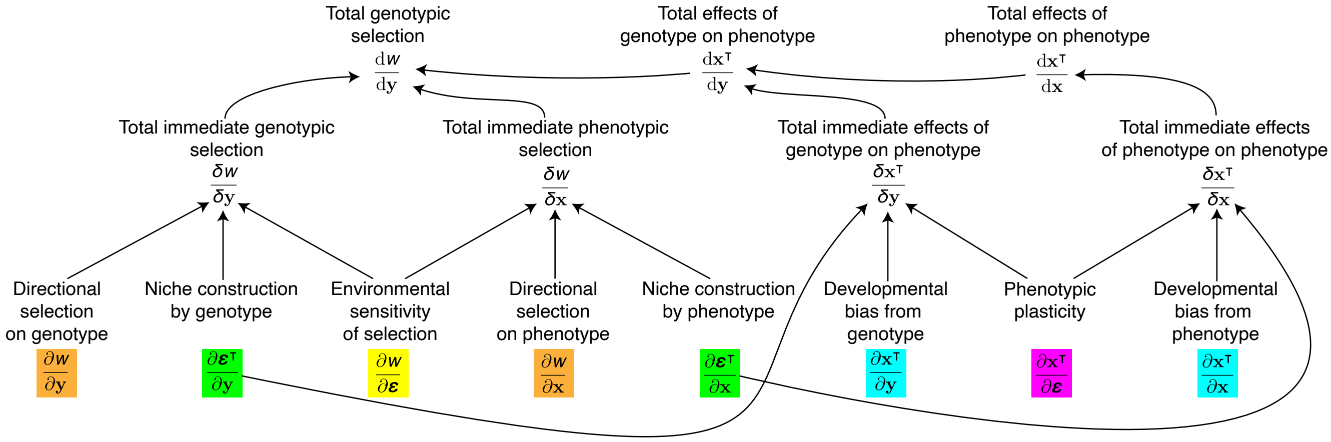


Figure S1: **Total genotypic selection depends on many factors.** Total genotypic selection — which is measured by the total selection gradient of the genotype $\mathrm{d}w/\mathrm{d}\mathbf{y}$ — describes selection response relatively completely in that it is premultiplied by a non-singular matrix $\mathbf{L}_{\mathbf{zy}}$ if there are no absolute mutational constraints, so total genotypic selection can identify evolutionary equilibria. In contrast, direct phenotypic and genotypic selection — measured by $\partial w/\partial \mathbf{z}$ — and total phenotypic and genotypic selection — measured by $\mathrm{d}w/\mathrm{d}\mathbf{z}$ — describe selection response less completely in that they are always premultiplied by a singular matrix and so do not generally identify evolutionary equilibria. Total genotypic selection depends on direct directional selection, developmental bias, plasticity, niche construction, and environmental sensitivity of selection. An arrow from a variable to another one indicates that the latter depends on the former.

Laland, K.N., Uller, T., Feldman, M.W., Sterelny, K., Müller, G.B., Moczek, A. et al (2015). The extended evolutionary synthesis: its structure, assumptions and predictions. *Proc. R. Soc. B*, **282**, 20151019.

Wensink, M.J., Caswell, H. and Baudisch, A. (2017). The rarity of survival to old age does not drive the evolution of senescence. *Evol. Biol.*, **44**(1), 5–10.

West-Eberhard, M.J. (2003). *Developmental Plasticity and Evolution.* Oxford Univ. Press, Oxford, UK.
