## Supplementary material for "How development affects evolution": Computer Code

### Computer code for:
### How development affects evolution

Mauricio González-Forero
School of Biology, University of St Andrews, Dyers Brae, St Andrews, KY16 9TH, Fife, UK

This file contains the computer code used to generate the figures in the main text. This code was prepared in Mathematica 12.1.1.0 and it is made available under a Creative Commons Attribution licence (CC BY).

#### Fig. 1. No social development, no niche construction, and no exogenous environmental change

Change developmental map by typing:
Case 1 for $g(y) = 1 + y$ (Fig. 1a,d,g,j),
Case 2 for $g(y) = 1 + y - \frac{3}{2}y^2$ (Fig. 1b,e,h,k), or
Case 3 for $g(y) = y^3 + \frac{5}{2}y + \frac{1}{5}$ (Fig. 1c,f,i,l)

```
In[1]:= Clear["Global`*"]

Case = 2;
(*Enter fitness and developmental map*)
w[x_, y_] := Exp[-(x^2 + y^2)]
g[y_] :=
 If[Case == 1, 1 + y, If[Case == 2, 1 + y - 3/2 y^2, If[Case == 3, y^3 - 5/2 y + 1/5]]]

(*Enter values for plot sizes*)
zLim = 2; (*Limits for the axes*)
gThickness = 0.02; (*Line thickness for g*)
eqThickness = 0.02; (*Line thickness for evolutionary equilibria*)
eqRadius = 0.2; (*Radius for disks indicating equilibria*)

(*The following points are the stable and unstable
  admissible equilibria which are found by solving for y in dwdy=0,
substituting this in g(y), and visually picking from the
 stream plot those points that are stable or unstable*)
StablePoints[case_] := If[case == 1, Disk[{-1/2, 1/2}, eqRadius],
  If[case == 2, {Disk[{1, 1/2}, eqRadius], Disk[{-Sqrt[2]/3, 2/3 - Sqrt[2]/3}, eqRadius]}, If[
    case == 3, {Disk[{-1.5550886683518514`, g[-1.5550886683518514`]}, eqRadius],
```

```
      Disk[{0.0690241712469999`, g[0.0690241712469999`]}, eqRadius],
      Disk[{1.4486053291218215`, g[1.4486053291218215`]}, eqRadius]}]]]
UnstablePoints[case_] := If[case == 2, Disk[{√2/3, 2/3 + √2/3}, eqRadius],
   If[case == 3, {Disk[{-1.016751601211979`, g[-1.016751601211979`]}, eqRadius],
     Disk[{1.0542107691950091`, g[1.0542107691950091`]}, eqRadius]}]]

(*The folloing are the stable points to add on the fitness landscape*)
StablePointsLandscape[case_] :=
 If[case == 1, Graphics3D[{Black, Ellipsoid[{StablePoints[Case][[1, 2]],
      StablePoints[Case][[1, 1]], w[StablePoints[Case][[1, 2]],
        StablePoints[Case][[1, 1]]]}, {eqRadius, eqRadius, eqRadius 3/4}]}],
   If[case == 2, {Graphics3D[{Black, Ellipsoid[
        {StablePoints[Case][[1, 1, 2]], StablePoints[Case][[1, 1, 1]],
         w[StablePoints[Case][[1, 1, 2]], StablePoints[Case][[1, 1, 1]]]},
        {eqRadius, eqRadius, eqRadius 3/4}]}], Graphics3D[{Black,
      Ellipsoid[{StablePoints[Case][[2, 1, 2]], StablePoints[Case][[2, 1, 1]],
         w[StablePoints[Case][[2, 1, 2]], StablePoints[Case][[2, 1, 1]]]},
        {eqRadius, eqRadius, eqRadius 3/4}]}]},
    If[case == 3, {Graphics3D[{Black, Ellipsoid[
         {StablePoints[Case][[1, 1, 2]], StablePoints[Case][[1, 1, 1]],
          w[StablePoints[Case][[1, 1, 2]], StablePoints[Case][[1, 1, 1]]]},
         {eqRadius, eqRadius, eqRadius 3/4}]}], Graphics3D[{Black,
       Ellipsoid[{StablePoints[Case][[2, 1, 2]], StablePoints[Case][[2, 1, 1]],
          w[StablePoints[Case][[2, 1, 2]], StablePoints[Case][[2, 1, 1]]]},
         {eqRadius, eqRadius, eqRadius 3/4}]}], Graphics3D[{Black,
       Ellipsoid[{StablePoints[Case][[3, 1, 2]], StablePoints[Case][[3, 1, 1]],
          w[StablePoints[Case][[3, 1, 2]], StablePoints[Case][[3, 1, 1]]]},
         {eqRadius, eqRadius, eqRadius 3/4}]}]}]]]

(*Plot fitness landscape and admissible path*)
Show[Plot3D[w[x, y], {x, -zLim, zLim}, {y, -zLim, zLim},
   AxesLabel → None, AxesStyle → Large, PlotStyle → Yellow],
 ParametricPlot3D[{g[y], y, w[g[y], y]}, {y, -zLim, zLim},
   PlotStyle → {Darker[Brown], Thickness[0.02]}],
 Sequence[StablePointsLandscape[Case]]]
Export[StringJoin[ToString[NotebookDirectory[]],
    StringJoin["Fig.1.Plain.Degen.1.FitnessLandscape.Direct.",
     ToString[Case]], ".pdf"], %];

(*Plot total fitness landscape of controls*)
Plot[w[g[y], y], {y, -zLim, zLim}, PlotRange → {0, 1},
 PlotStyle → {Darker[Brown], Thickness[0.02]}, AxesStyle → Large]
Export[StringJoin[ToString[NotebookDirectory[]],
    StringJoin["Fig.1.Plain.Degen.2.FitnessLandscape.Total.", ToString[Case]],
    ".pdf"], %];
```

```
(*Enter mutational covariance matrix.
  Calculate selection gradient, total selection gradient,
and total selection gradient of controls.
  Calculate phenotypic effects of controls and H matrix*)
Hy = {{0.01}};
partialwpartialz[x_, y_] = {{D[w[x, y], x]}, {D[w[x, y], y]}};
dwdz[x_, y_] = {{D[w[x, y], x]}, {D[w[g[y], y], y]}} /. {g[y] → x};
dwdy[x_, y_] = D[w[g[y], y], y] /. {g[y] → x};
dzdy[y_] = {{D[g[y], y], 1}};
Hz[y_] = Transpose[dzdy[y]].Hy.dzdy[y];

(*Plot H matrix*)
Plot[{Hz[y][[1, 1]], Hz[y][[1, 2]], Hz[y][[2, 2]]}, {y, -2, 2},
 AxesStyle → Large, PlotStyle → {{Lighter[Gray], Thickness[0.05]},
   {Cyan, Dashing[.2], Thickness[0.04]}, {Black, Dashing[.1], Thickness[0.02]}},
 PlotRange → {-.05, .1}, AspectRatio → 1]
Export[StringJoin[ToString[NotebookDirectory[]],
   StringJoin["Fig.1.Plain.Degen.3.H.", ToString[Case]], ".pdf"], %];

(*The following finds the maximum real part of the eigenvalues
 of the jacobian matrix of evolutionary dynamic system to identify
 stable and unstable equilibria: if λmax>0 at an equilibrium point,
the equililbrium is unstable;
if λmax<0 at an equilibrium point. To avoid numerical artifacts,
we use λmax>0.01 and λmax<0.01 to identify unstable and stable equilibria.*)
λmax[x_, y_] =
  Max[Re[Eigenvalues[Simplify[D[Flatten[Reverse[Hz[Y].partialwpartialz[X, Y]]],
       {{Y, X}}]]]]] /. {X → x, Y → y};

(*Color for stable and unstable equililbria*)
StableColor = {Blue};
UnstableColor = {Dashed, Blue};

(*Make stream plot*)
(*First, make a plot of evolutionary equilibria*)
dwdyPlot[case_] :=
 If[case == 1, Plot[-y, {y, -zLim, zLim}, Frame → True, FrameStyle → Large,
   AspectRatio → 1, PlotRange → {{-zLim, zLim}, {-zLim, zLim}},
   PlotStyle → {Blue, Thickness[gThickness]},
   Epilog → {StablePoints[case], {EdgeForm[Thick], FaceForm[Opacity[0]],
       UnstablePoints[case]}, {Magenta, Text[Style["★", 40], #] & /@ {{0, 0}}}}],
  If[case == 2, Show[Plot[y/(-1 + 3 y), {y, -zLim, zLim}, Frame → True,
     FrameStyle → Large, AspectRatio → 1, PlotRange →
      {{-zLim, zLim}, {-zLim, zLim}}, PlotStyle → {Blue, Thickness[gThickness]},
```

```
    RegionFunction → Function[{y}, λmax[y/(-1 + 3 y), y] < 0.0001],
    Epilog → {StablePoints[case], {EdgeForm[Thick], FaceForm[Opacity[0]],
       UnstablePoints[case]}, {Magenta, Text[Style["★", 40], #] & /@ {{0, 0}}}}],
   Plot[y/(-1 + 3 y), {y, -zLim, zLim}, Frame → True, FrameStyle → Large,
    AspectRatio → 1, PlotRange → {{-zLim, zLim}, {-zLim, zLim}},
    PlotStyle → {Blue, Dashing[.04], Thickness[gThickness]},
    RegionFunction → Function[{y}, λmax[y/(-1 + 3 y), y] > 0.0001]]], If[case == 3,
   Show[Plot[-(2 y)/(-5 + 6 y^2), {y, -zLim, zLim}, Frame → True, FrameStyle → Large,
     AspectRatio → 1, PlotRange → {{-zLim, zLim}, {-zLim, zLim}},
     PlotStyle → {Blue, Thickness[gThickness]}, RegionFunction →
      Function[{y}, λmax[-(2 y)/(-5 + 6 y^2), y] < 0.0001], Epilog → {StablePoints[case],
       {EdgeForm[Thick], FaceForm[Opacity[0]], UnstablePoints[case]},
       {Magenta, Text[Style["★", 40], #] & /@ {{0, 0}}}}],
    Plot[-(2 y)/(-5 + 6 y^2), {y, -zLim, zLim}, Frame → True, FrameStyle → Large,
     AspectRatio → 1, PlotRange → {{-zLim, zLim}, {-zLim, zLim}},
     PlotStyle → {Blue, Dashing[.04], Thickness[gThickness]},
     RegionFunction → Function[{y}, λmax[-(2 y)/(-5 + 6 y^2), y] > 0.0001]]]]]]
(*Then, superimpose the stream plot and the admissible path*)
Show[dwdyPlot[Case],
 StreamPlot[Flatten[Reverse[Hz[y].partialwpartialz[x, y]]], {y, -zLim, zLim},
  {x, -zLim, zLim}, StreamScale → {Full, 0.1, 0.03}, StreamStyle → Gray],
 Plot[g[y], {y, -zLim, zLim}, PlotRange → {{-zLim, zLim}, {-zLim, zLim}},
  PlotStyle → {Red, Thickness[gThickness]}]]
Export[StringJoin[ToString[NotebookDirectory[]],
   StringJoin["Fig.1.Plain.Degen.4.dzdt.", ToString[Case]], ".pdf"], %];
```

Out[12]=

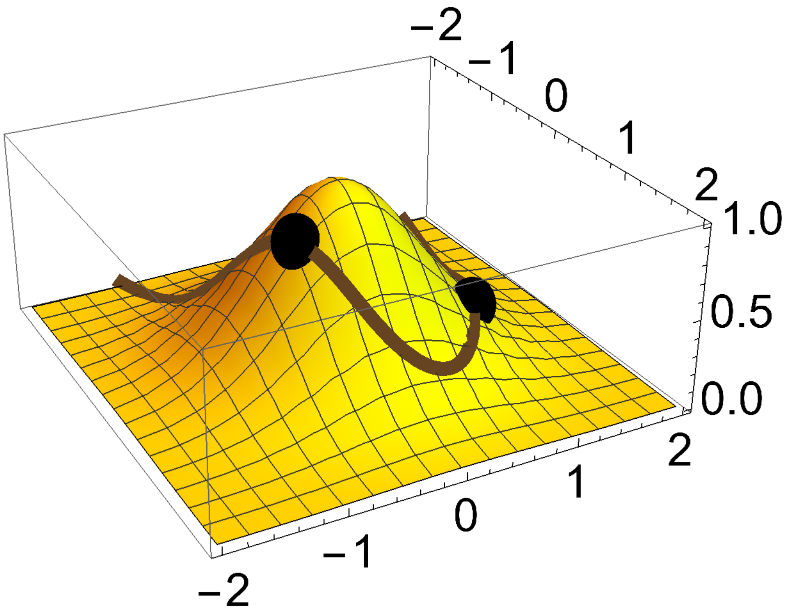


Out[14]=

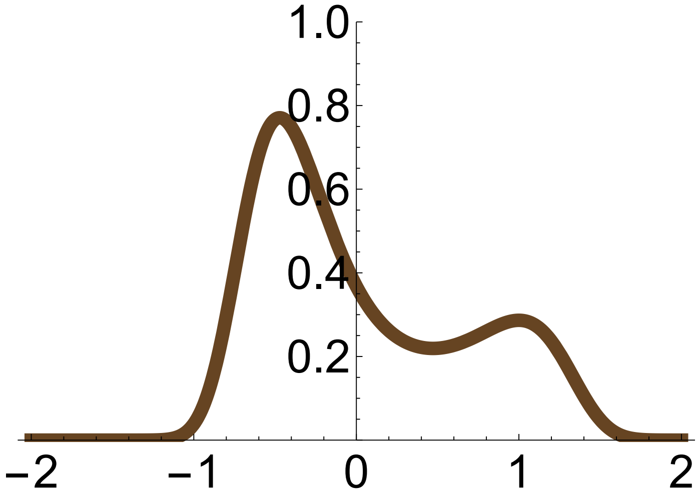

Out[22]=

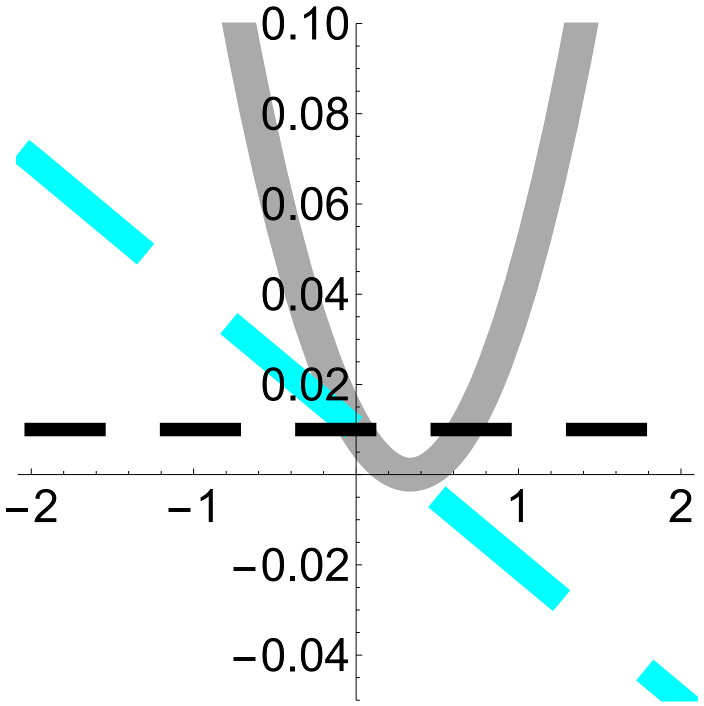


Out[28]=

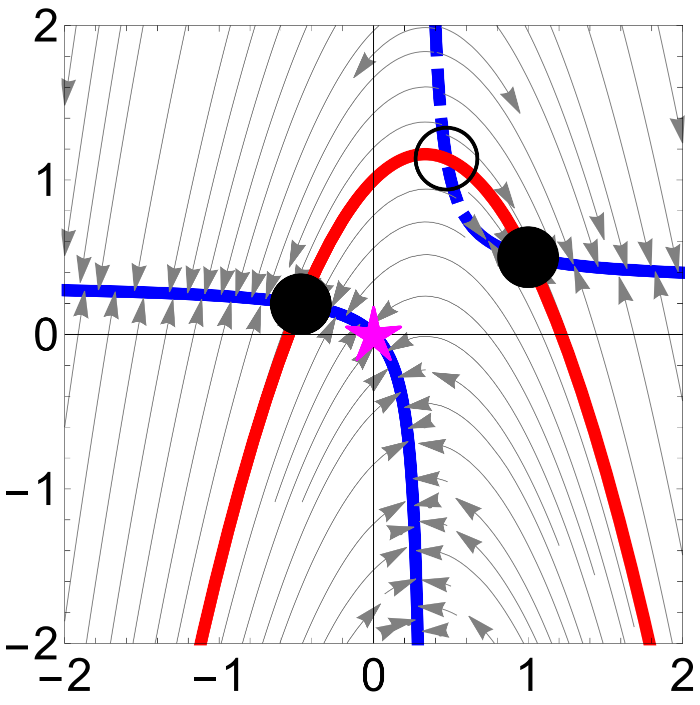


#### Fig. 2. No social development, no niche construction, and no exogenous environmental change

Change developmental map by typing:
Case 1 for $g(y) = 1 + y$ (Fig. 1a,d,g,j),
Case 2 for $g(y) = 1 + y - \frac{3}{2} y^2$ (Fig. 1b,e,h,k), or
Case 3 for $g(y) = y^3 + \frac{5}{2} y + \frac{1}{5}$ (Fig. 1c,f,i,l)

In[30]:=
```
Clear["Global`*"]
```

```
Case = 2;
Step = 1;
(*Enter fitness and developmental map*)
w[x_, y_] := Exp[-(x^2 + y^2)]
g[y_] :=
 If[Case == 1, Step + y, If[Case == 2, Step + y - 3/2 y^2, If[Case == 3, y^3 - 5/2 y + 1/5]]]

(*Enter values for plot sizes*)
zLim = 2; (*Limits for the axes*)
gThickness = 0.02; (*Line thickness for g*)
eqThickness = 0.02; (*Line thickness for evolutionary equilibria*)
eqRadius = 0.2; (*Radius for disks indicating equilibria*)

(*The following points are the stable and unstable
  admissible equilibria which are found by solving for y in dwdy=0,
substituing this in g(y), and visually picking from the
 stream plot those points that are stable or unstable*)
StablePoints[case_, step_] := If[case == 1 && step == 1,
  Disk[{-1/2, 1/2}, eqRadius], If[case == 1 && step == 0, Disk[{0, 0}, eqRadius],
   If[case == 1 && step == -1, Disk[{1/2, -1/2}, eqRadius], If[case == 2 && step == 1,
     {Disk[{1, 1/2}, eqRadius], Disk[{-Sqrt[2]/3, 2/3 - Sqrt[2]/3}, eqRadius]},
     If[case == 2 && step == 0, Disk[{0, g[0]}, eqRadius], If[case == 2 && step == -1,
       Disk[{Root[-2 + 10 * #1 - 9 * #1^2 + 9 * #1^3 & , 1, 0],
         g[Root[-2 + 10 * #1 - 9 * #1^2 + 9 * #1^3 & , 1, 0]]}, eqRadius], If[case ==
         3, {Disk[{-1.5550886683518514`, g[-1.5550886683518514`]}, eqRadius],
         Disk[{0.0690241712469999`, g[0.0690241712469999`]}, eqRadius],
         Disk[{1.4486053291218215`, g[1.4486053291218215`]}, eqRadius]}]]]]]]]
UnstablePoints[case_, step_] := If[case == 2 && step == 1, Disk[{Sqrt[2]/3, 2/3 + Sqrt[2]/3},
   eqRadius], If[case == 2 && (step == -1 || step == 0), Disk[{0, 10}, eqRadius],
   If[case == 3, {Disk[{-1.016751601211979`, g[-1.016751601211979`]}, eqRadius],
     Disk[{1.0542107691950091`, g[1.0542107691950091`]}, eqRadius]}]]]

(*The folloing are the stable points to add on the fitness landscape*)
StablePointsLandscape[case_, step_] := If[case == 1, Graphics3D[{Black, Ellipsoid[
     {StablePoints[case, step][[1, 2]], StablePoints[case, step][[1, 1]],
      w[StablePoints[case, step][[1, 2]], StablePoints[case, step][[1, 1]]]},
     {eqRadius, eqRadius, eqRadius 3/4}]}],
  If[case == 2 && step == 1, {Graphics3D[{Black, Ellipsoid[
        {StablePoints[case, step][[1, 1, 2]], StablePoints[case, step][[1, 1, 1]],
         w[StablePoints[case, step][[1, 1, 2]], StablePoints[case, step][[
           1, 1, 1]]]}, {eqRadius, eqRadius, eqRadius 3/4}]}],
    Graphics3D[{Black, Ellipsoid[{StablePoints[case, step][[2, 1, 2]],
        StablePoints[case, step][[2, 1, 1]],
```

```
        w[StablePoints[case, step][[2, 1, 2]], StablePoints[case, step][[
          2, 1, 1]]]}, {eqRadius, eqRadius, eqRadius 3/4}]}]},
   If[case == 2 && step == 0, {Graphics3D[{Black, Ellipsoid[
        {StablePoints[case, step][[1, 2]], StablePoints[case, step][[1, 1]],
         w[StablePoints[case, step][[1, 2]], StablePoints[case, step][[1, 1]]]},
        {eqRadius, eqRadius, eqRadius 3/4}]}]},
    If[case == 2 && step == -1, {Graphics3D[{Black, Ellipsoid[
         {StablePoints[case, step][[1, 2]], StablePoints[case, step][[1, 1]], w[
           StablePoints[case, step][[1, 2]], StablePoints[case, step][[1, 1]]]},
         {eqRadius, eqRadius, eqRadius 3/4}]}]},
     If[case == 3, {Graphics3D[{Black, Ellipsoid[{StablePoints[case, step][[
            1, 1, 2]], StablePoints[case, step][[1, 1, 1]],
           w[StablePoints[case, step][[1, 1, 2]], StablePoints[case, step][[
             1, 1, 1]]]}, {eqRadius, eqRadius, eqRadius 3/4}]}],
        Graphics3D[{Black, Ellipsoid[{StablePoints[case, step][[2, 1, 2]],
           StablePoints[case, step][[2, 1, 1]], w[StablePoints[case, step][[
             2, 1, 2]], StablePoints[case, step][[2, 1, 1]]]},
          {eqRadius, eqRadius, eqRadius 3/4}]}], Graphics3D[
        {Black, Ellipsoid[{StablePoints[case, step][[3, 1, 2]],
           StablePoints[case, step][[3, 1, 1]], w[StablePoints[case, step][[
             3, 1, 2]], StablePoints[case, step][[3, 1, 1]]]},
          {eqRadius, eqRadius, eqRadius 3/4}]}]}]]]]

(*Plot fitness landscape and admissible path*)
Show[Plot3D[w[x, y], {x, -2, 2}, {y, -2, 2},
  AxesLabel → None, AxesStyle → Large, PlotStyle → Yellow],
 ParametricPlot3D[{g[y], y, w[g[y], y]}, {y, -2, 2},
  PlotStyle → {Darker[Brown], Thickness[0.02]}],
 Sequence[StablePointsLandscape[Case, Step]]]
Export[StringJoin[ToString[NotebookDirectory[]],
   StringJoin["FigEx.2.Plain.Degen.1.FitnessLandscape.Direct.",
    ToString[Case], ".", ToString[Step]], ".pdf"], %];

(*Plot total fitness landscape of controls*)
Plot[w[g[y], y], {y, -2, 2}, PlotRange → {0, 1},
 PlotStyle → {Darker[Brown], Thickness[0.02]}, AxesStyle → Large]
Export[StringJoin[ToString[NotebookDirectory[]],
   StringJoin["FigEx.2.Plain.Degen.2.FitnessLandscape.Total.",
    ToString[Case], ".", ToString[Step]], ".pdf"], %];

(*Enter mutational covariance matrix.
  Calculate selection gradient, total selection gradient,
and total selection gradient of controls.
  Calculate phenotypic effects of controls and H matrix*)
Hy = {{0.01}};
partialwpartialz[x_, y_] = {{D[w[x, y], x]}, {D[w[x, y], y]}};
dwdz[x_, y_] = {{D[w[x, y], x]}, {D[w[g[y], y], y]}} /. {g[y] → x};
```

```
dwdy[x_, y_] = D[w[g[y], y], y] /. {g[y] → x};
dzdy[y_] = {{D[g[y], y], 1}};
Hz[y_] = Transpose[dzdy[y]].Hy.dzdy[y];

(*Plot H matrix*)
Plot[{Hz[y][[1, 1]], Hz[y][[1, 2]], Hz[y][[2, 2]]}, {y, -2, 2},
 AxesStyle → Large, PlotStyle → {{Lighter[Gray], Thickness[0.05]},
   {Cyan, Dashing[.2], Thickness[0.04]}, {Black, Dashing[.1], Thickness[0.02]}},
 PlotRange → {-.05, .1}, AspectRatio → 1]
Export[StringJoin[ToString[NotebookDirectory[]],
   StringJoin["FigEx.2.Plain.Degen.3.H.",
    ToString[Case], ".", ToString[Step]], ".pdf"], %];

(*The following finds the maximum real part of the eigenvalues
 of the jacobian matrix of evolutionary dynamic system to identify
 stable and unstable equilibria: if λmax>0 at an equilibrium point,
the equililbrium is unstable;
if λmax<0 at an equilibrium point. To avoid numerical artifacts,
we use λmax>0.01 and λmax<0.01 to identify unstable and stable equilibria.*)
λmax[x_, y_] =
  Max[Re[Eigenvalues[Simplify[D[Flatten[Reverse[Hz[Y].partialwpartialz[X, Y]]],
       {{Y, X}}]]]]]] /. {X → x, Y → y};

(*Color for stable and unstable equililbria*)
StableColor = {Blue};
UnstableColor = {Dashed, Blue};

(*Make stream plot*)
(*First, make a plot of evolutionary equilibria*)
dwdyPlot[case_, step_] :=
 If[case == 1, Plot[-y, {y, -zLim, zLim}, Frame → True, FrameStyle → Large,
   AspectRatio → 1, PlotRange → {{-zLim, zLim}, {-zLim, zLim}},
   PlotStyle → {Blue, Thickness[gThickness]}, Epilog → {StablePoints[case, step],
     {EdgeForm[Thick], FaceForm[Opacity[0]], UnstablePoints[case, step]},
     {Magenta, Text[Style["★", 40], #] & /@ {{0, 0}}}}], If[case == 2,
   Show[Plot[y/(-1 + 3 y), {y, -zLim, zLim}, Frame → True, FrameStyle → Large,
     AspectRatio → 1, PlotRange → {{-zLim, zLim}, {-zLim, zLim}},
     PlotStyle → {Blue, Thickness[gThickness]}, RegionFunction → Function[
       {y}, λmax[y/(-1 + 3 y), y] < 0.0001], Epilog → {StablePoints[case, step],
       {EdgeForm[Thick], FaceForm[Opacity[0]], UnstablePoints[case, step]},
       {Magenta, Text[Style["★", 40], #] & /@ {{0, 0}}}}],
    Plot[y/(-1 + 3 y), {y, -zLim, zLim}, Frame → True, FrameStyle → Large,
     AspectRatio → 1, PlotRange → {{-zLim, zLim}, {-zLim, zLim}},
     PlotStyle → {Blue, Dashing[.04], Thickness[gThickness]},
```

```
      RegionFunction → Function[{y}, λmax[y/(-1 + 3 y), y] > 0.0001]]], If[case == 3,
     Show[Plot[-(2 y)/(-5 + 6 y^2), {y, -zLim, zLim}, Frame → True, FrameStyle → Large,
       AspectRatio → 1, PlotRange → {{-zLim, zLim}, {-zLim, zLim}},
       PlotStyle → {Blue, Thickness[gThickness]}, RegionFunction → Function[
         {y}, λmax[-(2 y)/(-5 + 6 y^2), y] < 0.0001], Epilog → {StablePoints[case, step],
         {EdgeForm[Thick], FaceForm[Opacity[0]], UnstablePoints[case, step]},
         {Magenta, Text[Style["*", 40], #] & /@ {{0, 0}}}}],
      Plot[-(2 y)/(-5 + 6 y^2), {y, -zLim, zLim}, Frame → True, FrameStyle → Large,
       AspectRatio → 1, PlotRange → {{-zLim, zLim}, {-zLim, zLim}},
       PlotStyle → {Blue, Dashing[.04], Thickness[gThickness]},
       RegionFunction → Function[{y}, λmax[-(2 y)/(-5 + 6 y^2), y] > 0.0001]]]]]]
(*Then, superimpose the stream plot and the admissible path*)
Show[dwdyPlot[Case, Step],
 StreamPlot[Flatten[Reverse[Hz[y].partialwpartialz[x, y]]], {y, -zLim, zLim},
  {x, -zLim, zLim}, StreamScale → {Full, 0.1, 0.03}, StreamStyle → Gray],
 Plot[g[y], {y, -zLim, zLim}, PlotRange → {{-zLim, zLim}, {-zLim, zLim}},
  PlotStyle → {Red, Thickness[gThickness]}]]
Export[StringJoin[ToString[NotebookDirectory[]],
   StringJoin["FigEx.2.Plain.Degen.4.dzdt.",
    ToString[Case], ".", ToString[Step]], ".pdf"], %];
```

Out[42]=

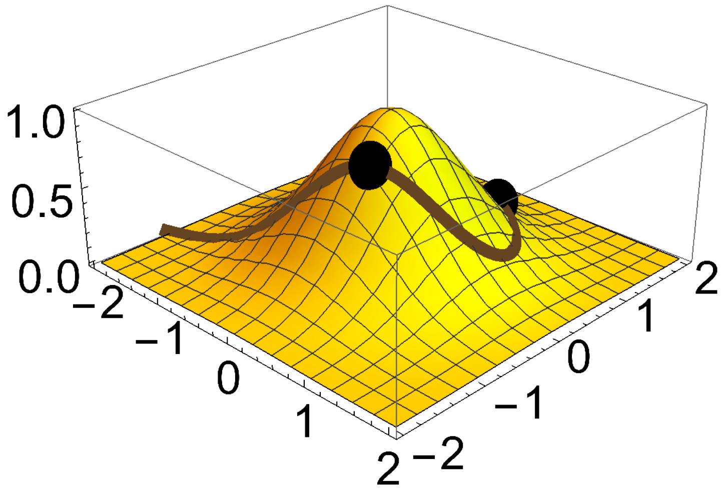

Out[44]=

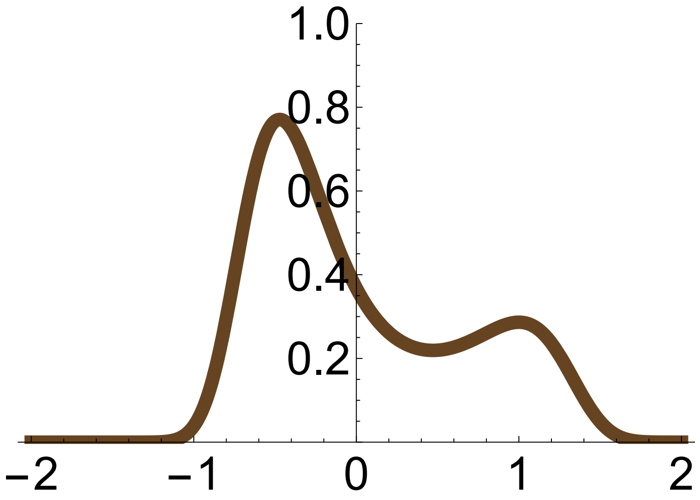


Out[52]=

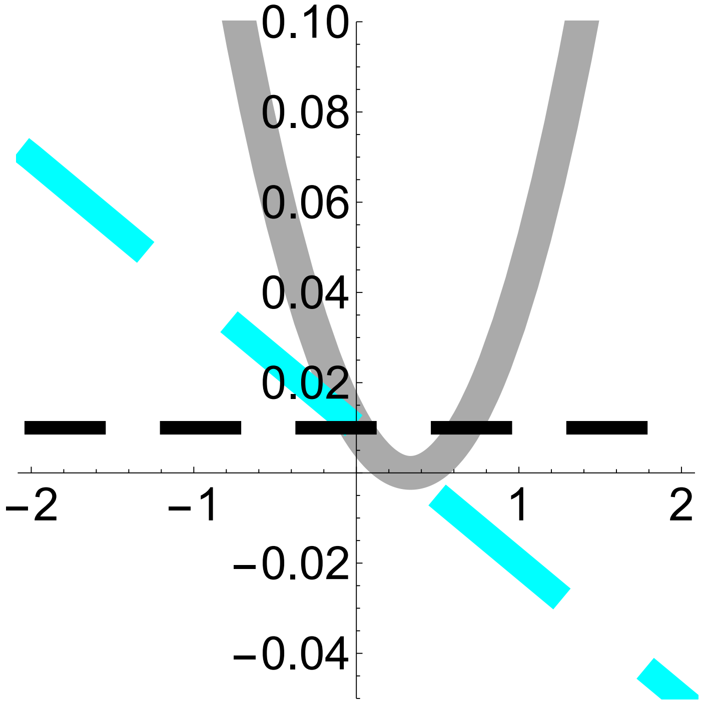

Out[58]=

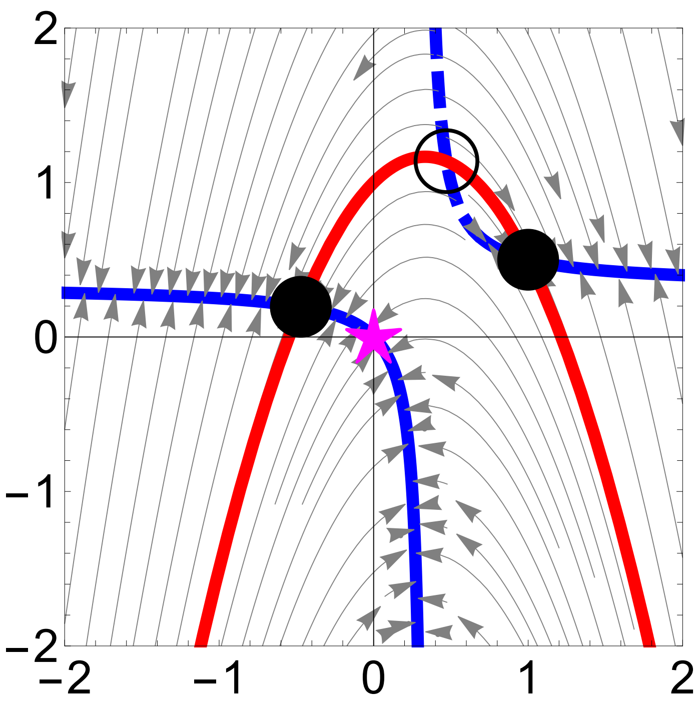


#### Fig. 3a-e. No social development, niche construction by state, environmental sensitivity of selection, and no exogenous environmental change

In[60]:=

```
Clear["Global`*"]

(*Enter fitness, developmental map, and environmental map*)
w[x_, y_, ϵ_] := ϵ Exp[- (x^2 + y^2)]
g[y_] := y
h[x_, y_] := 1 + 5 (x - .1)^2

(*Plot fitness landscape of the metaphenotype,
with a constant environment, and with the environmental constraint given*)
Show[DensityPlot3D[w[x, y, ϵ], {x, -2, 2}, {y, -2, 2},
   {ϵ, 0, 2}, ColorFunction → "Rainbow", AxesStyle → Large],
  Plot3D[1, {x, -2, 2}, {y, -2, 2}, AxesLabel → None,
   AxesStyle → Large, PlotStyle → Yellow],
  Plot3D[h[x, y], {x, -2, 2}, {y, -2, 2}, AxesLabel → None,
   AxesStyle → Large, PlotStyle → Darker[Green]]]
Export[StringJoin[ToString[NotebookDirectory[]],
    StringJoin["Fig.2.Niche.Degen.1.FitnessLandscape.Direct."], ".pdf"], %];

(*Plot fitness landscape of phenotype in constant environment*)
Show[Plot3D[w[x, y, 1], {x, -2, 2}, {y, -2, 2}, AxesLabel → None, AxesStyle → Large,
   PlotStyle → Yellow], ParametricPlot3D[{g[y], y, w[g[y], y, 1]},
   {y, -2, 2}, PlotStyle → {Darker[Brown], Thickness[0.02]}],
  Graphics3D[{Black, Ellipsoid[{0, 0, w[0, 0, h[0, 0]]}, {0.2, 0.2, 0.2 × 3 / 4}]}]]
Export[StringJoin[ToString[NotebookDirectory[]], StringJoin[
```

```
    "Fig.2.Niche.Degen.2.FitnessLandscape.Direct.ConstE."], ".pdf"], %];

(*Plot semi-
 total fitness landscape of phenotype given constant environmental constraint,
with the admissible path and a path under constant state*)
Show[Plot3D[w[x, y, h[x, y]], {x, -2, 2}, {y, -2, 2},
  AxesLabel → None, AxesStyle → Large, PlotStyle → Darker[Green]],
 ParametricPlot3D[{g[y], y, w[g[y], y, h[g[y], y]]}, {y, -2, 2},
  PlotStyle → {Darker[Brown], Thickness[0.02]}],
 Sequence[{Graphics3D[{Black, Ellipsoid[{-0.5282180222473377`,
       g[-0.5282180222473377`], w[g[-0.5282180222473377`],
        -0.5282180222473377`, h[g[-0.5282180222473377`],
         -0.5282180222473377`]]}, {0.2, 0.2, 0.2 × 3 / 4}]}],
   Graphics3D[{Black, Ellipsoid[{0.5588331454909822`, g[0.5588331454909822`],
       w[g[0.5588331454909822`], 0.5588331454909822`, h[g[0.5588331454909822`],
         0.5588331454909822`]]}, {0.2, 0.2, 0.2 × 3 / 4}]}]}]]
Export[StringJoin[ToString[NotebookDirectory[]], StringJoin[
    "Fig.2.Niche.Degen.3.FitnessLandscape.Semi."], ".pdf"], %];

(*Plot total fitness landscape of controls with a constant state*)
Plot[w[g[y], y, 1], {y, -2, 2},
 PlotStyle → {Darker[Brown], Thickness[0.02]}, AxesStyle → Large]
Export[StringJoin[ToString[NotebookDirectory[]], StringJoin[
    "Fig.2.Niche.Degen.4.FitnessLandscape.Total.ConstX."], ".pdf"], %];

(*Plot total fitness landscape of controls
 under the developmental constraint given*)
Plot[w[g[y], y, h[g[y], y]], {y, -2, 2},
 PlotStyle → {Darker[Brown], Thickness[0.02]}, AxesStyle → Large]
Export[StringJoin[ToString[NotebookDirectory[]],
   StringJoin["Fig.2.Niche.Degen.5.FitnessLandscape.Total."], ".pdf"], %];
```

Out[64]=

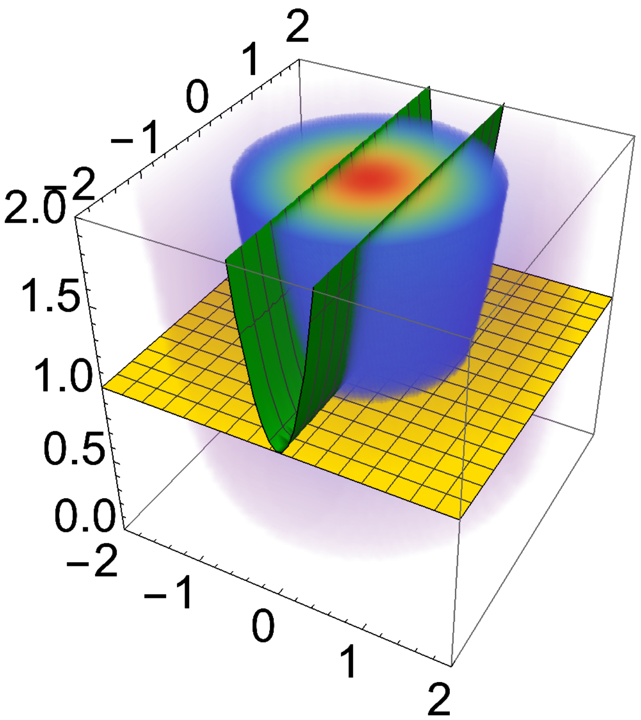


Out[66]=

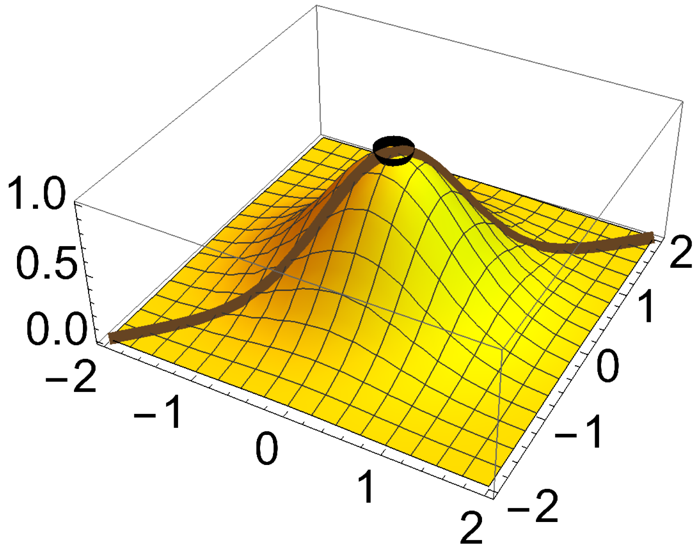

Out[68]=

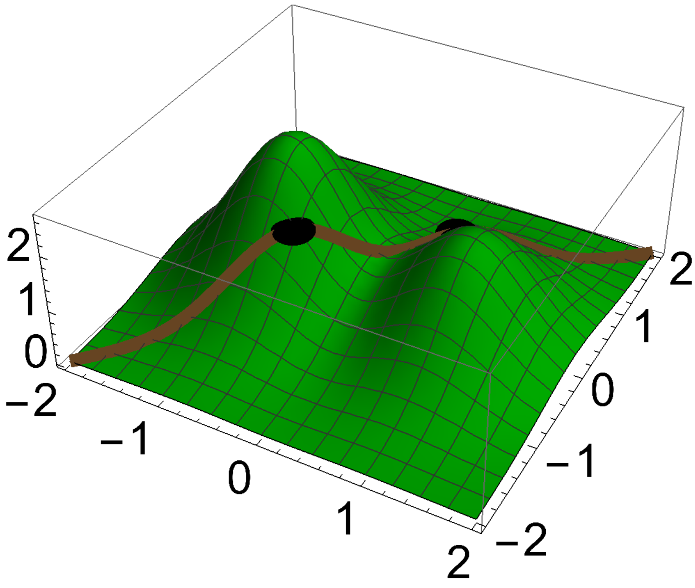


Out[70]=

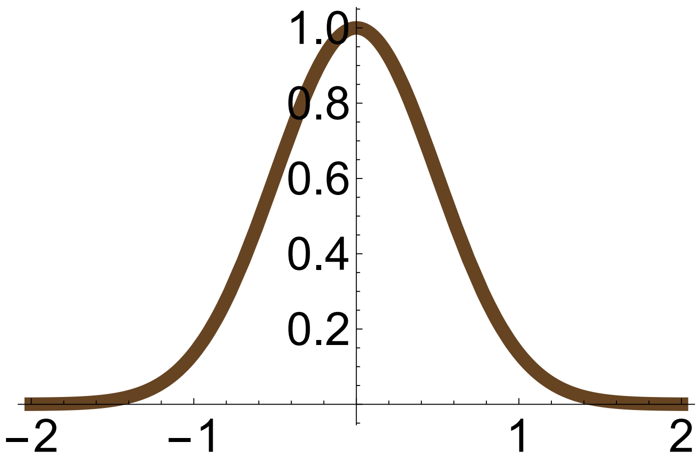


Out[72]=

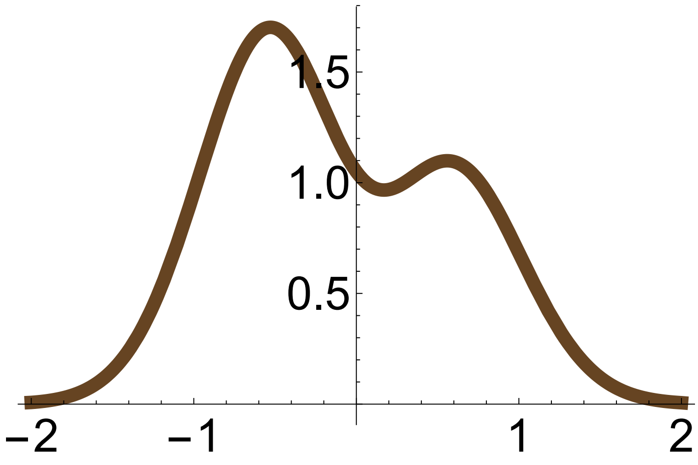


#### Fig. 3f-j. No social development, niche construction by control, plasticity, and no exogenous environmental change

In[74]:=
```
Clear["Global`*"]
```

```
(*Enter fitness, developmental map, and environmental map*)
```

```
w[x_, y_] := Exp[-(x^2 + y^2)]
g[y_, ϵ_] := 1 + ϵ y
h[y_] := 1 - 2 y

(*Plot fitness landscape of the metaphenotype,
with a constant environment, and with the environmental constraint given*)
Show[DensityPlot3D[w[x, y], {x, -2, 2}, {y, -2, 2},
   {ϵ, 0, 2}, ColorFunction → "Rainbow", AxesStyle → Large],
  Plot3D[1, {x, -2, 2}, {y, -2, 2}, AxesLabel → None,
   AxesStyle → Large, PlotStyle → Yellow],
  Plot3D[h[x, y], {x, -2, 2}, {y, -2, 2}, AxesLabel → None,
   AxesStyle → Large, PlotStyle → Darker[Green]]]
Export[StringJoin[ToString[NotebookDirectory[]],
    StringJoin["Fig.2f.Niche.Degen.1.FitnessLandscape.Direct."], ".pdf"], %];

(*Plot fitness landscape of phenotype in constant environment*)
Show[Plot3D[w[x, y], {x, -2, 2}, {y, -2, 2}, AxesLabel → None, AxesStyle → Large,
   PlotStyle → Yellow], ParametricPlot3D[{g[y, 1], y, w[g[y, 1], y]}, {y, -2, 2},
   PlotStyle → {Darker[Brown], Thickness[0.02]}], Graphics3D[{Black,
    Ellipsoid[{g[-0.5, 1], -0.5, w[g[-0.5, 1], -0.5]}, {0.2, 0.2, 0.2 × 3/4}]}]]
Export[StringJoin[ToString[NotebookDirectory[]], StringJoin[
     "Fig.2g.Niche.Degen.2.FitnessLandscape.Direct.ConstE."], ".pdf"], %];

(*Plot semi-
  total fitness landscape of phenotype given constant environmental constraint,
with the admissible path and a path under constant state*)
Show[Plot3D[w[x, y], {x, -2, 2}, {y, -2, 2},
   AxesLabel → None, AxesStyle → Large, PlotStyle → Darker[Green]],
  ParametricPlot3D[{g[y, h[y]], y, w[g[y, h[y]], y]}, {y, -2, 2},
   PlotStyle → {Darker[Brown], Thickness[0.02]}],
  Sequence[{Graphics3D[{Black, Ellipsoid[
        {g[-0.4446142795645972`, h[-0.4446142795645972`]], -0.4446142795645972`,
         w[g[-0.4446142795645972`, h[-0.4446142795645972`]],
          -0.4446142795645972`]}, {0.2, 0.2, 0.2 × 3/4}]}],
    Graphics3D[{Black, Ellipsoid[{g[0.8723221429525196`, h[0.8723221429525196`]],
         0.8723221429525196`, w[g[0.8723221429525196`, h[0.8723221429525196`]],
          0.8723221429525196`]}, {0.2, 0.2, 0.2 × 3/4}]}]}]]
Export[StringJoin[ToString[NotebookDirectory[]],
    StringJoin["Fig.2h.Niche.Degen.3.FitnessLandscape.Semi."], ".pdf"], %];

(*Plot total fitness landscape of controls with a constant environment*)
Plot[w[g[y, 1], y], {y, -2, 2},
  PlotStyle → {Darker[Brown], Thickness[0.02]}, AxesStyle → Large]
Export[StringJoin[ToString[NotebookDirectory[]], StringJoin[
     "Fig.2i.Niche.Degen.4.FitnessLandscape.Total.ConstE."], ".pdf"], %];

(*Plot total fitness landscape of controls
```

```
 under the developmental constraint given*)
Plot[w[g[y, h[y]], y], {y, -2, 2},
 PlotStyle → {Darker[Brown], Thickness[0.02]}, AxesStyle → Large]
Export[StringJoin[ToString[NotebookDirectory[]],
   StringJoin["Fig.2j.Niche.Degen.5.FitnessLandscape.Total."], ".pdf"], %];
```

Out[78]=

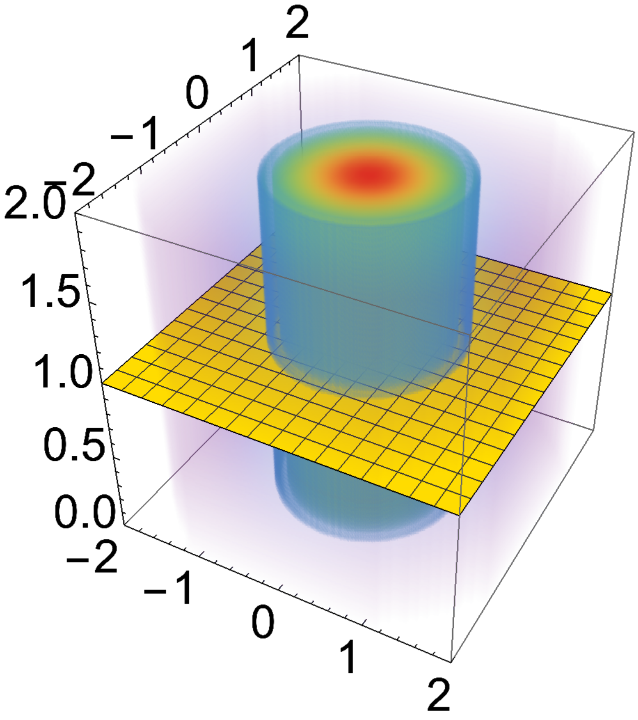


Out[80]=

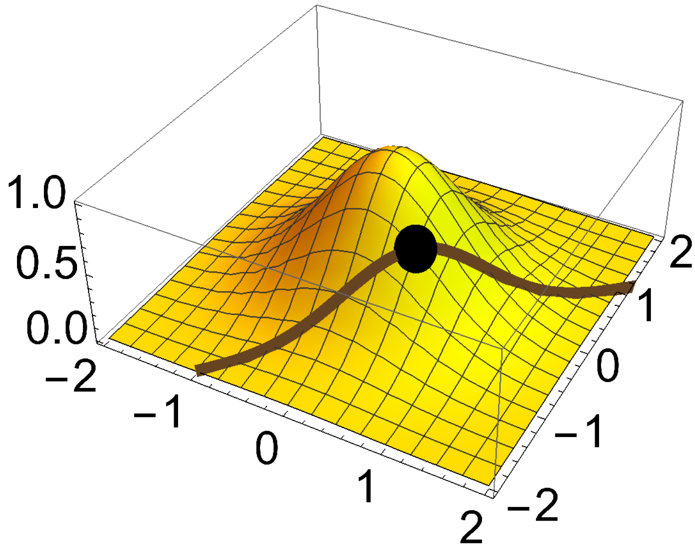

Out[82]=

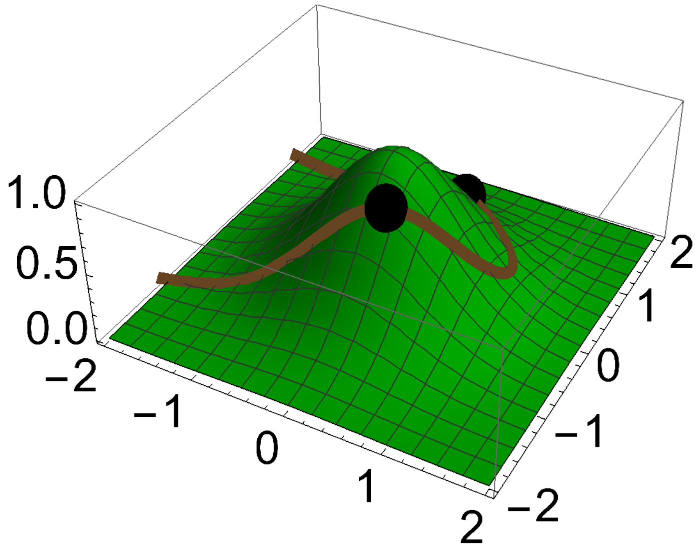


Out[84]=

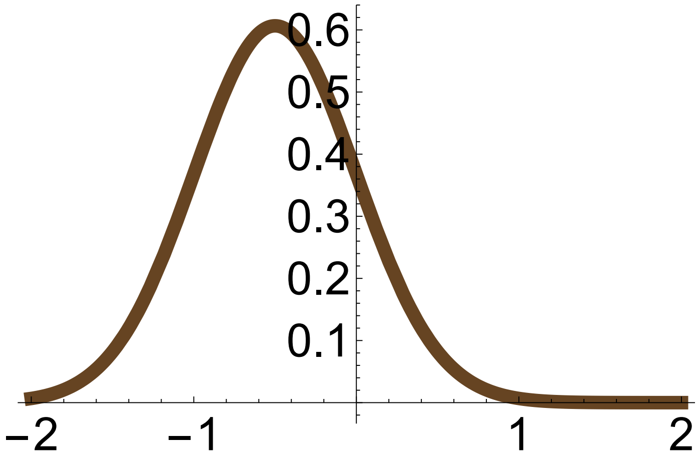


Out[86]=

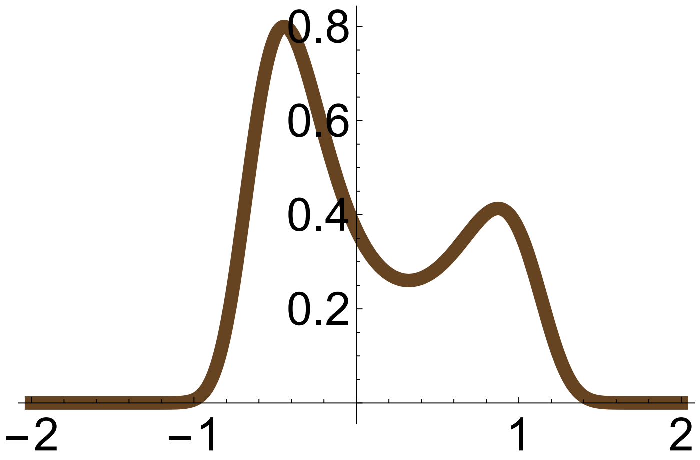


#### Fig. 4a,b. Social development, social developmental bias from partners' controls, no niche construction, and no exogenous environmental change

In[88]:=
```
Clear["Global`*"]

(*Enter the case for developmental map used (see code for Fig. 2)*)
Case = 1;
```

```
(*Enter fitness and developmental map*)
w[x_, y_] := Exp[- (x² + y²)]
g[y_, yb_] := 1 + y 1/2 yb

(*Enter mutational covariance matrix.
  Calculate selection gradient, total selection gradient,
and total selection gradient of controls.*)
Hy = {{0.01}};
partialwpartialz[x_, y_] = {{D[w[x, y], x]}, {D[w[x, y], y]}};
dwdz[x_, y_] =
  {{D[w[x, y], x]}, {D[w[g[y, yb], y], y]}} /. {yb → y} /. {g[y, y] → x};
dwdy[x_, y_] = D[w[g[y, yb], y], y] /. {yb → y} /. {g[y, y] → x};

(*Calculate phenotypic effects of controls.
  Calculate stabilized phenotypic effects of controls.
  Calculate L matrix*)
dzdy[y_] = {{D[g[y, yb], y], 1}} /. {yb → y};
szsy[y_] = {{D[g[y, yb], y] + D[g[y, yb], yb], 1}} /. {yb → y};
Lz[y_] = Transpose[szsy[y]].Hy.dzdy[y];

(*Enter values for plot sizes*)
zLim = 2; (*Limits for the axes*)
gThickness = 0.02; (*Line thickness for g*)
eqThickness = 0.02; (*Line thickness for evolutionary equilibria*)
eqRadius = 0.2; (*Radius for disks indicating equilibria*)

(*Plot L matrix*)
offset = 0;
Plot[{Lz[y][[1, 1]], Lz[y][[1, 2]], Lz[y][[2, 1]], Lz[y][[2, 2]]},
 {y, -zLim - offset, zLim + offset}, AxesStyle → Large,
 PlotStyle → {{Lighter[Gray], Thickness[0.05]},
   {Cyan, Dashing[.2], Thickness[0.04]}, {Orange, Dashing[.2], Thickness[0.04]},
   {Black, Dashing[.1], Thickness[0.02]}},
 PlotRange → {-.05, .1}, AspectRatio → 1]
Export[StringJoin[ToString[NotebookDirectory[]],
   StringJoin["Fig.3.Social.Degen.3.L."], ".pdf"], %];

(*The following points are the stable and unstable
  admissible equilibria which are found by solving for y in dwdy=0,
substituing this in g(y), and visually picking from the
 stream plot those points that are stable or unstable*)
StablePoints[case_] := {Disk[{0, 1}, eqRadius]}
UnstablePoints[case_] := Disk[{10, 1}, eqRadius]

(*The following finds the maximum real part of the eigenvalues
```

```
 of the jacobian matrix of evolutionary dynamic system to identify
 stable and unstable equilibria: if λmax>0 at an equilibrium point,
the equililbrium is unstable;
if λmax<0 at an equilibrium point. To avoid numerical artifacts,
we use λmax>0.01 and λmax<0.01 to identify unstable and stable equilibria.*)
λmax[x_, y_] =
  Max[Re[Eigenvalues[Simplify[D[Flatten[Reverse[Lz[Y].partialwpartialz[X, Y]]],
        {{Y, X}}]]]]] /. {X → x, Y → y};

(*Color for stable and unstable equililbria*)
StableColor = {Blue};
UnstableColor = {Dashed, Blue};

(*Make stream plot*)
(*First, make a plot of evolutionary equilibria*)
dwdyPlot[case_] :=
 Show[Plot[-2, {y, -zLim - offset, zLim + offset},
   Frame → True, FrameStyle → Large, AspectRatio → 1,
   PlotRange → {{-zLim - offset, zLim + offset}, {-zLim - offset, zLim + offset}},
   PlotStyle → {Blue, Thickness[gThickness]},
   Epilog → {{Directive[Blue, Thickness[gThickness]],
      Line[{{0, -2 - offset}, {0, 2 + offset}}]}, StablePoints[case],
     {EdgeForm[Thick], FaceForm[Opacity[0]], UnstablePoints[case]},
     {Magenta, Text[Style["★", 40], #] & /@ {{0, 0}}}}]]
(*Then, superimpose the stream plot and the admissible path*)
Show[dwdyPlot[Case], StreamPlot[Flatten[Reverse[Lz[y].partialwpartialz[x, y]]],
  {y, -zLim - offset, zLim + offset}, {x, -zLim - offset, zLim + offset},
  StreamScale → {Full, 0.1, 0.03}, StreamStyle → Gray],
 Plot[g[y, y], {y, -zLim - offset, zLim + offset},
  PlotRange → {{-zLim - offset, zLim + offset}, {-zLim - offset, zLim + offset}},
  PlotStyle → {Red, Thickness[gThickness]}]]
Export[StringJoin[ToString[NotebookDirectory[]],
   StringJoin["Fig.3.Social.Degen.4.dzdt."], ".pdf"], %];
```

Out[104]=

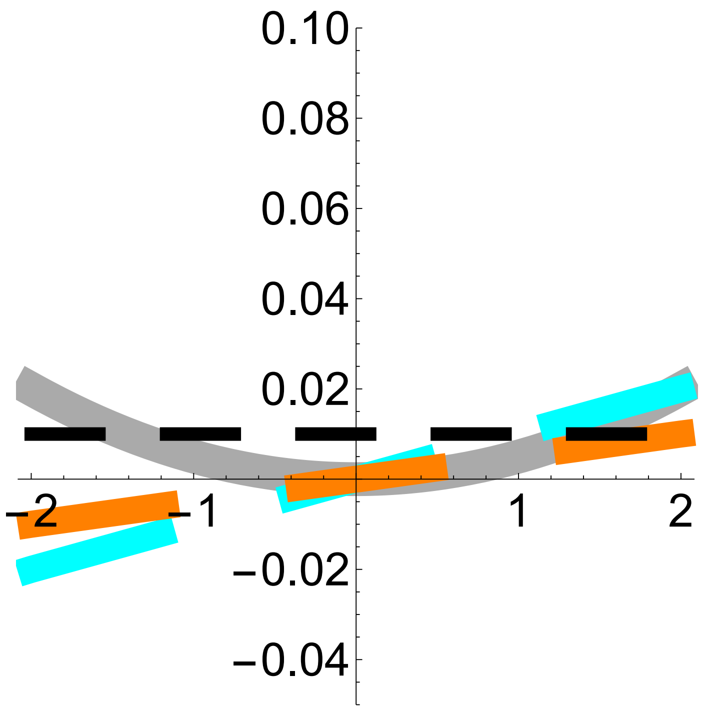


Out[112]=

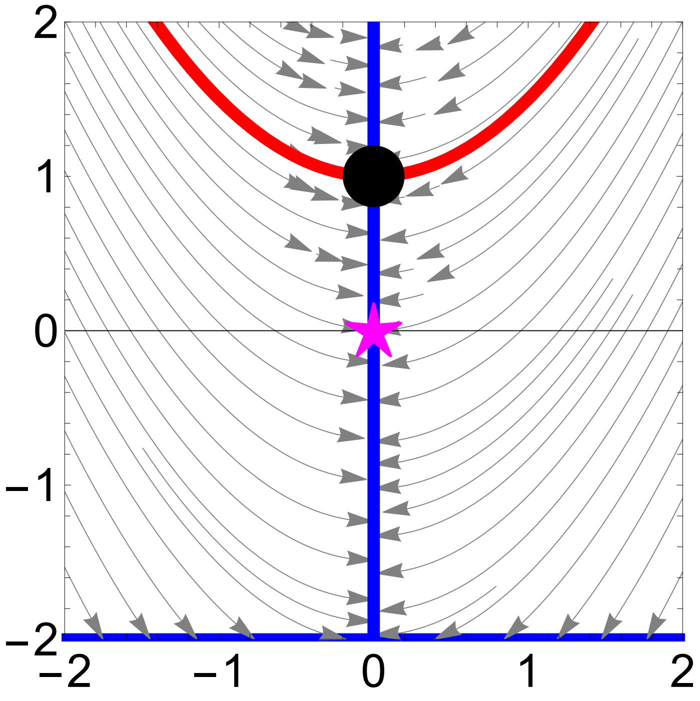


#### Fig. 4c,d. Social development, social developmental bias from partners' states, no niche construction, and no exogenous environmental change

In[114]:=
```
Clear["Global`*"]

(*Enter the case for developmental map used (see code for Fig. 2)*)
Case = 1;

(*Enter fitness and developmental map*)
```

```
w[x_, y_] := Exp[-(x^2 + y^2)]
g[y_, xb_] := 1 + y 1/2 xb

(*The following is the socio-
  devo equilibrium found by solving for x in x=g[y,x]*)
xs[y_] := - 2/(-2 + y)

(*Enter mutational covariance matrix.
  Calculate selection gradient, total selection gradient,
and total selection gradient of controls.*)
Hy = {{0.01}};
partialwpartialz[x_, y_] = {{D[w[x, y], x]}, {D[w[x, y], y]}};
dwdy[x_, y_] = D[w[g[y, xb], y], y] /. {g[y, xb] → x} /. {xb → x};

(*Calculate phenotypic effects of controls.
  Calculate stabilized phenotypic effects of controls.
  Calculate L matrix*)
dzdy[y_] = {{D[g[y, xb], y], 1}} /. {xb → xs[y]};
szsy[y_] = {{(1 - D[g[y, xb], xb])^-1 D[g[y, xb], y], 1}} /. {xb → xs[y]};
Lz[y_] = Transpose[szsy[y]].Hy.dzdy[y];

(*Plot L matrix*)
Plot[{Lz[y][[1, 1]], Lz[y][[1, 2]], Lz[y][[2, 1]], Lz[y][[2, 2]]}, {y, -2, 2},
 AxesStyle → Large, PlotStyle → {{Lighter[Gray], Thickness[0.05]},
   {Cyan, Dashing[.2], Thickness[0.04]}, {Orange, Dashing[.2], Thickness[0.04]},
   {Black, Dashing[.1], Thickness[0.02]}},
 PlotRange → {-.05, .1}, AspectRatio → 1]
Export[StringJoin[ToString[NotebookDirectory[]],
   StringJoin["Fig.3d.Social.Degen.3.L."], ".pdf"], %];

(*Enter values for plot sizes*)
zLim = 2; (*Limits for the axes*)
gThickness = 0.02; (*Line thickness for g*)
eqThickness = 0.02; (*Line thickness for evolutionary equilibria*)
eqRadius = 0.2; (*Radius for disks indicating equilibria*)

(*The following point is the stable admissible equilibria which
  is found by solving for y in dwdy=0, substituing this in g(y,xs),
and visually picking from the stream plot those points that are stable*)
StablePoints[case_] := {Disk[{Root[2 + 4 * #1 - 4 * #1^2 + #1^3 & , 1, 0],
    g[Root[2 + 4 * #1 - 4 * #1^2 + #1^3 & , 1, 0],
     xs[Root[2 + 4 * #1 - 4 * #1^2 + #1^3 & , 1, 0]]]}, eqRadius]}

(*Color for stable and unstable equililbria*)
StableColor = {Blue};
```

```
UnstableColor = {Dashed, Blue};

(*The following finds the maximum real part of the eigenvalues
 of the jacobian matrix of evolutionary dynamic system to identify
 stable and unstable equilibria: if λmax>0 at an equilibrium point,
the equilibrium is unstable;
if λmax<0 at an equilibrium point. To avoid numerical artifacts,
we use λmax>0.01 and λmax<0.01 to identify unstable and stable equilibria.*)
λmax[x_, y_] =
  Max[Re[Eigenvalues[Simplify[D[Flatten[Reverse[Lz[Y].partialwpartialz[X, Y]]],
        {{Y, X}}]]]]]] /. {X → x, Y → y};

(*Make stream plot*)
(*First, make a plot of evolutionary equilibria*)
dwdyPlot[case_] :=
 Show[Plot[(-1 + Sqrt[1 - 4 y^2])/y, {y, -zLim, zLim}, Frame → True, FrameStyle → Large,
   AspectRatio → 1, PlotRange → {{-zLim, zLim}, {-zLim, zLim}},
   PlotStyle → {Blue, Thickness[gThickness]},
   RegionFunction → Function[{y}, λmax[(-1 + Sqrt[1 - 4 y^2])/y, y] < 0.0001], Epilog →
    {StablePoints[case], {Magenta, Text[Style["★", 40], #] & /@ {{0, 0}}}}],
  Plot[(-1 + Sqrt[1 - 4 y^2])/y, {y, -zLim, zLim}, Frame → True, FrameStyle → Large,
   AspectRatio → 1, PlotRange → {{-zLim, zLim}, {-zLim, zLim}},
   PlotStyle → {Blue, Dashing[.04], Thickness[gThickness]},
   RegionFunction → Function[{y}, λmax[(-1 + Sqrt[1 - 4 y^2])/y, y] > 0.0001]]]
(*Then, superimpose the stream plot and the admissible path*)
Show[dwdyPlot[Case],
 StreamPlot[Flatten[Reverse[Lz[y].partialwpartialz[x, y]]], {y, -zLim, zLim},
  {x, -zLim, zLim}, StreamScale → {Full, 0.1, 0.03}, StreamStyle → Gray],
 Plot[g[y, xs[y]], {y, -zLim, zLim}, PlotRange → {{-zLim, zLim}, {-zLim, zLim}},
  PlotStyle → {Red, Thickness[gThickness]}]]
Export[StringJoin[ToString[NotebookDirectory[]],
   StringJoin["Fig.3c.Social.Degen.4.dzdt."], ".pdf"], %];
```

Out[125]=

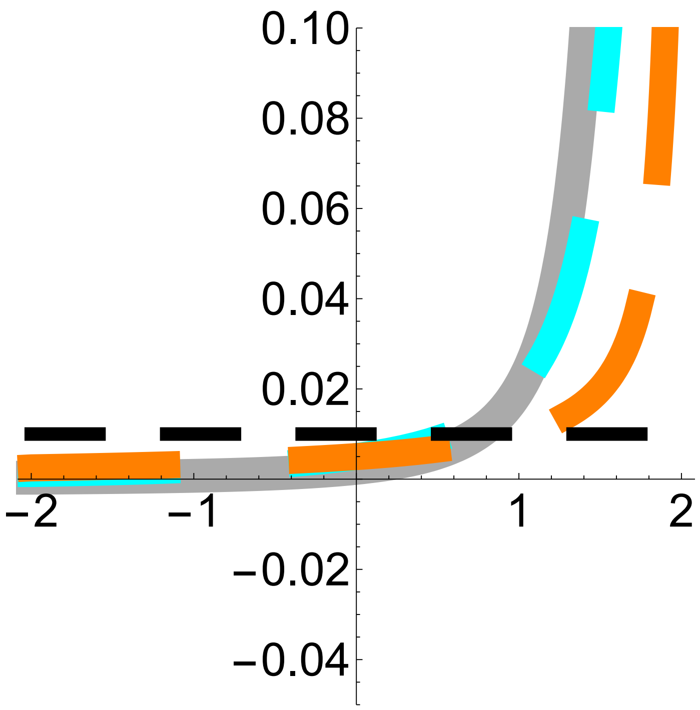


Out[136]=

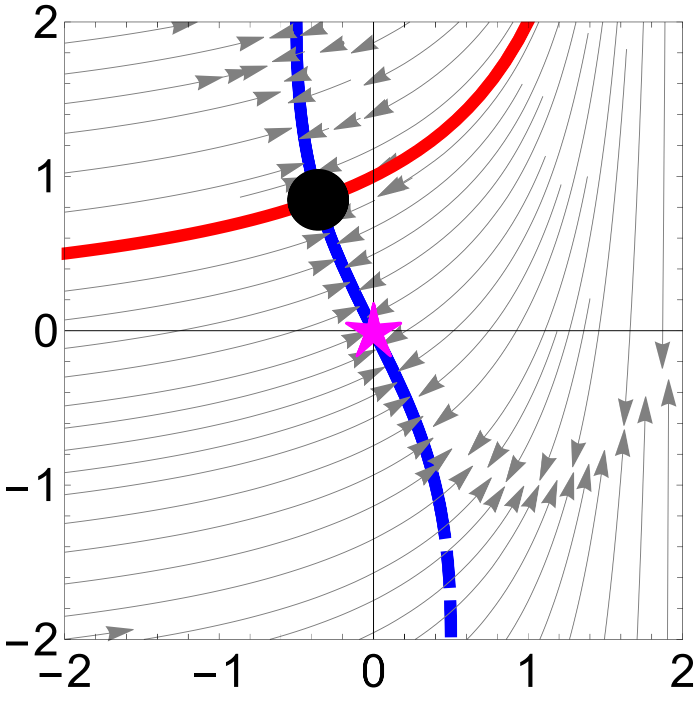


#### Fig. 5a,b. Social development, social developmental bias from partners' controls, no niche construction, and no exogenous environmental change

In[138]:=
```
Clear["Global`*"]

(*Enter the case for developmental map used (see code for Fig. 2)*)
Case = 2;

(*Enter fitness and developmental map*)
```

```
w[x_, y_] := Exp[-(x^2 + y^2)]
g[y_, yb_] := 1 + y - 3/2 y^2 - 4/5 yb(*1-y^2-yb*)

(*Enter mutational covariance matrix.
  Calculate selection gradient, total selection gradient,
and total selection gradient of controls.*)
Hy = {{0.01}};
partialwpartialz[x_, y_] = {{D[w[x, y], x]}, {D[w[x, y], y]}};
dwdz[x_, y_] =
  {{D[w[x, y], x]}, {D[w[g[y, yb], y], y]}} /. {yb → y} /. {g[y, y] → x};
dwdy[x_, y_] = D[w[g[y, yb], y], y] /. {yb → y} /. {g[y, y] → x};

(*Calculate phenotypic effects of controls.
  Calculate stabilized phenotypic effects of controls.
  Calculate L matrix*)
dzdy[y_] = {{D[g[y, yb], y], 1}} /. {yb → y};
szsy[y_] = {{D[g[y, yb], y] + D[g[y, yb], yb], 1}} /. {yb → y};
Lz[y_] = Transpose[szsy[y]].Hy.dzdy[y];

(*Plot L matrix*)
Plot[{Lz[y][[1, 1]], Lz[y][[1, 2]], Lz[y][[2, 1]], Lz[y][[2, 2]]}, {y, -2, 2},
 AxesStyle → Large, PlotStyle → {{Lighter[Gray], Thickness[0.05]},
   {Cyan, Dashing[.2], Thickness[0.04]}, {Orange, Dashing[.2], Thickness[0.04]},
   {Black, Dashing[.1], Thickness[0.02]}},
 PlotRange → {-.05, .1}, AspectRatio → 1]
Export[StringJoin[ToString[NotebookDirectory[]],
   StringJoin["FigEx.3.Social.Degen.3.L.", ToString[Case]], ".pdf"], %];

(*Enter values for plot sizes*)
zLim = 2; (*Limits for the axes*)
gThickness = 0.02; (*Line thickness for g*)
eqThickness = 0.02; (*Line thickness for evolutionary equilibria*)
eqRadius = 0.2; (*Radius for disks indicating equilibria*)

(*The following points are the stable and unstable
  admissible equilibria which are found by solving for y in dwdy=0,
substituing this in g(y), and visually picking from the
 stream plot those points that are stable or unstable*)
StablePoints[case_] := {Disk[{Root[10 - 18 * #1 - 21 * #1^2 + 45 * #1^3 & , 1, 0],
    g[Root[10 - 18 * #1 - 21 * #1^2 + 45 * #1^3 & , 1, 0],
     Root[10 - 18 * #1 - 21 * #1^2 + 45 * #1^3 & , 1, 0]]}, eqRadius]}
UnstablePoints[case_] := Disk[{0, 1}, eqRadius]

(*Color for stable and unstable equililbria*)
StableColor = {Blue};
UnstableColor = {Dashed, Blue};
```

```
(*The following finds the maximum real part of the eigenvalues
 of the jacobian matrix of evolutionary dynamic system to identify
 stable and unstable equilibria: if λmax>0 at an equilibrium point,
the equililbrium is unstable;
if λmax<0 at an equilibrium point. To avoid numerical artifacts,
we use λmax>0.01 and λmax<0.01 to identify unstable and stable equilibria.*)
λmax[x_, y_] =
  Max[Re[Eigenvalues[Simplify[D[Flatten[Reverse[Lz[Y].partialwpartialz[X, Y]]],
        {{Y, X}}]]]]]] /. {X → x, Y → y};

(*Make stream plot*)
(*First, make a plot of evolutionary equilibria*)
dwdyPlot[case_] :=
 Show[Plot[y/(-1 + 3 y), {y, -zLim, zLim}, Frame → True, FrameStyle → Large,
   AspectRatio → 1, PlotRange → {{-zLim, zLim}, {-zLim, zLim}},
   PlotStyle → {Blue, Thickness[gThickness]},
   RegionFunction → Function[{y}, λmax[y/(-1 + 3 y), y] < 0.0001], Epilog →
    {StablePoints[case], {Magenta, Text[Style["*", 40], #] & /@ {{0, 0}}}}],
  Plot[y/(-1 + 3 y), {y, -zLim, zLim}, Frame → True, FrameStyle → Large,
   AspectRatio → 1, PlotRange → {{-zLim, zLim}, {-zLim, zLim}},
   PlotStyle → {Blue, Dashing[.04], Thickness[gThickness]},
   RegionFunction → Function[{y}, λmax[y/(-1 + 3 y), y] > 0.0001]]]
(*Then, superimpose the stream plot and the admissible path*)
Show[dwdyPlot[Case],
 StreamPlot[Flatten[Reverse[Lz[y].partialwpartialz[x, y]]], {y, -zLim, zLim},
  {x, -zLim, zLim}, StreamScale → {Full, 0.1, 0.03}, StreamStyle → Gray],
 Plot[g[y, y], {y, -zLim, zLim}, PlotRange → {{-zLim, zLim}, {-zLim, zLim}},
  PlotStyle → {Red, Thickness[gThickness]}]]
Export[StringJoin[ToString[NotebookDirectory[]],
   StringJoin["FigEx.3.Social.Degen.4.dzdt.", ToString[Case]], ".pdf"], %];
```

Out[149]=

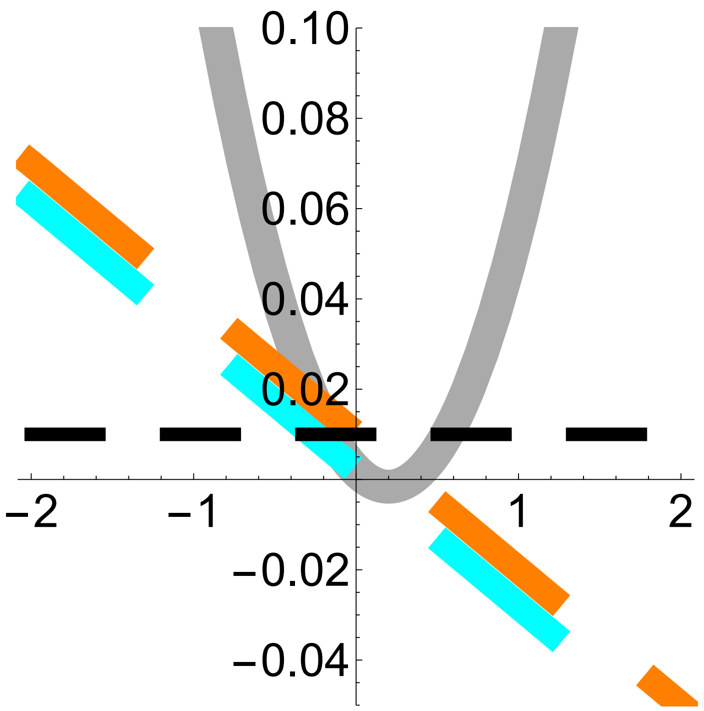


Out[161]=

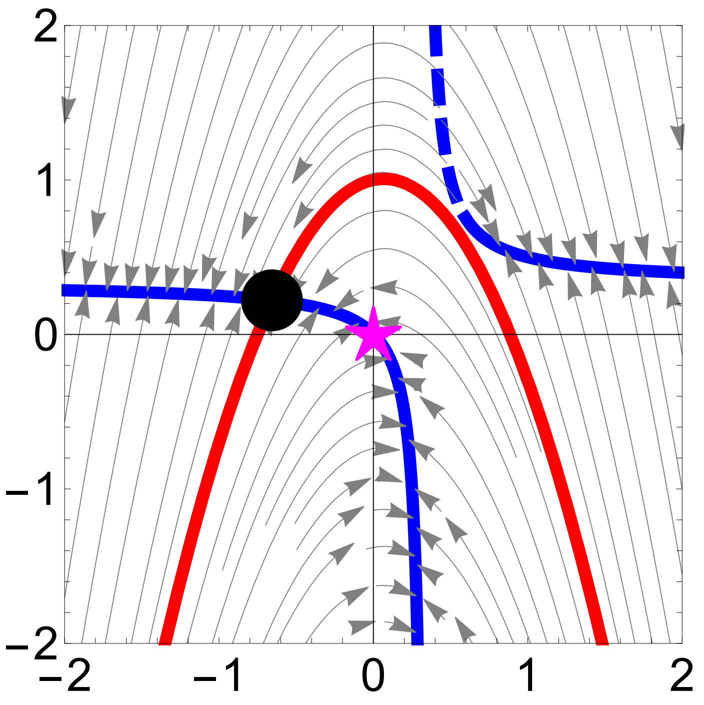


#### Fig. 5c,d. Social development, social developmental bias from partners' states, no niche construction, and no exogenous environmental change

In[163]:=
```
Clear["Global`*"]

(*Enter the case for developmental map used (see code for Fig. 2)*)
Case = 2;

(*Enter fitness and developmental map*)
```

```
w[x_, y_] := Exp[-(x^2 + y^2)]
g[y_, xb_] := 1 + y - 3/2 y^2 - 4/5 xb(*1-y^2-xb*)

(*The following is the socio-
  devo equilibrium found by solving for x in x=g[y,x]*)
xs[y_] := 5/9 (1 + y - 3/2 y^2) (*(1-y^2)/2*)

(*Enter mutational covariance matrix.
  Calculate selection gradient, total selection gradient,
and total selection gradient of controls.*)
Hy = {{0.01}};
partialwpartialz[x_, y_] = {{D[w[x, y], x]}, {D[w[x, y], y]}};
dwdy[x_, y_] = D[w[g[y, xb], y], y] /. {xb → x} /. {g[y, x] → x};

(*Calculate phenotypic effects of controls.
  Calculate stabilized phenotypic effects of controls.
  Calculate L matrix*)
dzdy[y_] = {{D[g[y, xb], y], 1}} /. {xb → xs[y]};
szsy[y_] = {{(1 - D[g[y, xb], xb])^-1 D[g[y, xb], y], 1}} /. {xb → xs[y]};
Lz[y_] = Transpose[szsy[y]].Hy.dzdy[y];

(*Plot L matrix*)
Plot[{Lz[y][[1, 1]], Lz[y][[1, 2]], Lz[y][[2, 1]], Lz[y][[2, 2]]}, {y, -2, 2},
 AxesStyle → Large, PlotStyle → {{Lighter[Gray], Thickness[0.05]},
   {Cyan, Dashing[.2], Thickness[0.04]}, {Orange, Dashing[.2], Thickness[0.04]},
   {Black, Dashing[.1], Thickness[0.02]}},
 PlotRange → {-.05, .1}, AspectRatio → 1]
Export[StringJoin[ToString[NotebookDirectory[]],
   StringJoin["FigEx.3d.Social.Degen.3.L.", ToString[Case]], ".pdf"], %];

(*Enter values for plot sizes*)
zLim = 2; (*Limits for the axes*)
gThickness = 0.02; (*Line thickness for g*)
eqThickness = 0.02; (*Line thickness for evolutionary equilibria*)
eqRadius = 0.2; (*Radius for disks indicating equilibria*)

(*The following point is the stable admissible equilibria which
  is found by solving for y in dwdy=0, substituing this in g(y,xs),
and visually picking from the stream plot those points that are stable*)
StablePoints[case_] := {Disk[{Root[10 - 2 * #1 - 45 * #1^2 + 45 * #1^3 & , 1, 0],
    g[Root[10 - 2 * #1 - 45 * #1^2 + 45 * #1^3 & , 1, 0],
     xs[Root[10 - 2 * #1 - 45 * #1^2 + 45 * #1^3 & , 1, 0]]]}, eqRadius]}

(*Color for stable and unstable equililbria*)
StableColor = {Blue};
```

```
UnstableColor = {Dashed, Blue};

(*The following finds the maximum real part of the eigenvalues
 of the jacobian matrix of evolutionary dynamic system to identify
 stable and unstable equilibria: if λmax>0 at an equilibrium point,
the equililbrium is unstable;
if λmax<0 at an equilibrium point. To avoid numerical artifacts,
we use λmax>0.01 and λmax<0.01 to identify unstable and stable equilibria.*)
λmax[x_, y_] =
  Max[Re[Eigenvalues[Simplify[D[Flatten[Reverse[Lz[Y].partialwpartialz[X, Y]]],
        {{Y, X}}]]]]]] /. {X → x, Y → y};

(*Make stream plot*)
(*First, make a plot of evolutionary equilibria*)
dwdyPlot[case_] :=
 Show[Plot[y/(-1 + 3 y), {y, -zLim, zLim}, Frame → True, FrameStyle → Large,
   AspectRatio → 1, PlotRange → {{-zLim, zLim}, {-zLim, zLim}},
   PlotStyle → {Blue, Thickness[gThickness]},
   RegionFunction → Function[{y}, λmax[y/(-1 + 3 y), y] < 0.0001], Epilog →
    {StablePoints[case], {Magenta, Text[Style["★", 40], #] & /@ {{0, 0}}}}],
  Plot[y/(-1 + 3 y), {y, -zLim, zLim}, Frame → True, FrameStyle → Large,
   AspectRatio → 1, PlotRange → {{-zLim, zLim}, {-zLim, zLim}},
   PlotStyle → {Blue, Dashing[.04], Thickness[gThickness]},
   RegionFunction → Function[{y}, λmax[y/(-1 + 3 y), y] > 0.0001]]]
(*Then, superimpose the stream plot and the admissible path*)
Show[dwdyPlot[Case],
 StreamPlot[Flatten[Reverse[Lz[y].partialwpartialz[x, y]]], {y, -zLim, zLim},
  {x, -zLim, zLim}, StreamScale → {Full, 0.1, 0.03}, StreamStyle → Gray],
 Plot[g[y, xs[y]], {y, -zLim, zLim}, PlotRange → {{-zLim, zLim}, {-zLim, zLim}},
  PlotStyle → {Red, Thickness[gThickness]}]]
Export[StringJoin[ToString[NotebookDirectory[]],
   StringJoin["FigEx.3c.Social.Degen.4.dzdt.", ToString[Case]], ".pdf"], %];
```

Out[174]=

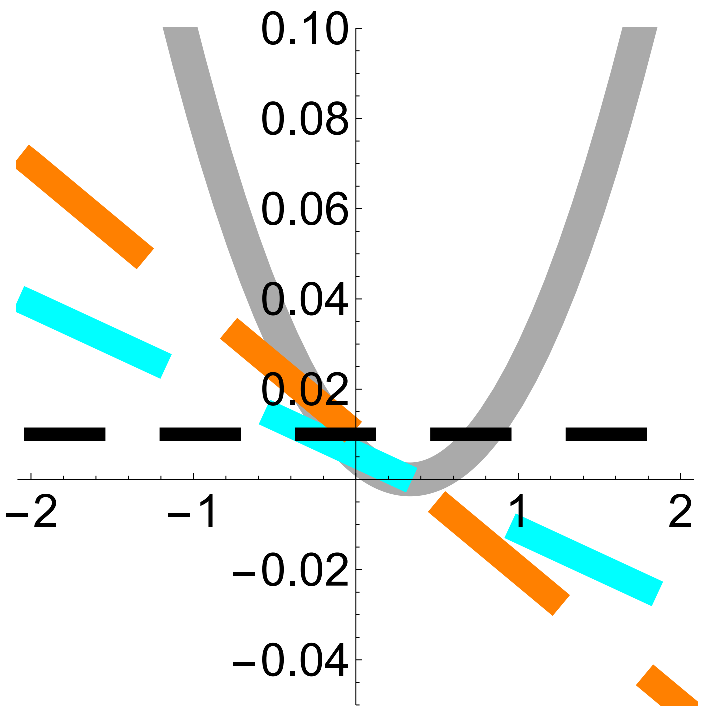


Out[185]=

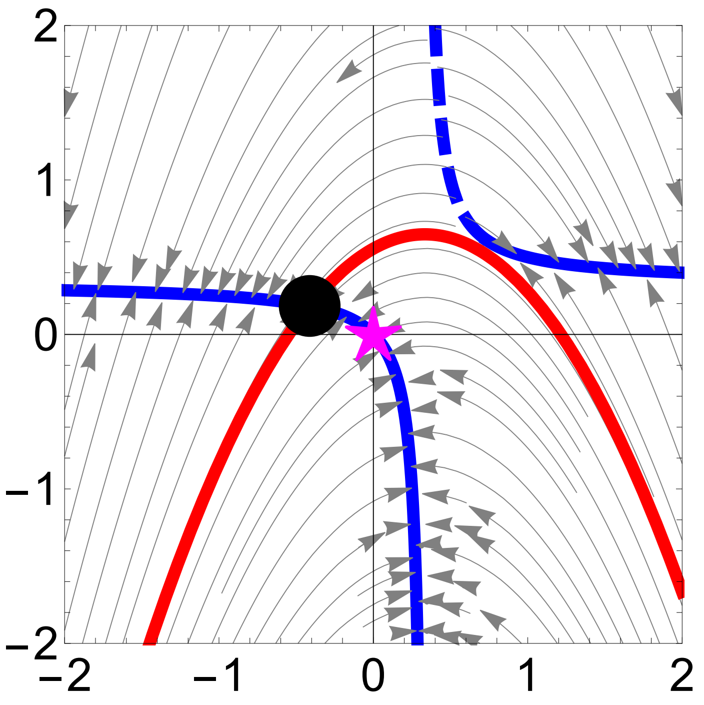


#### Fig. 6. No social development, no niche construction, but with exogenous environmental change and adaptive plasticity

##### Fig. 4a-l

In[187]:=
```
Clear["Global`*"]

(*Enter the case for developmental map used (see code for Fig. 2)*)
Case = 2;
```

```
(*Enter fitness*)
w[x_, y_] := Exp[- (x^2 + y^2)]

(*Enter selection gradient at optimal state,
to be used in developmental map*)
dwy0[y_] = D[w[0, y], y];
(*Enter developmental map*)
g[y_, ϵ_] := 1 - y^2 - ϵ dwy0[y]

(*Plot fitness landscape and admissible path at three environmental values*)
Table[
 Export[StringJoin[ToString[NotebookDirectory[]],
    StringJoin["Fig.4.Plastic.Degen.1.FitnessLandscape.Direct.",
     ToString[Case], "e.", ToString[ϵ]], ".pdf"],
   Show[Plot3D[w[x, y], {x, -2, 2}, {y, -2, 2}, AxesLabel → None,
     AxesStyle → Large, PlotStyle → Yellow],
    ParametricPlot3D[{g[y, ϵ], y, w[g[y, ϵ], y]}, {y, -2, 2},
     PlotStyle → {Darker[Brown], Thickness[0.02]}]]];, {ϵ, {0, 1, 10}}]
(*The following allows visualization of how the admissible
 path evolves as the environment evolves*)
Manipulate[Show[Plot3D[w[x, y], {x, -2, 2}, {y, -2, 2},
   AxesLabel → None, AxesStyle → Large, PlotStyle → Yellow],
  ParametricPlot3D[{g[y, ϵ], y, w[g[y, ϵ], y]}, {y, -2, 2},
   PlotStyle → {Darker[Brown], Thickness[0.02]}]], {ϵ, 0, 100}]

(*Plot total fitness landscape of controls at three environmental values*)
Table[
 Export[StringJoin[ToString[NotebookDirectory[]],
    StringJoin["Fig.4.Plastic.Degen.2.FitnessLandscape.Total.",
     ToString[Case], "e.", ToString[ϵ]], ".pdf"],
   Plot[w[g[y, ϵ], y], {y, -2, 2}, PlotRange → {0, 1}, PlotStyle →
     {Darker[Brown], Thickness[0.02]}, AxesStyle → Large]];, {ϵ, {0, 1, 10}}]
(*The following allows visualization of how total fitness landscape
 of controls evolves as the environment evolves*)
Manipulate[Plot[w[g[y, ϵ], y], {y, -2, 2}, PlotRange → {0, 1},
  PlotStyle → {Darker[Brown], Thickness[0.02]}, AxesStyle → Large], {ϵ, 0, 100}]

(*Enter mutational covariance matrix.
  Calculate selection gradient, total selection gradient,
and total selection gradient of controls.
  Calculate phenotypic effects of controls and H matrix*)
Hy = {{0.01}};
partialwpartialz[x_, y_] = {{D[w[x, y], x]}, {D[w[x, y], y]}};
dwdz[x_, y_] = {{D[w[x, y], x]}, {D[w[g[y, ϵ], y], y]}} /. {g[y, ϵ] → x};
dwdy[x_, y_] = D[w[g[y, ϵ], y], y] /. {g[y, ϵ] → x};
dzdy[y_, ϵ_] = {{D[g[y, ϵ], y], 1}};
```

```
Hz[y_, ϵ_] := Transpose[dzdy[y, ϵ]].Hy.dzdy[y, ϵ];

(*Plot H matrix at three environmental values*)
Table[
 Export[StringJoin[ToString[NotebookDirectory[]], StringJoin[
     "Fig.4.Plastic.Degen.3.H.", ToString[Case], "e.", ToString[ϵ]], ".pdf"],
   Plot[{Hz[y, ϵ][[1, 1]], Hz[y, ϵ][[1, 2]], Hz[y, ϵ][[2, 2]]}, {y, -2, 2},
    AxesStyle → Large, PlotStyle → {{Lighter[Gray], Thickness[0.05]}, {Cyan,
       Dashing[.2], Thickness[0.04]}, {Black, Dashing[.1], Thickness[0.02]}},
    PlotRange → {-0.05, 0.1}, AspectRatio → 1]];, {ϵ, {0, 1, 10}}]
(*The following allows visualization of how the H matrix
 evolves as the environment evolves*)
Manipulate[Plot[{Hz[y, ϵ][[1, 1]], Hz[y, ϵ][[1, 2]], Hz[y, ϵ][[2, 2]]},
  {y, -2, 2}, AxesStyle → Large, PlotStyle → {{Lighter[Gray], Thickness[0.05]},
    {Cyan, Dashing[.2], Thickness[0.04]}, {Black, Dashing[.1], Thickness[0.02]}},
  PlotRange → {-0.05, 0.1}, AspectRatio → 1], {ϵ, 0, 100}]

(*Enter values for plot sizes*)
zLim = 2; (*Limits for the axes*)
gThickness = 0.02; (*Line thickness for g*)

(*Make stream plot superimposed with the
 admissible path at various environmental values*)
Table[
 Export[StringJoin[ToString[NotebookDirectory[]],
    StringJoin["Fig.4.Plastic.Degen.4.dzdt.", ToString[Case], "e.",
     ToString[ϵ]], ".pdf"], Show[(*dwdyPlot[Case],*)StreamPlot[
     Flatten[Reverse[Hz[y, ϵ].partialwpartialz[x, y]]], {y, -zLim, zLim},
     {x, -zLim, zLim}, StreamScale → {Full, 0.1, 0.03}, StreamStyle → Gray],
    Plot[g[y, ϵ], {y, -zLim, zLim}, PlotRange → {{-zLim, zLim}, {-zLim, zLim}},
     PlotStyle → {Red, Thickness[gThickness]}]]]];, {ϵ, {0, 1, 10}}]
(*The following allows visualization of how the stream plot evolves
 as the environment evolves*)Manipulate[Show[(*dwdyPlot[Case],*)
  StreamPlot[Flatten[Reverse[Hz[y, ϵ].partialwpartialz[x, y]]], {y, -zLim, zLim},
   {x, -zLim, zLim}, StreamScale → {Full, 0.1, 0.03}, StreamStyle → Gray],
  Plot[g[y, ϵ], {y, -zLim, zLim}, PlotRange → {{-zLim, zLim}, {-zLim, zLim}},
   PlotStyle → {Red, Thickness[gThickness]}]], {ϵ, 0, 100}]


Out[192]= {Null, Null, Null}
```

Out[193]=

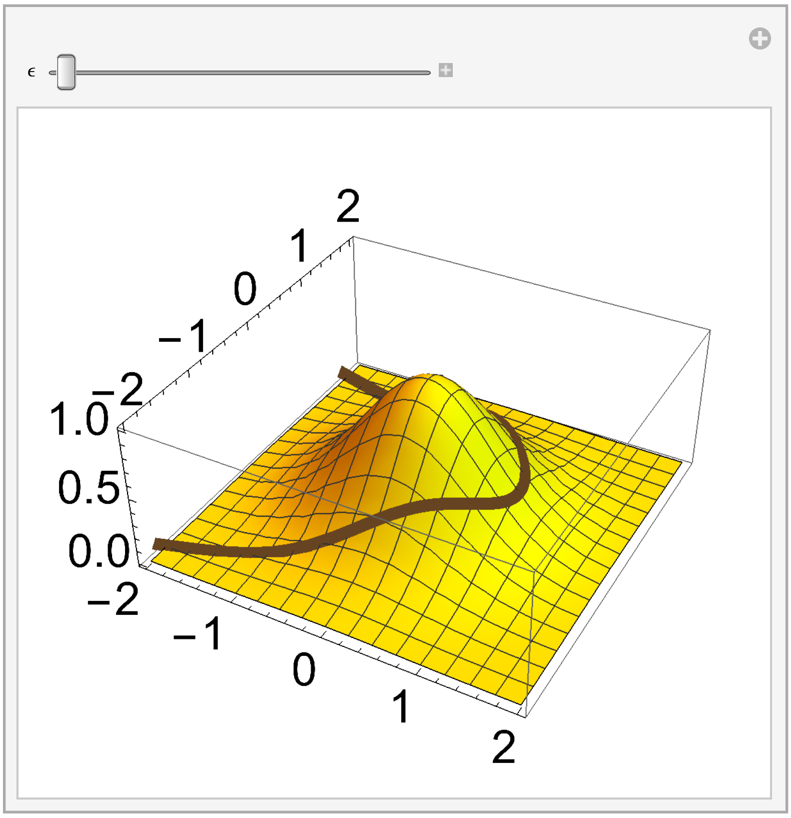


Out[194]= {Null, Null, Null}

Out[195]=

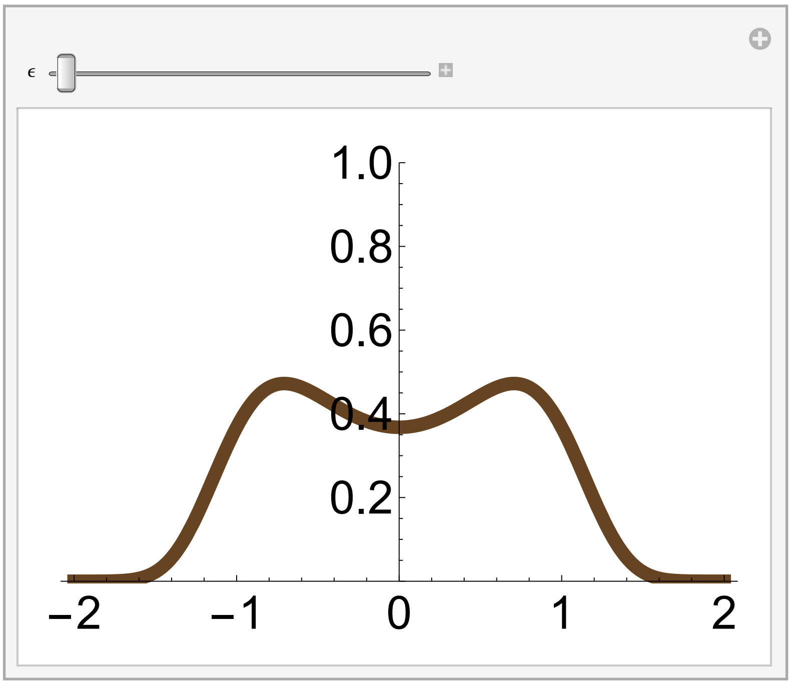


Out[202]= {Null, Null, Null}

Out[203]=

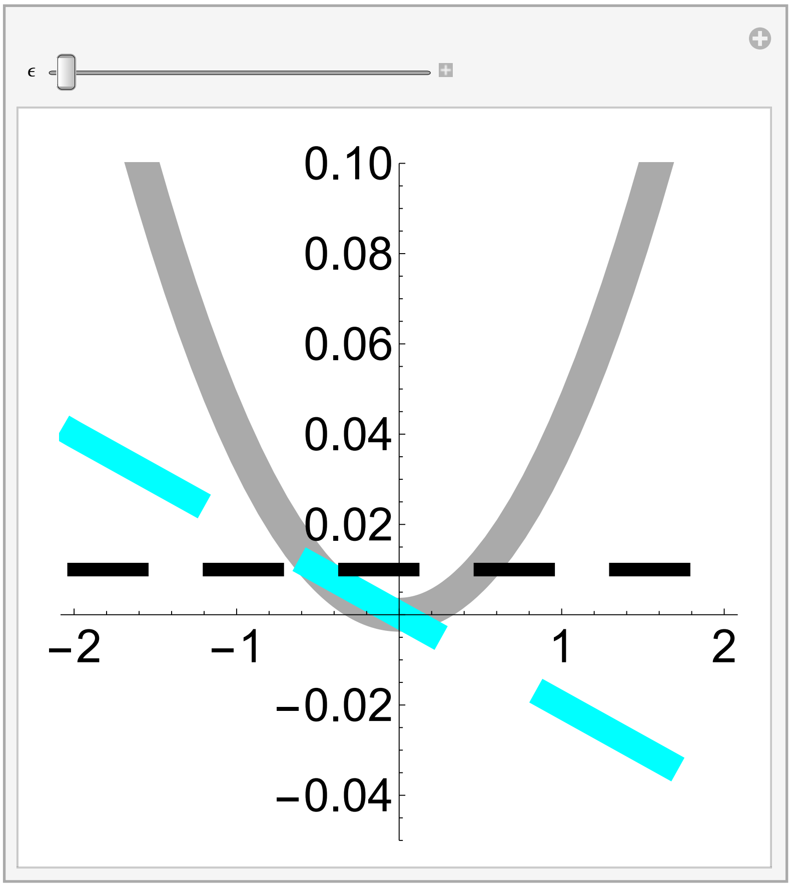


Out[206]= {Null, Null, Null}

Out[207]=

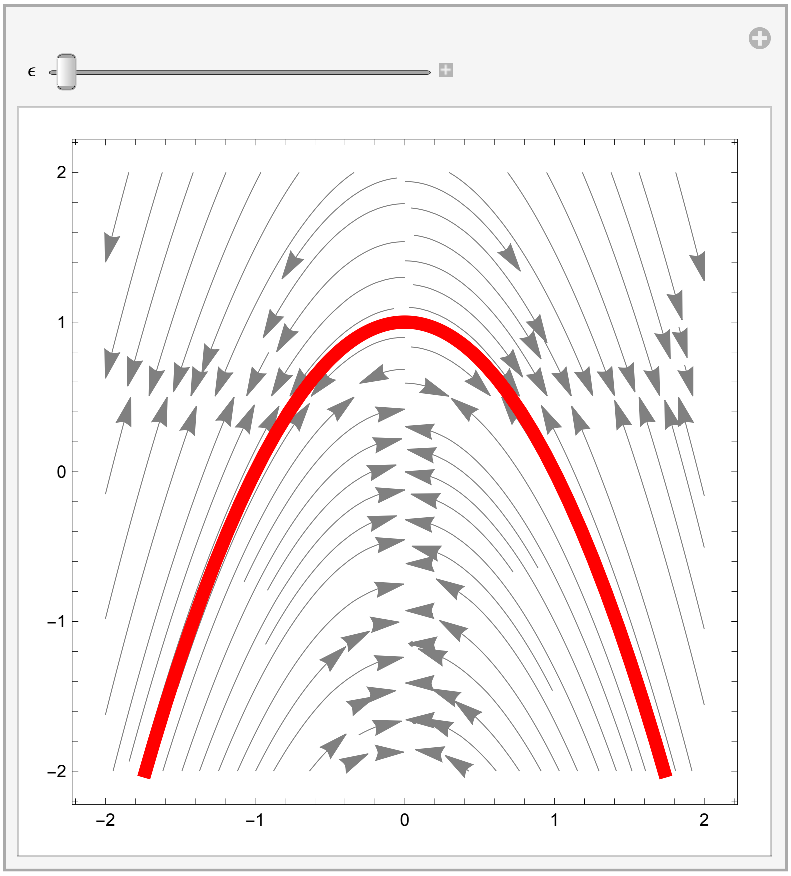

#### Fig. 6m

In[208]:=
```
(*Make plot of evolutionary dynamics*)

(*Enter environmental map*)
h[τ_] = 1/100 τ;

(*Enter plasticity matrix*)
dxdϵ[y_, τ_] = D[g[y, ϵ], ϵ] × D[h[τ], τ] /. {ϵ → h[τ]};

(*Enter values for plot sizes*)
zLim = 2;

(*Enter range of values for the initial condition of y*)
yLimSol[case_] := {-1.5, 1.5}
(*Enter the step size to vary the initial conditon of y*)
yStepSol[case_] := .5
(*The initial conditions for x(τ) and y(τ) at τ=0 are g(y(0),h(0)) and y(0),
respectively. *)

Show[Table[
  (*Enter stopping time*)
  maxT = 10 000 ;
  (*Solve dynamic equations*)
  sol = NDSolve[{X'[τ] ==
       (Hz[Y[τ], h[τ]].partialwpartialz[X[τ], Y[τ]])[[1, 1]] + dxdϵ[Y[τ], τ],
     Y'[τ] == (Hz[Y[τ], h[τ]].partialwpartialz[X[τ], Y[τ]])[[2, 1]],
     X[0] == g[Y0, h[0]], Y[0] == Y0}, {X[τ], Y[τ]}, {τ, 0, maxT}];
  (*Obtain solution*)
  Xsol[T_] = Evaluate[sol[[1, 1, 2]]] /. {τ → T};
  Ysol[T_] = Evaluate[sol[[1, 2, 2]]] /. {τ → T};
  SelResp[Y0] = Transpose[partialwpartialz[Xsol[τ], Ysol[τ]]].
    Hz[Ysol[τ], h[τ]].partialwpartialz[Xsol[τ], Ysol[τ]];
  PlasResp[Y0] = Transpose[partialwpartialz[Xsol[τ], Ysol[τ]]].
    {{dxdϵ[Ysol[τ], τ]}, {0}};
  (*Plot solution*)
  Show[ParametricPlot[{Ysol[τ], Xsol[τ]}, {τ, 0, maxT}, PlotStyle → Black,
    Frame → True, FrameStyle → Large, PlotRange → {{-zLim, zLim}, {-zLim, zLim}},
    Epilog → {Magenta, Text[Style["★", 40], #] & /@ {{0, 0}}}],
   ListPlot[{{Ysol[0], Xsol[0]}}, PlotMarkers → {"□", 40}, PlotStyle → Black],
   ListPlot[{{Ysol[maxT], Xsol[maxT]}},
    PlotMarkers → {"■", 40}, PlotStyle → Black]],
  {Y0, yLimSol[Case][[1]], yLimSol[Case][[2]], yStepSol[Case]}]]
Export[StringJoin[ToString[NotebookDirectory[]],
   StringJoin["Fig.4.Plastic.Degen.5.dzdt.", ToString[Case]], ".pdf"], %];
```

Out[213]=

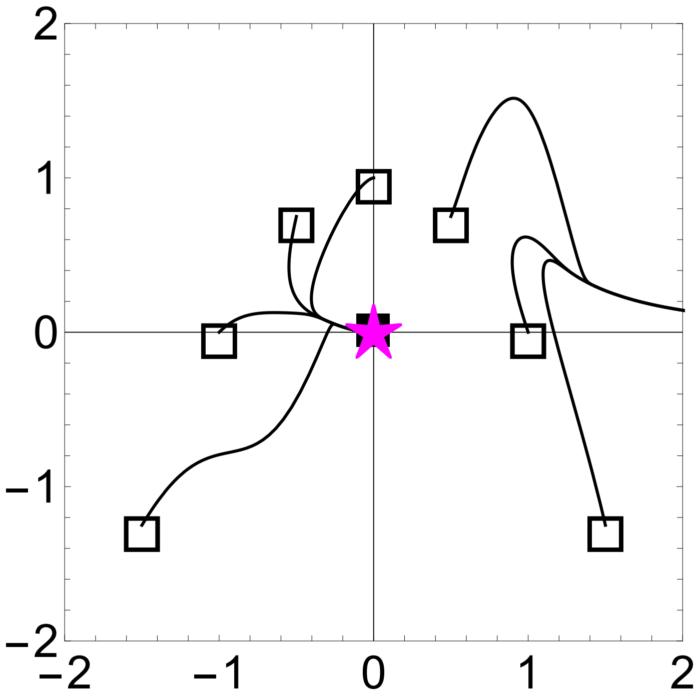
